# Salmonids reveal principles of regulatory evolution following autotetraploidization

**DOI:** 10.64898/2025.12.18.695160

**Authors:** Marie-Odile Baudement, Diego Perojil Morata, Gareth B. Gillard, Pooran S. Dewari, Manu Kumar Gundappa, Tomasz Podgorniak, Lars Grønvold, Damir Baranasic, Audrey Laurent, François Giudicelli, Bojan Zunar, Erika Carrera-García, Aline Perquis, Aurélien Brionne, Tan Thi Nguyen, Rose Ruiz Daniels, Gabriela A. Merino, David Thybert, Garth R. Ilsley, Alexandra Louis, Torgeir R. Hvidsten, Camille Berthelot, Peter W. Harrison, Hugues Roest Crollius, Yann Guiguen, Boris Lenhard, Simen R. Sandve, Julien Bobe, Matthew P. Kent, Sigbjørn Lien, Daniel J. Macqueen

## Abstract

Early vertebrate autotetraploidization events may have enabled major innovations by expanding the genetic material for functional diversification, yet their ancient timing obscures how genome doubling reshaped gene regulatory evolution. Salmonids provide a unique window to these mechanisms, because they experienced a comparatively recent autotetraploidization and are earlier in the rediploidization process - which creates new genes and regulatory elements during evolution. Here, using large-scale multiomics spanning embryonic and adult tissues in two salmonids, we investigate gene regulatory evolution following genome doubling and rediploidization, which we show is governed by developmental and tissue-specific context, with a period of maximal constraint at advanced stages of embryogenesis. This work advances understanding of vertebrate genome evolution, while providing an open resource supporting salmonid aquaculture and conservation.

## Main Text

Whole genome duplication (WGD, i.e. polyploidy) occurred at many stages of eukaryotic evolution (*1–3*), including three sequential rounds in the common ancestors to all vertebrates (1R), jawed vertebrates (2R) (*4–7*) and teleost fish (3R) (*8*, *9*). While many duplicated genes created by WGD (ohnologs) (*1*) are lost, retained ohnologs are enriched in functions linked to development, signaling and transcriptional regulation (*2*, *10*). Ohnologs retained from vertebrate WGDs have gained more complex regulation and specialized in expression during evolution (*11*, *12*). Events in the common ancestors to all vertebrates (∼530 Mya, 1R) (*5–7*, *13*) and teleosts (∼275 Mya, 3R) (*14*) are proposed autotetraploidizations (*5–7*, *15*), where a diploid genome doubled within-species, leading to four identical chromosome sets that paired non-preferentially during meiosis, resulting in tetrasomic inheritance (*1*). A separate WGD (2R) in the jawed vertebrate ancestor may have occurred by allotetraploidization (*5–7*, *13*), where genetically distinct species hybridized, leading to two subgenomes within the same nucleus (*1*).

Rediploidization involves a return to stable bivalent pairing, resulting in the disomic inheritance of unlinked ohnolog loci that can diverge in sequence and hence function (*1*, *16*, *17*). While this may be immediate in allopolyploids with distinct subgenomes, for autopolyploids, it requires the evolution of genomic reorganizations (*18*). Understanding the functional outcomes of rediploidization is necessary to unravel the genomic basis for major vertebrate radiations. However, studies of regulatory changes following the 1R-3R WGDs are challenged by the vast evolutionary time separating these events from living species.

The salmonid ancestor underwent autotetraploidization 80-100 Mya, representing a fourth round (4R) of WGD (*19–21*). Around half of all salmonid genes are retained as functional ohnolog pairs from this event (*18*, *22*), three-fold more than the 1R-3R events (*23*). Salmonid ohnologs are located in large collinear regions that have diverged in sequence to a highly variable extent (*18*, *20*). This signature results from asynchronous rediploidization (*17*, *20*), which likewise followed WGDs at the stem of teleosts (3R) (*15*), acipenseriforms (*24*) and potentially all vertebrates (1R) (*6*). Asynchronous rediploidization creates a stratification of ohnolog divergence times within affected genomes, providing an ideal study system to explore the function and regulation of ohnologs at distinct stages of evolution. A large fraction of ohnolog pairs created by the salmonid 4R have evolved divergent expression (*18*, *25*). However, the mechanistic underpinnings of ohnolog regulatory evolution remains poorly understood in terms of promoter and enhancer activity.

To address this gap, we generated large-scale multiomics datasets for two salmonids, which were used to investigate changes in gene expression and regulatory element activity following the 4R autotetraploidization, revealing the role of ontogeny, tissue-specific context and rediploidization in shaping genome functional evolution.

### Functional and comparative annotation of salmonid genomes

Following ENCODE (*26*) and within the Functional Annotation of Animal Genomes (FAANG) initiative (*27*, *28*), AQUA-FAANG aimed to deliver comprehensive genome functional annotations for farmed fishes of global economic importance (*29*). To this end, we obtained species-matched samples from Atlantic salmon and rainbow trout (hereafter: salmon and trout), among the world’s most commercially-valuable fishes (*30*), with wild populations of substantial cultural and conservation importance (*31*). Sampling spanned key stages of embryogenesis (‘DevMap’) and a tissue panel (for both sexes) at sexually immature and mature stages (‘BodyMap’) (Fig. 1A). After developing robust experimental protocols (Fig. S1; Text S1-S4), RNA-seq was used to quantify gene expression, while ATAC-seq (*32*) and ChIP-seq (for histone marks: H3K27ac, H3K4me1, H3K4me3 and H3K27me3) was used to assess chromatin openness and epigenetic state (Fig. 1B). Altogether, we generated 771 high-quality sequencing datasets (Tables S1-S3; Fig. S2-S9) summing to 77.7 billion read pairs, with raw data shared on the <u>FAANG data portal</u> (*33*). Processed datasets are shared at salmobase.org (*34*), alongside new comparative genomic tools for data visualization and exploration (described below).

**Fig. 1.**
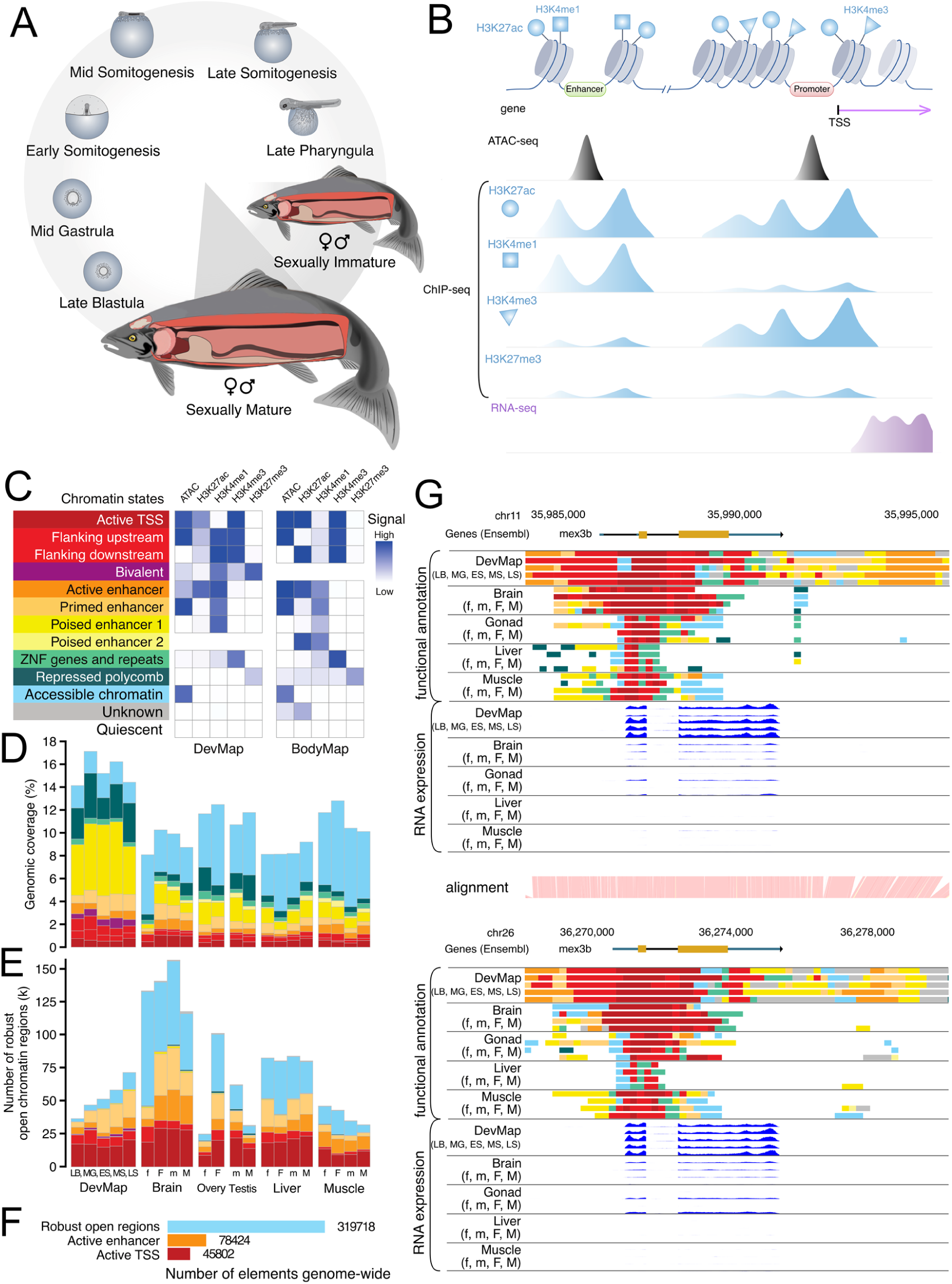
Regulatory annotation of salmonid genomes. Data shown for salmon (equivalent data for trout shown in Fig. S10). (**A**). Overview of sampling spanning embryogenesis (DevMap) and adult tissues at two stages of sexual maturation (BodyMap). (**B**) Regulatory elements were profiled using ATAC-seq and ChIP-seq and gene expression using RNA-seq. (C) 12-state ChromHMM models for DevMap and BodyMap datasets, including signal intensity for individual functional assays per chromatin state. (**D** & **E**) Coverage of the genome (**D**) and number of robust open chromatin regions (**E**) per chromatin state across samples (DevMap: LB = late blastulation, MG = mid-gastrulation; ES = early somitogenesis; MS = mid somitogenesis; LS = late somitogenesis; BodyMap: tissue samples separated into immature (lowercase ‘f’ or ‘m’) and mature (uppercase ‘F’ and ‘M’) for females and males, respectively. (**F**) Number of robust open chromatin regions the salmon genome, including those comprising active promoters and enhancers according to ChromHMM annotation. (**G**) Integration of functional annotation data for a pair of *mex3b* ohnologs (ENSSSAG0000008892 and ENSSSAG00000085168) encoding a conserved RNA-binding protein. Predicted chromatin states (colored as panel **C**) and corresponding gene expression are visualized across samples. This example illustrates conservation of regulatory elements between ohnologs that correlate with conserved gene expression. Equivalent data for rainbow trout *mex3b* ohnologs provided in Fig. S17.

We identified 319,718 robust open chromatin regions in salmon and 194,002 in trout, expected to harbour regulatory elements bound by transcription factors (TFs) and protein complexes associated with RNA polymerase II (*32*). To enrich the functional context of these regions, we modelled chromatin epigenetic state using ChromHMM (*35*), integrating our ChIP-seq datasets. This identified 45,802 and 28,138 unique active promoters, and 78,424 and 42,164 unique active enhancers, in the salmon and trout genomes, respectively (Fig. 1C-F, Fig. S10). We performed two analyses to validate these annotations. First, Cap Analysis Gene Expression (CAGE) (*36*) using DevMap samples in both species showed that most open chromatin regions annotated as ‘Active TSS’ were supported by CAGE-defined promoters, with support observed for 83–91% in salmon and 53–70% in trout (Fig. S11). Second, we found that active promoters and enhancers were enriched in vertebrate TF binding sites (TFBS), matching sample type expectations (Fig. S12-13; Text S5). For example, the TFBS targeted by the Pou5f1(Oct4):Sox2 complex, essential for zebrafish zygotic genome activation (ZGA) and gastrulation initiation (*37*), was enriched in highly active salmon and trout regulatory elements at the same developmental stages (Fig. S12). We also identified enrichment of binding sites for canonical tissue TFs in both species, with extensive between-species sharing of the same enriched TFBSs (Text S5, Fig. S13-14).

For comparative analysis, we identified 10,521 high-confidence ohnolog pairs conserved in both species and 5,832 ‘singletons’, where one ohnolog was lost in their common ancestor (Tables S4-S5; Fig. S15). To compare non-coding regions, we adapted Cactus (*38*) to align duplicated and orthologous sequences within and between the genomes of salmon and trout (Methods; Fig. S16; Table S6), alongside single orthologous regions from their sister lineage (northern pike, *Esox lucius*) that didn’t undergo the 4R WGD (*19*). Our comparative dataset captured 26,157 genes and 194,528 unique promoters or enhancers showing activity in at least one sample across both species. We provide examples of the utility of these comparative tools to infer gene regulatory dynamics among salmonid ohnolog pairs (Fig. 1G; Fig. S17).

### Dynamic genome regulation across ontogeny and tissues

We classified our DevMap and BodyMap datasets using dimensionality reduction approaches (*39*). Across embryogenesis, self-organizing maps (SOM) identified respective clusters of genes (Fig. 2A; Fig. S18-19) or genomic regions (Fig. 2B; Fig. S20-21) showing closely-related changes in expression or chromatin openness. SOM captured the maternal to zygotic transition (MZT) according to clusters of maternal mRNAs showing highest levels during late cleavage or early blastulation, with declining levels thereafter (Fig. 2A; M1-4 in salmon; M1-3 in trout), alongside genes up-regulated (Fig. 2A; B1-2 and G1; both species) and chromatin regions showing maximal openness (Fig. 2B; clusters 1-7 for salmon; 1-11 for trout) at blastulation and gastrulation stages. The maternal RNA SOM clusters showed enriched Gene Ontology (GO) terms including ‘cell cycle’ and ‘chromosome segregation (Fig. S22-23, Text S6, Table S7-S8). Transcriptomic clusters specific to segmentation stages, leading to the most developmentally advanced pharyngula stages (S1-S2 and P1-P2; both species), were enriched for terms associated with organogenesis and energy production (Fig. 2A; Fig. S22-23, Text S6, Table S7-S8). A developmental trajectory of embryogenesis was evident, with topologically-related SOM clusters co-localizing on UMAP graphs (Fig. 2A-B). Comparing the two salmonids, closely-related SOM clusters were enriched in orthologous genes and syntenic genomic regions (Fig. 2A-B, cross-species bands between clusters; Fig. S24).

**Fig. 2.**
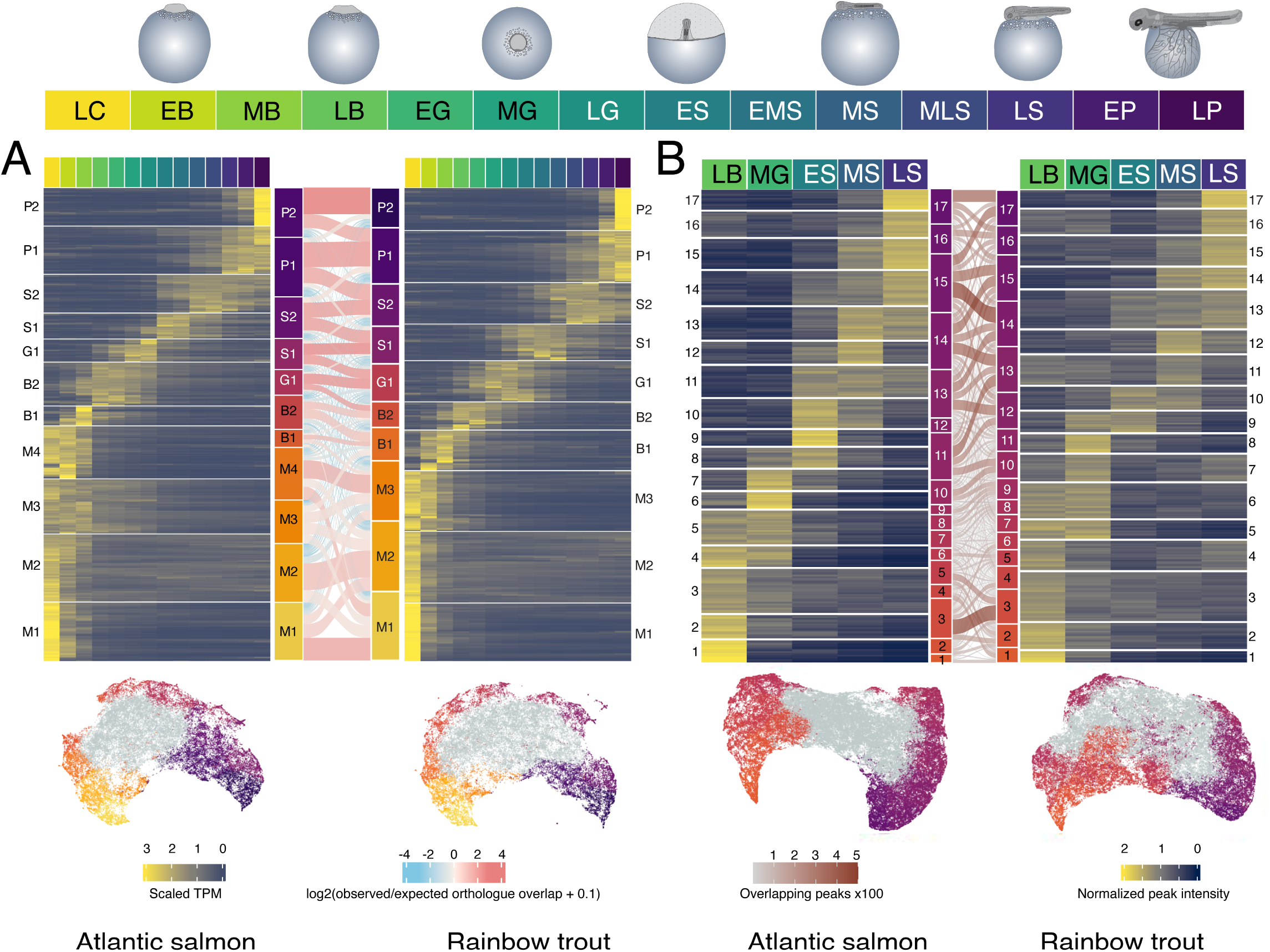
Functional activity of salmonid genomes during embryogenesis. (**A**) Transcriptome and (**B**) chromatin openness dynamics during embryogenesis. Respective data shown are SOM clusters for mean normalised transcripts per million values (TPM, RNA-Seq) and mean normalised read intensity values in robust open chromatin regions (ATAC-seq). Full SOM clusters shown in Figs S18-21. For panel A, light red bands illustrate the observed-to-expected ratio of orthologous gene expression between the two species across all SOM cluster combinations. For panel **B**, brown bands between clusters show the number of syntenic open chromatin regions based on whole genome alignment. On UMAP plots shown under each heatmap, dots represent mean normalised TPM values of individual genes (**A**) or mean normalised ATAC-seq read intensity values for individual robust open chromatin regions (**B**). Grey dots on each UMAP are genes or chromatin regions captured within constitutive SOM clusters (Figs S18-S21), respectively. Schematic at top highlights embryonic stages with colours matched to SOM clusters: LC = late cleavage; EB, MB, LB = early-, mid- and late blastulation; EG, MG, LG= early, mid- and late gastrulation; ES, EMS, MS, MLS, LS = early, early-mid, mid-, mid-to-late and late somitogenesis; EP, LP = early and late pharyngula.

These analyses were repeated using our BodyMap datasets, capturing changes in gene expression and chromatin openness across tissues, sexes, and maturation stages that were similar in both species (Fig. S25; supported by Fig. S26-32; Tables S7-S8; Text S7).

Overall, the AQUA-FAANG datasets capture dynamic changes in genome activity across life-stages and tissue types, providing a novel framework to dissect the evolution of gene regulation following the salmonid 4R autotetraploidization.

### Ohnolog regulatory divergence mirrors constraints shaping gene expression across species

Ontogeny and tissue-specific context are established to influence gene expression evolution across vertebrate species (*11*, *40–44*). Do similar constraints act on rediploidization following autotetraploidization? To address this, we quantified salmon ohnolog pair expression divergence using our transcriptomic datasets, adapting a published strategy (*11*) (Fig. 3A-B). Salmonid 4R ohnolog pairs were split into two sub-classes to explicitly account for the rediploidization process (*17*, *18*). The first consists of 8,757 ‘Early rediploidization’ pairs, where ohnolog sequence divergence started before salmon and trout separated ∼25 Mya (*19*). The second consists of 1,103 ‘Late rediploidization’ pairs, where ohnolog divergence occurred independently in salmon and trout, owing to a prolonged phase of tetrasomic inheritance (*17*, *20*).

**Fig. 3.**
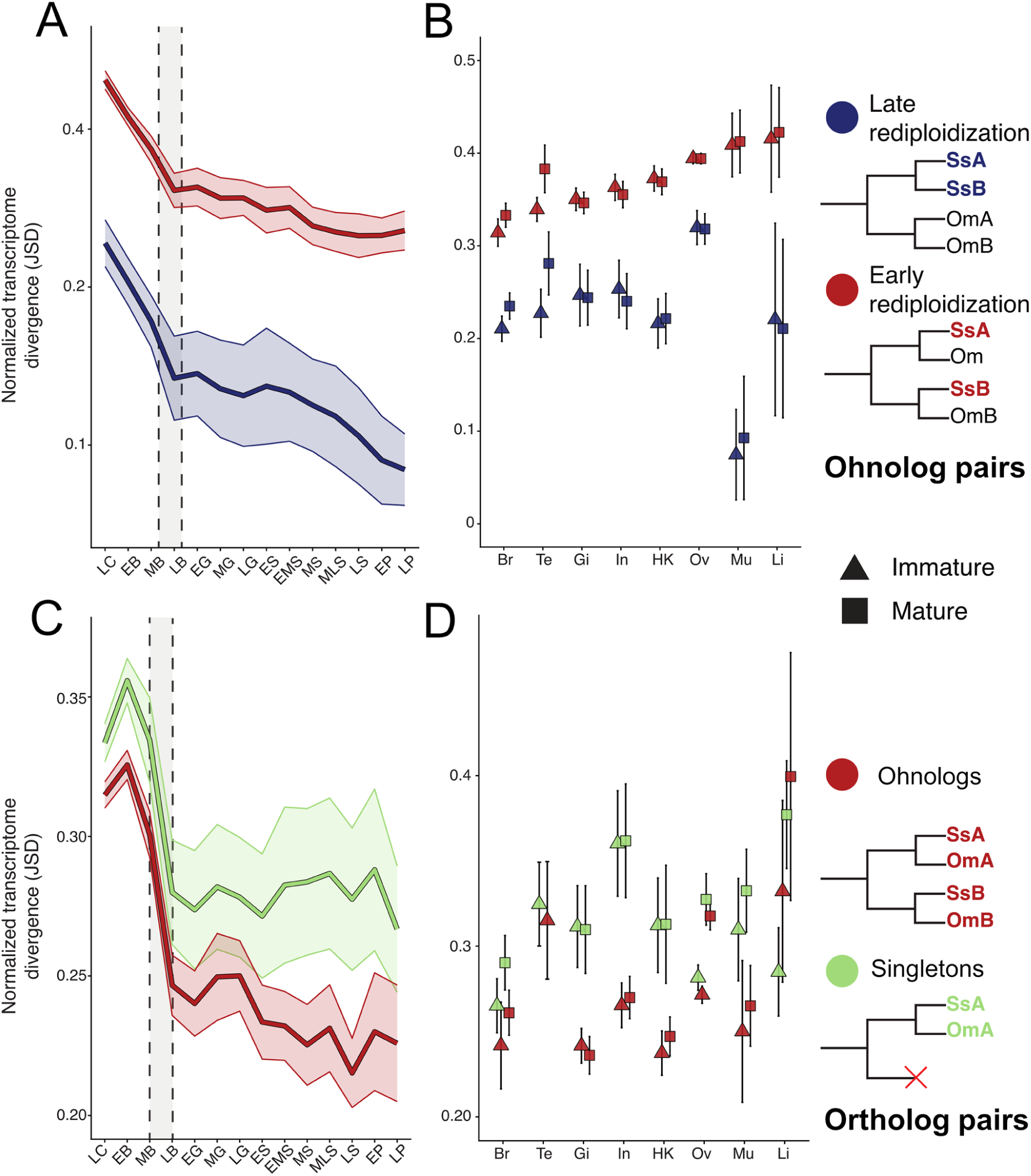
Gene regulatory evolution following the salmonid 4R autotetraploidization. (**A**-**B**) Mean normalized Jensen–Shannon divergence (JSD) ± 1 SD (100 bootstrap samples) between salmon ohnolog pairs across 14 stages of embryonic development (DevMap: A; grey area between dashed lines represents the MZT period) or adult tissues of immature and mature fish (triangles and squares, respectively) (BodyMap: B). Lines are colored by Early (red) and Late (blue) rediploidization categories. (**C**-**D**) Mean JSD ± 1 SD for ortholog pairs from salmon and trout for two groups representing ohnologs (retained as pairs in both species) or singletons (one gene lost in same ohnolog pair in the salmon-trout ancestor) for the same DevMap (**C**) and BodyMap (**D**) samples. JSD was calculated from quantile-normalised TPM values. Higher normalized mean JSD indicates greater expression divergence between gene pairs. DevMap stage abbreviations are as per the Fig. 1 legend. BodyMap tissue abbreviations: Br = brain; Te = testis; Gi= gill; In = distal intestine; HK = head kidney; O =ovary; Mu = skeletal muscle; Li = liver.

As expected, rediploidization was the dominant factor explaining regulatory divergence (*17*), with Early rediploidization ohnolog pairs showing greater divergence for all sample types (Fig. 3A-B; non-overlapping 95% CI between rediploidization sub-classes). Ohnolog expression divergence decreased across embryogenesis, being highest at the earliest cleavage stage (pre-MZT), and lowest at the most developmentally-advanced segmentation/pharyngula stages (Fig. 3A), thought to represent the vertebrate phylotypic period (*43*). The observed pattern of decreasing regulatory divergence across embryogenesis was highly correlated between Early and Late rediploidization ohnolog pairs (Pearson’s R = 0.97, *p* = 1.1 x 10^-8^, Fig. S33), albeit with a more pronounced drop in divergence for Late rediploidization ohnolog pairs at advanced segmentation stages (Fig. 3A).

Adult tissue types differed extensively in ohnolog expression divergence, with brain showing the least divergence for Early rediploidization pairs (Fig. 3B), consistent with past work (*45*). Sexual maturation had little effect on ohnolog expression divergence for most tissues, including ovaries (Fig. 3B; overlapping 95% CI). However, ohnolog expression divergence was greater in sexually mature compared to immature males, specifically in testis (Fig. 3B; non-overlapping 95% CI).

Unlike embryogenesis, expression divergence across tissues was not significantly correlated between Early and Late rediploidization ohnolog pairs (Fig. S33). This was explained by increased expression divergence in ovaries, alongside reduced divergence in muscle and liver, for the Late rediploidization ohnolog pairs.

We next explored the extent to which salmonid 4R ohnolog expression divergence mirrors across-species regulatory evolution. We repeated our expression divergence analysis, this time using 1-to-1 ortholog pairs from salmon and trout (Fig. 3C-D). To account for impacts of WGD on regulatory evolution across species, we separated these ortholog pairs into two groups. The first consists of 8,661 ortholog pairs representing Early rediploidization ohnologs (with both ohnolog copies retained in salmon and trout) and the second 5,099 ortholog pairs representing singletons in both species.

Across-species expression divergence of ortholog pairs belonging to the ohnolog group was highly correlated with that of salmon ohnolog pairs across embryonic development (Pearson’s R = 0.96, *p* = 1.0 x 10⁻⁷), and weakly correlated across adult tissue types (Pearson’s R = 0.54, *p* = 0.037) (Fig. S33). Expression divergence of ortholog pairs differed between the ohnolog and singleton groups. In both cases, expression divergence decreased across embryogenesis, with a strong drop around the MZT, matching that observed for salmon ohnolog pairs (Fig. 3A, C). However, after the MZT, ortholog pairs from the ohnolog group showed reduced expression divergence during segmentation, compared to singletons (Fig. 3C; lack of overlap between 95% CI). This result indicates that orthologs evolving under the highest constraints on expression during mid-to-late embryogenesis are overrepresented as ohnolog pairs.

Across tissues, we observed variable and often significant differences in how ortholog pairs have evolved across species, depending on whether they represent ohnologs or singletons (Fig. 3D). For gill, head kidney and distal intestine, ortholog pairs from the ohnolog group showed significantly less expression divergence than singletons (Fig. 3D). However, in brain, expression divergence of ortholog pairs was similar for both groups, which can be explained by comparatively slower evolution of singleton orthologs (Fig. 3D). Specific to ovaries, regulatory divergence of ortholog pairs across species was driven by maturation stage, rather than their retention status following WGD (Fig. 3D).

In sum, these results reveal the nature of post-WGD regulatory evolution across ontogeny and tissues, implicating pleiotropic constraints during a period of rapid organ development at mid-to-late embryogenesis (*40*) as one strong factor selecting for the long-term retention of ohnologs, while also highlighting ohnolog retention as a driver of tissue regulatory evolution both within and across species, but with highly variable effects across tissue types.

### Functional evolution of regulatory elements retained from autotetraploidization

Our DevMap and BodyMap atlases enable robust inferences on regulatory element evolution. Using our whole genome alignment, we divided all promoters and enhancers from salmon into three categories of post-WGD evolution: “Shared” - where two duplicated sequences retained from WGD are aligned and each shows activity; “Alignable only” – where, despite being aligned, activity is restricted to one of two duplicated sequences retained from WGD; and “Exclusive” – where the active element is limited to one chromosome (Fig. 4A). To explicitly capture the rediploidization process, we further separated these elements based on their locations within collinear blocks showing Early or Late rediploidization (*17*) (Fig. 4B; Tables S9-10).

**Fig. 4.**
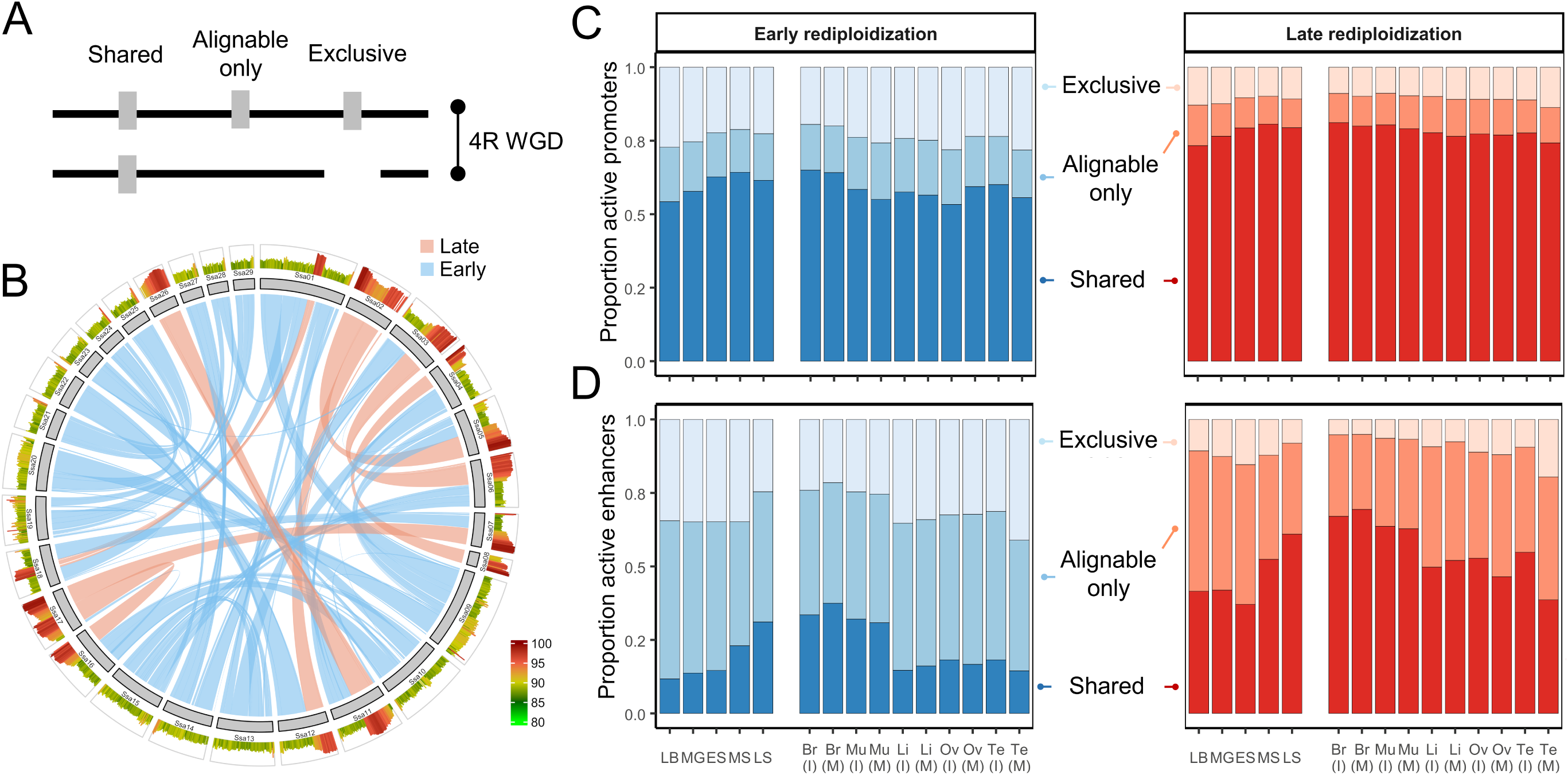
Functional retention of regulatory elements following the salmonid 4R autotetraploidization. (**A**) Schematic of Shared, Alignable only, and Exclusive regulatory elements, categorized by whole genome alignment. (**B**) Circos plot of 29 salmon chromosomes. Ribbons show collinear regions retained from the salmonid 4R WGD, with blue and red colours indicating Early and Late rediploidization regions (Tables S9-10), where duplicated regions share a respective 88.6% (SD: 1.51%) and 95.1% (SD: 2.67%) nucleotide sequence identity. The outermost panels show percentage nucleotide sequence identity (as per the provided key) between collinear blocks at 1 Mb intervals. For each alignment (Shared, Alignable only, Exclusive) and Rediploidization (early, late) category, we present: The proportion of active promoters (**C**) and enhancers (**D**) in each sample type.

Overall, 53.5%, 21.6% and 24.9% of active promoters belonged to the Shared, Alignable only, and Exclusive categories, respectively (Table S11). The proportion of Shared promoters is consistent with the estimated 55% retention rate for salmonid 4R ohnolog pairs (*18*). For active enhancers, 33.3%, 43.8% and 22.9% belonged to the same categories (Table S11). Compared to promoters, the lower proportion of enhancers from the Shared category, alongside higher proportion from the Alignable only category (both *p* < 0.0001, Fisher’s Exact Test) shows that the faster evolutionary turnover of enhancers observed between species (*46*, *47*) also occurs following WGD.

Early rediploidization regions harbored proportionately fewer Shared promoters and enhancers, alongside more from the Exclusive category (Text S8). Thus, a longer period of divergence has facilitated the loss of duplicated regulatory elements created by rediploidization, while providing more evolutionary opportunities for the gain of novel regulatory elements post-WGD.

### Ontogenetic and tissue-specific constraints on regulatory element evolution

We next explored variation in the functional retention of regulatory elements across sample types (Fig. 4C-D; supporting tests in Tables S12-S13), revealing that promoters were active in more sample types than enhancers (Fig. S34), consistent with past work (*48*). For both types of regulatory element, the breadth of activity across sample types and extent of sample type-specific activity varied across alignment categories (Fig. S35, Text S8). Mirroring ohnolog expression divergence (Fig. 3), rediploidization was the dominant factor determining the proportion of Shared promoters and enhancers across all sample types (Fig. 4C-D).

The proportion of active promoters per alignment category showed limited variation across sample types, with Shared promoters most common (Fig. 4C; Text S8). Much greater heterogeneity across sample types was observed for enhancers (Fig. 4D). During embryogenesis, the highest proportion of Shared enhancers occurred at late somitogenesis, reaching a proportion similar to brain - the highest among adult tissues (Fig. 4D). Mature testis showed the highest proportion of Exclusive enhancers, while liver and gonads showed the lowest proportion of Shared enhancers (Fig. 4D).

Thus, the retention of enhancers duplicated following the salmonid 4R WGD was strongly shaped by ontogeny and tissue-specific context, whereas promoter evolution was less dynamic.

### Enhancer evolution is coupled to ohnolog expression regulation

*Cis*-regulatory elements govern transcriptional regulation (*47*, *49–51*), and in mammals, enhancer evolution is linked to rates of gene expression evolution (*46*). However, the association between enhancer evolution and ohnolog expression divergence following WGD remains poorly understood. We addressed this knowledge gap, using analyses that did not split genes by rediploidization history to exploit the stratification of ohnolog sequence divergence when making inferences on regulatory changes.

We first asked if enhancer alignment category is correlated with ohnolog regulatory evolution across sample types. We observed that as ohnolog expression divergence increased, the proportion of Shared enhancers decreased, while the proportion of Alignable only and Exclusive enhancers increased (Table S14). While the observed correlation directions are consistent with enhancers contributing to ohnolog regulatory divergence, each test failed to reached significance (Pearson’s R = -0.41, 0.39, 0.37 for Shared, Alignable only and Exclusive enhancers, respectively; each *p-*adj > 0.17). However, significant correlations were observed when performing the same tests for promoters, in the same direction as enhancers (respective Pearson’s R = -0.72, 0.78 and 0.59; *p-*adj = 0.004, 0.002 and 0.02 for Shared, Alignable only and Exclusive promoters) (Table S14). Thus, the functional retention of promoters following WGD is more directly linked to ohnolog regulatory evolution than enhancers.

However, the above analyses indirectly test for a relationship between enhancer evolution and regulatory divergence among ohnolog pairs at the sample level. As a second more direct test, we defined *cis*-regulatory domains for all salmon genes then sub-divided ohnolog pairs into “conserved” and “diverged” regulation classes based on expression correlation across tissues and development (Fig. 5A-B, Fig. S36), before comparing the composition and conservation of enhancers assigned to ohnolog pairs from each class. This distance-based approach assumes gene transcription is regulated by nearby enhancers within the defined *cis-*regulatory domains. While some of these gene-to-enhancer assignments are likely inaccurate, e.g. due to long-distance regulation, the proximity of enhancers to promoters has been shown to be a highly accurate predictor for their physical interaction (*52*), with the same approach used for cross-species analyses (*46*).

**Fig. 5.**
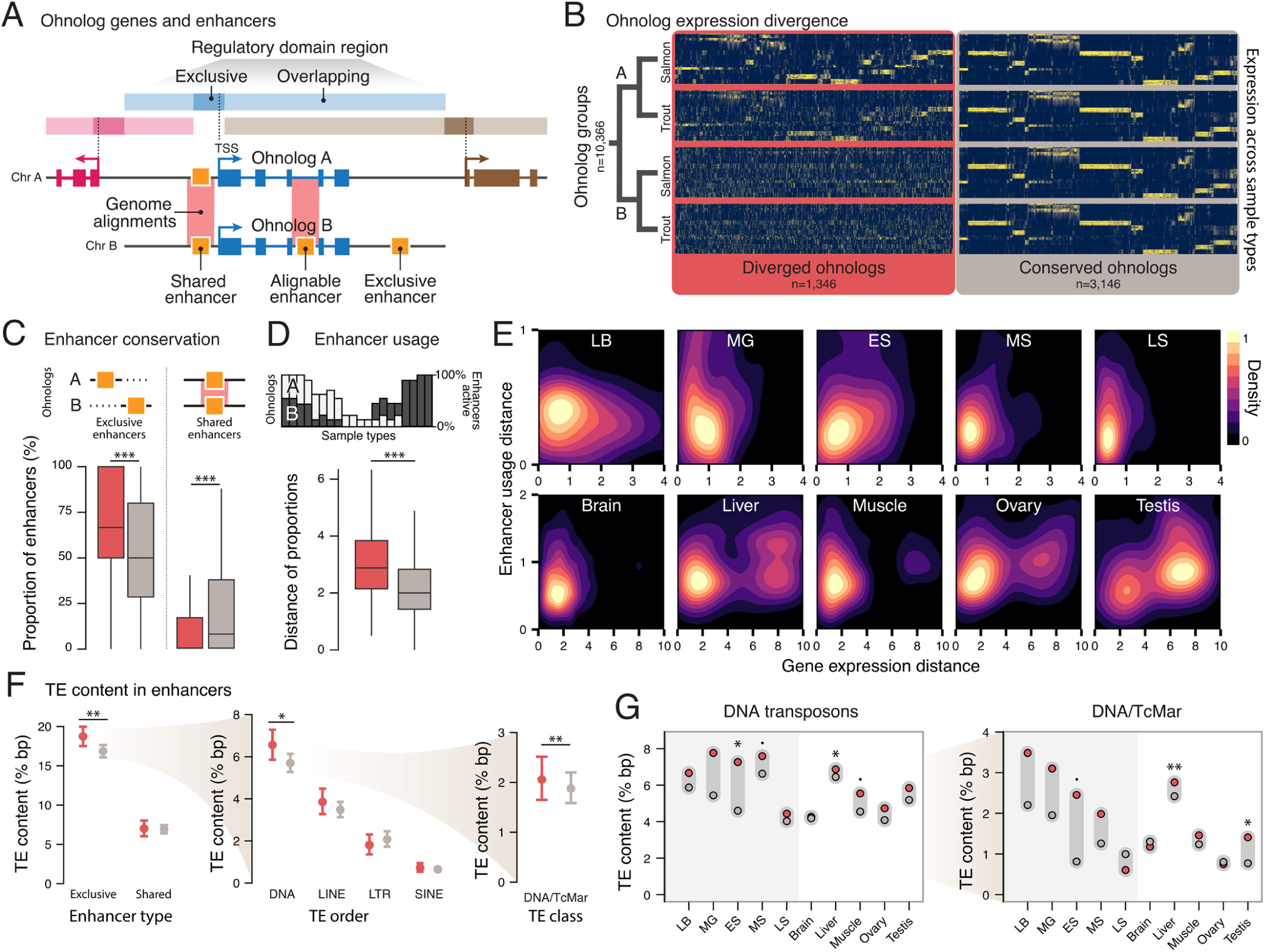
Enhancer evolution is coupled to ohnolog regulatory divergence. **(A**) Our definition of *cis*-regulatory domains (*46*) of genes includes an ‘exclusive’ region - 5 to + 1 kb surrounding the TSS, and an ‘overlapping’ region, extending until a neighboring exclusive region or maximum distance of 1 Mb from the TSS. Active enhancers were compared within these domains for ohnolog pairs from the Shared, Alignable only and Exclusive categories. (**B**) Ohnolog pairs were split into “conserved” (gray) or “diverged” (red) regulation sub-classes (see Fig. S37), which were compared in terms of: (**C**) proportions of Exclusive and Shared enhancers and (**D**) Enhancer usage distance, representing total enhancer activity profiles across sample types (expressed as Euclidian distance). (**E**) Density maps of ohnolog pair enhancer-usage Euclidian distance vs. expression Euclidian distance, split by embryogenesis stage or adult tissue type. (**F**) Percentage of TE basepairs within Exclusive or Shared enhancers for ohnologs from both regulation sub-classes, shown for all TEs, then specifically for Exclusive enhancers by TE order (DNA, LINE, LTR, SINE), and finally the DNA transposon TcMar. (**G**) TE content in Exclusive enhancers across embryogenesis stage or adult tissue types. Symbols: * = p < 0.05, ** = p < 0.01, *** = p < 0.001).

We found that ohnolog pairs showing conserved regulation had more active enhancers (*p* = 2.2 x 10^-16^, Wilcoxon test) (Fig. S37). They also showed a higher proportion of Shared enhancers (*p* = 8.0 x 10⁻²⁶, odds ratio = 2.04) and reduced proportion of Exclusive enhancers (p = 5.8 x 10⁻³⁷; odds ratio = 0.62) (Fig. 5C). Breaking down enhancer characteristics by sample type, ohnologs with conserved expression had higher levels of Shared enhancers active in the same samples (Fig. S37, Wilcoxon test: *p* = 1.2 x 10⁻⁶). On the other hand, Alignable only enhancers were equally present in ohnologs with diverged or conserved regulation (Fig. S37).

To further explore the link between enhancers and ohnolog pair expression divergence, we calculated differences in total enhancer activity profiles across sample types, quantified by Euclidean distance (termed ‘enhancer-usage distance’, see Methods) (Fig. 5D). Ohnolog pairs showing divergent expression had higher enhancer-usage distances than pairs showing conserved expression (Fig. 5D; respective medians = 1.15 vs 0.89; *p* = 1.6 x 10⁻²⁸). We further asked if enhancer usage distance is associated with ohnolog pair expression distance (see Methods; Fig. 5E, Fig. S38). While ohnolog expression distance decreased across embryogenesis, consistent with our JSD-based analysis (Fig. 3), enhancer usage distance was more variable. This suggests ohnolog expression divergence during embryogenesis is mainly constrained by Shared enhancers, rather than total enhancer activity profiles. Conversely, we observed stronger across-tissue differences in both ohnolog expression distance (Kruskal-Wallis test χ² = 166.6, df = 4, *p* < 2.2 × 10⁻¹⁶) and enhancer-usage distance (χ² = 77.2, df = 4, *p* = 6.7 x 10⁻¹⁶). Brain showed the lowest distance and dispersion in both cases, whereas liver, ovary, and testis displayed the broadest range of values (Fig. S38).

Overall, these results reveal that the evolution of ohnolog expression is associated with the loss of ancestral enhancer sequences or the gain of new enhancer sequences post-WGD.

### Transposable elements have shaped enhancer evolution after WGD

One mechanism to evolve new enhancer sequences is through transposable element (TE) activity. Indeed, the salmonid WGD was associated with a burst of TE activity, which led to increased rates of TE derived *cis*-regulatory element evolution (*52*). On the other hand, TEs also disable functional elements within genomes, meaning their overlap with regulatory elements provides insights into constraints on gene regulation across ontogeny or tissues. Yet, how TEs contribute to enhancer functional evolution following WGD remains poorly understood. To explore these questions, we identified all active enhancers within *cis*-regulatory domains from salmon that overlapped with a high-quality TE annotation (*52*). Overall, we found that Exclusive enhancers overlapped TEs to a much larger extent (>2 fold) than Shared enhancers (Fig. 5F). This could be explained by TE insertions either acting to create *de novo* enhancers (*53*, *54*), or by reduced constraint on the functions of Exclusive enhancers.

To assess the contribution of TEs to ohnolog regulatory divergence, we contrasted the TE-sequence content in enhancers of ohnolog pairs with conserved or diverged regulation (Fig. 5F-G; Fig. S39). The proportion of TE sequences in Exclusive enhancers was weakly associated with ohnolog regulatory divergence (Fig. 5F, *p* = 0.0031). However, this was not the case for Shared enhancers (Fig. 5F). The most active TE group following WGD in salmonids are DNA transposons, specifically the TcMar (DNA/TcMar) (*52*). In line with this, the overlap of DNA TEs with Exclusive enhancers was increased for diverged ohnolog pairs (Fig. 5F; *p* = 0.038 for all DNA elements; *p* = 0.0051 for DNA/TcMar elements), which was not the case for retrotransposons. Further analysis highlighted an increased overlap between DNA or DNA/TcMar TEs and Exclusive enhancers for ohnologs showing divergent regulation in early somitogenesis, liver and testis (Fig. 5G, Fig. S39).

Past work has hypothesised that the contribution of TEs towards gene regulation following the salmonid WGD was shaped by tissue-specific selective constraint (*53*). We tested this hypothesis using our panel of tissues and embryonic stages. Enhancers active in brain and late embryogenesis, representing sample types where ohnolog expression divergence and enhancer functional conservation were most constrained following WGD (Fig. 3-4), showed the lowest overall overlap with TEs for enhancers located within both the Early and Late rediploidization categories (Fig. S40). Further, as the overlap between TEs and active enhancers increased across sample types, so did the proportion of Alignable only and Exclusive enhancers, while the proportion of Shared enhancers decreased (Fig. S40; Table S15).

Together, these data indicate that TE insertions broadly contribute to regulatory evolution following WGD, acting to disable duplicated enhancers and create new enhancers. Moreover, selection against TE insertions appears strongest for enhancers retained as duplicates, especially those active in sample types under the most constrained regulatory evolution following WGD.

### Ontogeny governs the evolution of enhancer-CNEs following WGD

The strongest retention of Shared enhancers following WGD was for late embryogenesis and brain (Fig. 4C). A past mammalian study profiled different tissues across ontogeny to reveal that enhancers most active in brain were also most conserved across species, especially at stages showing prominent brain development (*55*). Building on this finding, we asked if the conservation of enhancer sequences across species influences their evolution following WGD, and if so, whether this is shaped by distinct constraints across ontogeny and tissue types.

Here, we exploited the fact that conserved non-coding elements (CNEs) often lie within enhancers governing development and tissue identity, marking functionally important regulatory sequences under purifying selection across species (*56*, *57*). We overlapped all salmon active enhancers with a genome-wide dataset of northern pike CNEs stratified by phylogenetic age, with the oldest category predating the split of ray- and lobe-finned fish (Fig. 6; Table S16). We investigated the retention of salmon CNEs located within active enhancers (hereafter: enhancer-CNEs) following WGD by their alignment to pike CNEs (Table S17). Using this approach, Exclusive enhancer-CNEs represent singletons, where one in a former ohnolog pair was lost, while the retained CNE is conserved with pike. Among 24,007 high-confidence enhancer-CNEs, 61.4%, 31.2% and 7.3% belonged to the Shared, Alignable only and Exclusive categories, respectively (Fig. 6A). Thus, 92.6% of all enhancer-CNEs were retained as duplicated sequences from WGD, with activity in both duplicates (Shared category) being around twice as common as activity restricted to one locus (Alignable only category).

**Fig. 6.**
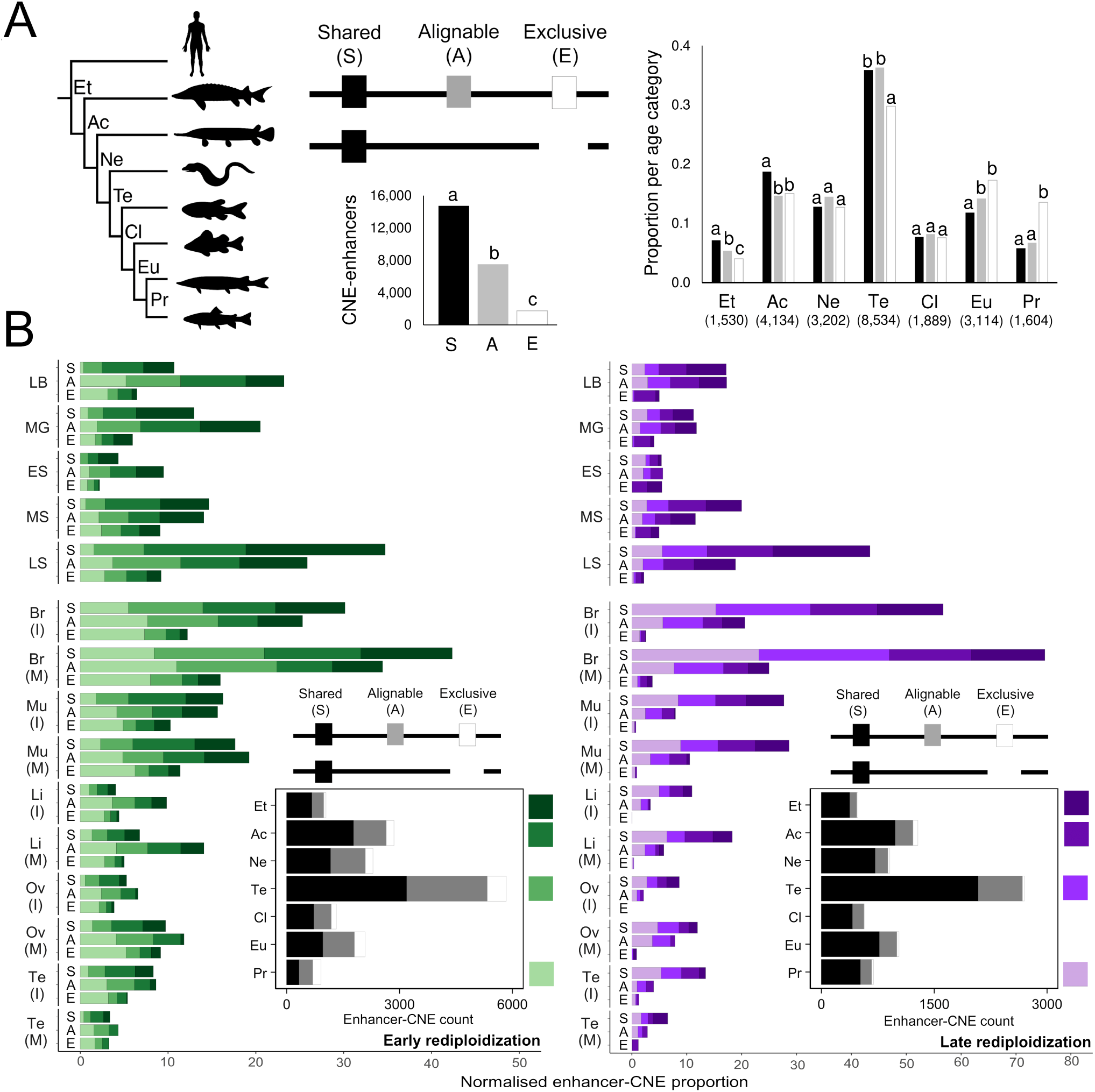
Evolution of enhancer-CNEs following the salmonid 4R autotetraploidization. (A) The tree on the left illustrates the different phylogenetic ages of enhancer-CNEs: ‘Et’ (Euteleostomi), ‘Ac’ (Actinopteri), ‘Ne’ (Neopterygii), ‘Te’ (Teleostei), ‘Cl’ (Clupeocephala), ‘Eu’ (Euteleostomorpha), and ‘Pr’ (Protacanthopterygii). Also shown is the number of Atlantic salmon enhancer-CNEs belonging to the Shared (S), Alignable only (A), and Exclusive (E) categories, along with their proportions per phylogenetic age category, within each alignment category. On the provided barplots, different lowercase letters indicate significant differences between alignment categories within each provided comparison (Fisher’s Exact Test). (B) Normalised proportions of Atlantic salmon enhancer-CNEs for each alignment category (S, A, E), stratified by phylogenetic age and different sample-types for Early (left) and Late (right) rediploidization regions. Sample abbreviations are as per the Fig. 1 and 2 legends, with maturation stage split by ‘I’ (immature) and ‘M’ (mature) for tissue samples.

The most common enhancer-CNEs (35.5%) belonged to the teleost age category (Fig. 6A; Fig. S41). Significant overrepresentation of enhancer-CNEs from the Shared category was observed for the two oldest CNE categories, comprising 23.6% of all enhancer-CNEs (Fig. 6A). Thus, selection on the retention of enhancer-CNEs following WGD was strongest for the most ancient vertebrate enhancers.

Conversely, the small number of Exclusive enhancer-CNEs were enriched in the youngest CNEs and underrepresented in CNEs of the teleost age category (Fig. 6A). Shared enhancer-CNE sequence pairs showed higher sequence conservation than the Alignable only category, both overall (respective median nucleotide sequence identity: 95.0 vs. 92.9%; Wilcoxon test, *p* = 2.2 x 10^-16^) or stratifying by CNE age category (Fig. S41). For enhancer-CNEs from the Alignable only category, the CNE that lost enhancer activity has evolved significantly faster (reduced nucleotide percentage identity with single orthologous pike CNE), with the youngest CNEs most affected (Fig. S41). Thus, the phylogenetic age and functional retention of enhancer-CNEs strongly shapes their ongoing sequence evolution following WGD.

Enhancer-CNEs were strongly overrepresented in Late rediploidization regions, overall and for each phylogenetic age category (Table S18). This effect was entirely explained by Shared enhancer-CNEs, which were overrepresented overall (Fig. S42), and per age category, while enhancer-CNEs from the Alignable only and Exclusive categories were enriched in Early regions (Fig. S42; Table S19). Remarkably, Shared enhancer-CNEs within Late regions comprised 39.9% of all Shared enhancer-CNE space in the genome, greater than two-fold higher than expected by chance based on the length of these regions (Table S19). This overrepresentation strongly suggests selection acted against the rediploidization and divergence of these elements during the early stages of post-WGD evolution.

Finally, we partitioned enhancer-CNEs from Early and Late rediploidization regions across alignment categories, CNE phylogenetic ages, and sample types (Fig. 6B). These data were normalised to allow statistical comparisons in all combinations (Fisher’s Exact Test results: Tables S20-22). A strong enrichment of Shared enhancer-CNEs from the two oldest CNE categories was observed at late somitogenesis compared to earlier embryonic stages for both Early and Late regions (Fig. 6B; Table S22; *p =* 0). Specific to the oldest CNE category, in both Early and Late regions, the proportion of Shared enhancer-CNEs at late somitogenesis was significantly higher than brain (Fig. 6B; Table S22; *p =* 0). Comparing adult tissue types, brain captured the highest proportion of enhancer-CNEs for both rediploidization regions, with a significantly higher proportion of Shared-enhancer CNEs from the teleost-specific and Protacanthpterygii categories compared to embryonic stages and other adult tissues (Fig. 6B). In Late rediploidization regions, Shared enhancer-CNEs dominated all CNE age categories except early stages of embryogenesis (Table S21). In Early regions, a smaller proportion of enhancer-CNEs were active in liver, testis and ovary compared to muscle and brain (Fig. 6B).

Overall, these results highlight a key role for development and ontogeny in shaping the evolution of enhancer-CNEs following WGD. The striking overrepresentation of the oldest CNE category in Shared enhancer-CNEs active at late somitogenesis is consistent with post-WGD constraints on ancient regulatory programs governing vertebrate organogenesis (*40*), and predicts that similar patterns will follow other vertebrate WGD events.

## Discussion

This study paints a novel perspective of genome evolution following autotetraploidization, where ohnolog regulatory evolution is governed by the same ontogenetic and tissue-specific constraints acting between species. Multiple lines of evidence capture a period of within-genome constraint on regulatory evolution at late embryogenesis, which can be explained by the pleiotropic regulatory networks governing organogenesis at this life stage in all vertebrates (*40*). Similarly, in adult tissues, the brain represents the maxima of post-WGD regulatory constraint, while gonads and liver experience weaker regulatory selection (*44*, *55*). While our study advances understanding of functional evolution following the salmonid 4R WGD, additional data types can penetrate additional levels of regulation, including 3D chromatin architecture, while resolving changes to specific cell types (*58*).

Our findings illustrate the importance of better capturing rediploidization processes in future studies of regulatory evolution following WGD. This comes into sharp focus in light of recent discoveries that asynchronous rediploidization followed other autotetraploidization events (*6*, *15*, *24*) and may represent a ‘rule’ of genome evolution in vertebrates, and possibly other eukaryotic lineages. Thus, the stratification of regulatory divergence we describe in salmonid genomes may have occurred in many genomes, including during earlier stages of vertebrate diversification. In this respect, salmonids provide an ideal vertebrate system to further explore the regulatory basis of functional innovations derived from autopolyploidization, including lineage-specific novelties arising following delayed rediploidization (*17*).

Finally, our study marks the sharing of an open genomic resource in highly processed formats, ready for analysis by the research community and other stakeholders. This resource can be readily leveraged to support precision breeding goals in salmonid aquaculture (*28*, *29*), for instance by helping to dissect traits with a regulatory basis, supporting the long-term sustainability of global food production. These datasets will also enable explorations of the role played by regulatory elements in the adaptation of wild salmonid populations to rapidly changing environments across their natural ranges, while enriching comparative analyses of regulatory evolution across vertebrate lineages.

## Supporting information

Supplementary Materials including Methods and Supplementary Figures

## Acknowledgments

We thank the various members of the AQUA-FAANG consortium working on other teleost species for valuable discussions that shaped the direction of this study.

## Funding

This study was funded by the AQUA-FAANG project, which received funding from the European Union’s Horizon 2020 research and innovation programme under grant agreement No 817923 (www.aqua-faang.eu). Further support was provided by the Biotechnology and Biological Sciences Research Council (BBSRC), including Institute Strategic Programme grants BBS/E/RL/230001B and BBS/E/D/10002070 and a Strategic Longer and Larger Award to DJM (BB/Z51746X/1). DB was supported by EU funds through the CroAGE project (code: NPOO.C3.2.R2-I1.06.0060 NextGenerationEU). BZ was supported by EU funds through the croESTRO project (code: NPOO.C3.2.R2-I1.06.0024 NextGrenerationEU).

## Author contributions

Conceptualization: DJM, SL, YG, SRS, JB, BL, MPK

Methodology: M-OB, DPM, GG, PSD, MKG, TP, LG, DB, AL(au), FG, BZ, EC-G, AP, AB, TTN, DT, HRC, YG, SRS, BL, JB, MPK, SL, DJM

Investigation: M-OB, DPM, GG, PSD, MKG, TP, LG, DB, AL(au), FG, BZ, EC-G, AP, AB, TTN, GAM, EP, DT, GRI, AL(ou), TRH, HRC, YG, SRS, BL, JB, MPK, DJM.

Visualization: M-OB, DPM, GG, PSD, MKG, TP, LG, DB, SRS

Funding acquisition: DJM, SL, YG, SRS, JB, BL, MPK Project administration: DJM, SL, MKP, M-O, PWH

Supervision: DJM, SL, TRH, CB, PWH, DT, HRC, YG, JB, SRS, BL, MKP

Writing – original draft: DJM, M-OB, DPM, GG, PSD, MKG, LG, DB, SRS

Writing – review & editing: all authors reviewed and approved the final manuscript

## Competing interests

The authors declare that they have no competing interests.

## Data and materials availability

All sequencing datasets produced in this study are available through the FAANG data portal (https://data.faang.org/projects/AQUA-FAANG). ENA accession numbers for different studies are shared in Table S2, and as follows: <u>PRJEB51855</u>, <u>PRJEB51854</u> and <u>PRJEB53399</u> (Atlantic salmon DevMap: RNA-seq, ATAC-seq and ChIP-seq, respectively); <u>PRJEB51857</u>, <u>PRJEB51856</u> and <u>PRJEB55010</u> (rainbow trout DevMap: RNA-seq, ATAC-seq and ChIP-seq, respectively); <u>PRJEB47409</u>, <u>PRJEB47408</u> and <u>PRJEB55063</u> (Atlantic salmon BodyMap: RNA-seq, ATAC-seq and ChIP-seq, respectively); <u>PRJEB57191</u>, <u>PRJEB57190</u> and <u>PRJEB57956</u> (rainbow trout BodyMap: RNA-seq, ATAC-seq and ChIP-seq, respectively). All code required to reproduce the analyses and result reported is provided at: https://github.com/AQUA-FAANG/AQUA-FAANG-salmonids.

## Notes

### Competing Interest Statement

The authors have declared no competing interest.

### Summary of Updates

Author names corrected in this version only.

