## Supplementary Materials including Methods and Supplementary Figures for "Salmonids reveal principles of regulatory evolution following autotetraploidization"

**Includes:**

Materials and Methods
Supplementary Text
Figs S1 to S42

**Attached elsewhere**

Tables S1 to S22
Data S1 to S5

### Materials and Methods

#### Full protocols and sample information

In sections that follow, we provide an overview of several key steps of our study, including sampling, sequencing library preparation, and sequencing for both target salmonid species. These descriptions are based on detailed protocols provided in the Supplementary Text (Text S1-4). Information about sample types and biological replicates per species and sequencing assay can be found in Supplementary Tables S1-S2. A visual summary of all protocols is provided in Fig. S1.

#### Sampling

*DevMap sampling*

A full sampling protocol is provided in the Supplementary Text (Text S1). Briefly, eggs from a single Atlantic salmon or rainbow trout female were fertilized with milt from a single Atlantic salmon or rainbow trout male, respectively, leading to monoparental batches of fertilized eggs. Fertilized eggs were incubated at 8°C (Atlantic salmon) or 9°C (rainbow trout) in tray systems receiving recirculating, filtered water. At each sampling time point, eggs were collected then processed/stored for subsequent RNA extraction and library preparation for RNA-seq, ATAC-seq and ChIP-seq (see later sections).

For RNA extraction, three eggs were punctured and submerged in 1,000 µL of TRIzol™ (ThermoFisher) in 1.5 mL Safe-Lock Eppendorf™ tubes with homogenization beads (27 tubes sampled per developmental stage). This was followed by homogenization using a TissueLyser II (Sigma) for 2 mins at 30hz and storage at -80°C. For ATAC-seq library preparation, eggs were dechorionated and the embryo separated from the yolk sac using watchmaker forceps. After gentle mincing with a scalpel, the embryo material was digested using 0.25% Trypsin EDTA. Following PBS washes, the embryos were cryopreserved in 300 µL cryopreservation solution (10% DMSO, 40% FBS, 50% DMEM) using 2 mL tubes and a Mr Frosty freezing container (Nalgene®) with a 1°C/min cooling rate at -80°C. n=3 biological replicates were sampled per developmental stage. The number of embryos required per biological replicate was as follows: blastula stages: 40; gastrula stages: 30; early somitogenesis: 20; mid-to-late somitogenesis: 10; eyed stage/pharyngula: 5. The volume of 0.25% Trypsin EDTA and duration of trypsinization also varied for each stage, as described within Text S1. Embryo material was prepared for ChIP-seq as described above for ATAC-seq prior to the digestion step, followed by crosslinking using 875 µL of 1% formaldehyde. Crosslinking was stopped using 125 µL of 1M Glycine, followed by resuspension in 1 mL of PBS and storage at -80°C. n=3 replicates were sampled for each developmental stage. The number of embryos used per ChIP-seq replicate varied by developmental stage as for ATAC-seq.

*BodyMap sampling*

A full sampling protocol is provided in Text S1. n=12 Atlantic salmon and n=12 rainbow trout were used. In both cases this included n=3 fish per sex (i.e., male, female) for two stages of ontogeny (sexually immature, sexually mature). The maturation status of all the sampled fish was established by visual inspection of the gonads, followed by histological analysis (images provided within full protocol described in Text S1). Fish were fasted for 24 hr before a lethal/non-recovery dose of MS222 was administered, followed by killing using a schedule 1 method. Seven tissues were immediately sampled (in order: brain, liver, gonads, distal intestine, gill, muscle), cut in pieces, washed in cold PBS then placed in RNAlater solution at 4˚C overnight (RNA-seq) or snap-frozen on dry ice (ATAC-seq, ChIP-seq) then stored at -80˚C. The brain was collected following removal of the pituitary gland and brain stem, with remaining regions thoroughly mixed before splitting each sample. A strip from the middle of the liver was extracted, avoiding the vascular network. A slice (maximum 1 cm thick) from the middle of the gonad was placed in Dietrich solution for 10 days then dehydrated with ethanol washes, embedded in paraffin, and cut into 4 µm sections using a microtome. Eosin staining was used to assess gonadal maturation state (images provided within protocol described in Text S1). The distal intestine samples were collected after gentle removal of faecal remains and mucus. The head kidney tissue was carefully removed, washed in PBS and cut into slices for sampling. Gill samples were restricted to the filaments, with rakers discarded. Fast-twitch ('white') skeletal muscle was sampled from the anterior epaxial myotomal muscle adjacent to the dorsal fin.

The number of biological replicates used from the above samples for RNA-seq, ATAC-seq and ChIP-seq is provided per species, sex and maturation status in Table S1.

**Functional assays and sequencing**

*Total RNA extraction - DevMap*

A full protocol is provided within Text S2. Briefly, total RNA extraction was performed after thawing the homogenized embryo samples (in TRIzol™; see above) at room temperature, followed by centrifugation to separate the aqueous phase from the organic phase, debris and oils. Phase separation was achieved by adding 200 µL of 1-Bromo-3-Chloropropane, followed by vortexing for 15s and incubation at room temperature for 30 min. After centrifugation, 400 µL of the aqueous layer was extracted. RNA precipitation was achieved by the addition of 200 µL of a 1:1 solution of isopropanol:RNA precipitation solution (1.2M Sodium Chloride, 0.8M Potassium Citrate Sesquihydrate). Following overnight precipitation at -20ºC, samples were centrifuged and pellets washed in 70% ethanol under rotation for 30 min. Three such ethanol washes were performed. Pellets were resuspended in 20-200 µL of nuclease-free water (volume used depending on pellet size) at room temperature for 60 mins, flicking the tubes every 10 mins. RNA concentration was established using a Qubit 4 fluorimeter (Invitrogen) with a Broad Range RNA kit. RNA purity was confirmed with a Nanodrop 1000 spectrometer (Thermo Fisher Scientific). RNA integrity was confirmed using a Tapestation 4200 (Agilent), recording RNA integrity (RIN) scores for all samples.

*Total RNA extraction - BodyMap*

A full protocol is provided within Text S2. Briefly, total RNA extraction from tissues of Atlantic salmon was performed using a column-based miRNeasy Mini kit (Qiagen, #217004) following the manufacturer's instructions. Each tissue was removed from RNAlater solution, placed in 700 µL Qiazol with one 5 mm stainless steel bead (Qiagen, #69989), homogenized with a TissueLyser (Qiagen) for 15-40 s (until the solution appeared clear), then mixed with chloroform and centrifuged at 12,000g for 15 min. The upper phase was extracted and mixed with 100% ethanol, passed through the RNeasy filter included in the kit, then washed, DNAse-digested (Qiagen, #79254) and washed again. An optional column drying (17,000g, 1 min) after the washing steps was applied to improve RNA purity. Total RNA samples were eluted in 50 µL water, and their quality assessed using a Nanodrop 1000 spectrometer and Bioanalyzer 2100 system (Agilent), before storage at -80°C.

Total RNA extraction from tissues of rainbow trout was performed using a phenol-chloroform extraction method. Tissue samples (100 mg) were submerged in 1 mL Trizol (Euromedex, #TR118) and homogenized (Precellys, Bertin Instruments, # P000669-PR240-A) in Eppendorf tubes for 40 s with ceramic beads (1.4mm, #P000933-LYSK0-A.0 OZYME). 200 µL of chloroform was added, before the tubes were inverted several times, and the mixed samples incubated for 5 mins at room temperature before centrifugation (12,000g at 4°C for 30 mins). The upper phase was transferred to a new tube, mixed with isopropanol (500µL per 1 mL Trizol) by vortexing, incubated for 2 hr at -20°C, before centrifugation at 12,000g (4°C for 25 mins). The resultant pellet was carefully washed twice with 1 mL of 75% ethanol. The washed pellet was dried and resuspended in ultra-pure water prior to quality assessment, as for Atlantic salmon above.

#### Library preparation and sequencing

*RNA-seq - DevMap and BodyMap*

RNA-seq library preparations were performed by Novogene using total RNA samples described above from both species (Tables S1, S2). In all cases, the NEBNext® Ultra™ Directional RNA Library Prep for Illumina® kit (#E7420S/L) was used with the NEBNext Poly(A) mRNA Magnetic Isolation Module (#E7490) and NEBNext® Multiplex Oligos for Illumina® (#7600). All libraries were produced following manufacturer's instructions. Briefly, mRNA was isolated from 0.1-1 µg of total RNA using the poly(A) mRNA enrichment step, fragmented and random primed prior to first and second strand cDNA synthesis step. Next, 3' ends were end-repaired and dA-tailed, ligated to the adapter and size selected, PCR-amplified and cleaned-up. Libraries were sequenced on the Illumina Novaseq 6000 platform generating paired-end 150bp reads.

*ATAC-seq - BodyMap*

A full protocol including tissue-specific modifications is provided within Text S3. Briefly, ATAC-seq was based on Omni-ATAC-seq (*59*), a modified version of ATAC-seq (*32*, *60*) optimized for frozen tissues. Omni-ATAC-seq is divided into three main steps: nuclear isolation, transposition and PCR with size selection. Nuclear isolation involves tissue disruption, filtration and homogenization, release of nuclei from cells, then purification and concentration of nuclei suspension. Optimizations of nuclear preparations were applied to different tissues (see Text S3). For example, in the case of mature testis, additional washes and a second gradient centrifugation improved the ratio of nuclei from testis vs. contaminating sperm cells. Alternatively, for muscle, tissue disruption required grinding on dry ice to effectively disrupt the muscle fibres and release the nuclei (*61*).

For each processed sample, nuclei were counted and confirmed for quality before transposition, done using Illumina Tagment DNA Enzyme (transposase) and Buffer (Illumina, 20034197). The transposition reaction was sensitive to the ratio of transposase to nuclei. We found the transposition to be optimal using 50,000 nuclei per 2.5 µL transposase, except for gill (25,000 nuclei per 2.5 µL transposase (see Text S3). Following 30 mins transposition at 37ºC, the DNA was purified using a MinElute PCR Purification Kit (Qiagen, #28004) and frozen at -20ºC. The above steps were performed in tissue-specific batches, then, once a larger number of eluted DNA was accumulated, the samples were PCR-amplified (8-13 total PCR cycles) and size-selected (180-700bp). The number of PCR cycles and size selection steps were optimized to fit the best product size distribution for sequencing. The DNA profile (nucleosome banding) was confirmed using a High Sensitivity Bioanalyzer (Agilent). The concentration of the libraries was assessed by using High Sensitivity Qubit 4.0 kit. Libraries were sequenced on the Illumina Novaseq 6000 platform by Novogene.

*ATAC-seq – DevMap*

A full protocol is provided within Text S3. Briefly, the ATAC-seq method used with embryo samples was an adaptation of the BodyMap Omni-ATAC method, differing in the steps leading up to nuclear isolation, which were based on a published protocol used for single nuclei RNA-seq (*62*). Cryopreserved embryo samples (see above section) were thawed on ice by pipetting up and down in 900 µL of ice-cold PBS containing Protease Inhibitor Cocktail (PIC, Sigma) This step was immediately followed by a round of centrifugation and resuspension of the resultant pellet in 1,000 µL of PBS containing PIC. After a further round of centrifugation, nuclei isolation was performed by resuspending the pellet in 1,000 µL of Tween with salts and Tris (TST) buffer containing PIC, followed by pipetting up and down on ice for 5 mins. The nuclei isolation was stopped by the addition of Salts and Tris (ST) buffer, and centrifuged (see Text S3). 75,000 nuclei were for the transposition reaction, following the Omni-ATAC-seq protocol described for BodyMaps above. Libraries were sequenced on the Illumina Novaseq 6000 platform by Novogene.

*ChIP-seq – DevMap*

A full protocol is provided within Text S4. Briefly, we used a ChIPmentation approach with all DevMap samples. ChIPmentation is a tagmentation-based ChIP-seq method that allows one-step library preparation on bead-bound chromatin (*63*). We used a µChIPmentation Kit for Histones (Diagenode #C01011011), optimized for low input samples, such as those from embryos. Briefly, crosslinked embryo samples were thawed on ice and washed once with ice-cold PBS prior to homogenization in 1 mL of homogenization buffer (PBS and 1x Protease Inhibitor Cocktail, PIC) using a 2 mL Dounce homogenizer (20-60 cycles on ice).

We used Chromatin EasyShear Kit - High SDS (Diagenode # C01020012) for cell lysis and chromatin shearing. 5 x $10^5$ homogenized cells were lysed in 315 µL of tL1 lysis buffer with 1 x PIC on ice for 20-30 mins. After complete lysis was observed by microscope, 185 µL of HBSS buffer was added and samples were sonicated using a Bioruptor Pico bath sonicator. Samples were sonicated in 0.65 mL Bioruptor® Microtubes #C30010011 for 3-7 cycles (1 cycle = 30 sec 'on' followed by 30 sec 'off') until chromatin fragments, as analysed using a Tapestation 4200 (High-Sensitivity D1000 assays), were in the 200-800 bp size range.

We followed Diagenode's instructions for µChIPmentation with the following modifications: i) pull-down was done using 100 ng of sheared chromatin per assay (equivalent to approximately 37,000 cells) and ii) 0.8-1 µg antibody was used (H3K27ac #C15410196, H3K4me1 #C15410194, H3K4me3 #C15410003, H3K27me3 #C15410195) per reaction. For library preparation, we used 24 unique dual indexes for tagmented libraries Set I and Set II from Diagenode (#C0101134 and #C0101135); samples were tagmented at 37°C for 12 mins and amplified for 14-15 PCR cycles. Libraries were sequenced on an Illumina Novaseq 6000 by Novogene, generating 150bp paired-end reads (targeting 45 million reads per ChIP sample,). For each condition, a fraction of chromatin was extracted before immunoprecipitation for control (6 µL), and underwent the µChIPmentation protocol in a similar way as for the samples, with some adaptations (see Text S4). An equimolar mix of input controls per developmental stage was created for sequencing (90M reads per input control sample).

*ChIP-seq – BodyMap*

Full protocols including tissue-specific modifications are provided within Text S4. The Diagenode µChIPmentation approach, as used for DevMaps, was used for brain tissue in rainbow trout. For all other tissues from both species, ChIP-seq was performed using frozen tissues, with an optimized classic protocol involving seven steps: tissue disruption and homogenization, crosslinking, sonication, immunoprecipitation, washing, de-crosslinking and elution of immunoprecipitated DNA. The first step aimed to produce a homogenous and clean isolation of nuclei, which required additional modifications for muscle and mature male gonads. Counting of nuclei was performed before crosslinking. After crosslinking, multiple washes were immediately performed to clean the nuclei. Next, samples were concentrated to

10 million nuclei / mL (400 µL total volume) and sonicated with a probe sonicator (EpiShear, ActiveMotif, #53052), with all parameters fixed (55% intensity, 1 sec ON/1sec OFF cycle rhythm, 30 sec cycle) except the number of cycles, which was tissue-specific. After sonication, the sample was centrifuged for 10 min at 8,000g to separate sonicated chromatin from debris. To assess the profile of sonicated chromatin using an Agilent Bioanalyser, an aliquot of each sample was de-crosslinked, and the DNA purified and quantified. Samples with a size distribution of 100-600 bp (median: 350bp) were diluted 4-fold with immunoprecipitation (IP) buffer before storage at -80ºC, prior to immunoprecipitation.

Immediately before immunoprecipitation, two separate preparation steps were performed in parallel: i) chromatin pre-clearing (30 min incubation of chromatin with Protein A/G beads without antibody) and ii) coupling of antibodies to the Protein A/G beads, which was subsequently used in an overnight IP step (4ºC, in rotation) with pre-cleared chromatin. An aliquot (5 ng) of precleared chromatin from each sample was not immunoprecipitated, and used as an input control (each control library was produced using a pool of group-specific input samples). Four antibodies (Diagenode) were used with following amounts per immunoprecipitation: 1µg for H3K4me1 (#C15410194) and H3K4me3 (#C15410003), and 2µg for H2K27ac (#C15410196) and H3K27me3 (#C15410195). Next, immunoprecipitated samples were washed multiple times with graded salinity buffers, allowing removal of weak bonds between Protein A/G beads-antibody complexes and unspecific/non-targeted protein epitopes, before the immunoprecipitated chromatin was eluted from the beads and de-crosslinked, its DNA purified using a MinElute PCR Purification Kit (Qiagen, #28004) and its concentration assessed using a Qubit High Sensitivity DNA Kit (Thermofisher, #Q32854).

Immunoprecipitated DNA was stored at -20ºC until library preparation, done using a MicroPlex Library Preparation Kit V3 (Diagenode; #C05010002), following the manufacturer's recommendations. Briefly, a minimum of 0.2 ng of immunoprecipitated DNA was end-repaired and ligated with stem-loop adaptors through blunt-end ligation, then dual-indexed (96 Dual indexes for MicroPlex Kit v3 Set I-IV,# C05010004- C05010007) and PCR-amplified (from 7 to 8 cycles for inputs and from 8 to 13 cycles for immunoprecipitated products), purified (Agencourt AMPure XP Beads, Beckman Coulter, #A63880) and sequenced on an Illumina Novaseq 6000 platform by Novogene, generating 150bp paired-end reads (targeting 45 million reads per ChIP sample, 90 million reads per input control sample).

### Primary data analyses

*Reference genomes*

The reference genomes and annotations used for all reported data analyses were downloaded from the Ensembl rapid release database; for Atlantic salmon: assembly Ssal_v3.1, softmasked fasta file (Salmo_salar-GCA_905237065.2-softmasked.fa) and gene annotation GTF file (Salmo_salar-GCA_905237065.2-2021_07-genes.gtf.gz). For rainbow trout: assembly USDA_OmykA_1.1, softmasked fasta file (Oncorhynchus_mykiss-GCA_013265735.3-softmasked.fa) and gene annotation GTF file (Oncorhynchus_mykiss-GCA_013265735.3-2020_12-genes.gtf).

*RNA-seq alignment and quantification*

RNA-seq data was processed using a standardised approach through nf-core pipeline (*64*, *65*) rnaseq v3.8.1 (*66*) with standard parameters. Briefly, the pipeline involves adapter and quality trimming of reads (Trim Galore!) (*67*), alignment of reads to the reference genome using STAR (*68*) before projection of alignments to the transcriptome for BAM-level quantification of gene and transcript level read counts using RSEM (*69*). MultiQC (*70*) reports were produced and assessed for all samples.

*ATAC-seq alignment and peak calling*

ATAC-seq data was processed using a standardised approach through nf-core pipeline atacseq v1.2.1 (*71*) with standard parameters. Briefly, the pipeline involves adapter and quality trimming of reads (Trim Galore!), alignment of reads to the reference genome using the Burrows-Wheeler Alignment (BWA) tool (*72*) and (narrow) peak calling using MACS2 (*73*, *74*). The estimated genome size used was: 2,756,563,003 for Atlantic salmon, and 2,070,645,207 for rainbow trout. MultiQC reports were produced and assessed for all samples.

*ChIP-seq alignment and peak calling*

All ChIP-seq data was processed using a standardised approach through the nf-core pipeline chipseq v1.2.2 (*71*) with standard parameters. Briefly, the pipeline involves adapter and quality trimming of reads (Trim Galore!), alignment of reads to the reference genome using BWA, and peak calling using MACS2 with representative ChIP input control samples used to establish background signal. The estimated genome size used was as detailed above for ATAC-seq. Broad peaks were called for H3K4me1, H3K27me3 and narrow peaks for H3K4me3, H3K27ac. MultiQC reports were produced and assessed for all samples.

*Blacklist regions*

To remove signal-artefact regions from the sequencing data for downstream analysis, we generated blacklist files for the Atlantic salmon and rainbow trout genomes. First, uint8 mappability files were created for 75, 100, and 150 k-mers across each chromosome in the genome using Umap (*75*) then all k-mer mappability results were combined. Finally, blacklist regions were defined using Blacklist (*76*) taking in all available ChIP-seq input BAM files (32 and 29 respective input controls for Atlantic salmon and rainbow trout) and combined uint8 files. Blacklists were generated separately for classic ChIP (BodyMap) and μChIPmentation (DevMap) input controls, with the resulting blacklisted regions merged.

*Chromatin state annotations*

Chromatin states were annotated across the Atlantic salmon and rainbow trout genomes using ChromHMM Input signal data were BAM alignments for the ATAC-seq and ChIP-seq (H3K27ac, H3K4me1, H3K4me3, and H3K27me3 marks) data generated as described above. Sampled matched input controls were given to estimate background signals for the ChIP-seq data. Prior to ChromHMM, BAM alignments were converted to BED format and filtered to keep reads only in chromosomes and exclude blacklist regions. The alignments were binarized using the ChromHMM *BinarizeBed* function with default bin size (200 bp), before running the ChromHMM *LearnModel* function. After testing models with various state numbers, with the aim to capture the most biologically relevant chromatin states (*35*) we proceeded with: two separate 12-state models for Atlantic salmon DevMaps (i.e. embryo stages) and BodyMaps (i.e. tissue samples; brain, gonad, liver, muscle); and for rainbow trout, a 12-state model for DevMaps, and three separate 10-state models for each included tissue (brain, liver, muscle) (Fig. S10A). Note that each rainbow trout tissue was treated separately, as data quality was more variable between tissues compared to Atlantic salmon. After generating the ChromHMM models, the non-blacklisted BAM files representing ATAC-seq and multiple histone mark ChIP-seq data were binarized using the ChromHMM *BinarizeBam* function with default bin size (200 bp). The combination of binarized signals from ATAC-seq and ChIP-seq marks were used together with the ChromHMM model for the respective species and condition to annotate regions across the genome representing different chromatin states at 200 bp resolution (see Results, main article).

*Robust open chromatin definition and functional annotation*

A set of robust open chromatin regions, defined as those that are highly reproducible across biological replicates per condition, was generated using the ATAC-seq narrow peak files resulting from the analyses described above, post-filtering to remove peaks either not found on chromosome sequences, or completely overlapping blacklist regions. The reproducibility of peaks between all possible pairs of biological replicates per condition was determined using the Irreproducible Discovery Rate (IDR) approach (*77*) as done by the ENCODE project. IDR measures the reproducibility of the scoring of peaks between replicates to determine an optimal cut-off for peak significance. IDR was run using options --rank='p.value' and --soft-idr-threshold=0.1. This gave a set of significantly reproducible peaks between pairs of replicates. To deal with having three biological replicates per condition, the resulting reproducible peaks from each pair were merged together with bedtools *merge* (=>1bp overlap), taking the maximum IDR score for the resulting peak. With one set of reproducible peaks per condition, the summits of those peaks were re-calculated using the MACS2 *refinepeak* function and all replicate ATAC-seq BAM alignments were merged together.

Finally, the reproducible ATAC-seq peaks with summits were assigned with chromatin states per condition by intersecting ChromHMM annotations using bedtools *intersect*. When a peak overlapped more than one ChromHMM annotation in a given condition, the assigned chromatin state was chosen in order of importance, based on how confidently we could call the state given the combination of assay signals required. This order is the same order of the chromatin states shown in Fig. 1B (see main text) from top to bottom. A single unified set of robust open chromatin regions (reproducible ATAC-seq peaks) was generated per species by merging all overlapping peak regions (=>1bp overlap) across all conditions. Read counts from each ATAC-seq and ChIP-seq assays were counted with *featureCounts* using the BAM alignments for each biological replicate against the unified peak set of each species. This table of assay counts within robust open chromatin peaks for all samples was used for downstream analyses.

**Transcription factor binding site (TFBS) enrichment analysis**

TFBS enrichment analysis was carried out with *Gimme maelstrom* (GimmeMotifs v0.17.1 (*78*)) separately on the identified active enhancer and active promoter regions of Atlantic salmon and rainbow trout. The input was log-transformed ATAC-seq counts (pooled across biological replicates) for robust open chromatin peaks described above. The analyses were run with default parameters, except with the region size set to 200 bp. The motif database used was JASPAR2022_vertebrates.pfm. By investigating the results, we found that several closely-related Jaspar TFBS were associated with different, but closely-related TFBS showing an enrichment in matched sample types for Atlantic salmon and rainbow trout. To reconcile this discrepancy for the purpose of comparative analysis, we downloaded a table including pairwise correlation values for all Vertebrate JASPAR TFBS (*79*). TFBS that shared a correlation value $\geq 0.9$ were considered a single TFBS for the purpose of comparison between Atlantic salmon and rainbow trout, which led to an increase in the number of shared enriched TFBS.

**Homology prediction**

Ensembl Compara (*80*) 106 gene trees (downloaded from http://ftp.ensembl.org/pub/misc/aqua-faang/ensembl_trees/gene_trees/Compara.106.protein_default.nhx.emf.gz) were used as the basis to identify: i) ohnolog pairs retained from the salmonid-specific WGD and ii) orthologs conserved between Atlantic salmon and rainbow trout and with northern pike (*Esox lucius*) as an outgroup species to the WGD. Estimation of homology relationships is obscured by incorrect gene-tree estimation, gene loss and lineage-specific rediploidization, where ohnologs independently diverged in sequence after the two salmonid species diverged (*17*, *20*). To validate that duplicated genes identified in the Ensembl gene trees derived from the salmonid-specific WGD, we leveraged large collinear/syntenic blocks in which the vast majority of these genes reside (*18*). Our approach firstly extracts Protacanthopterygii clades from the gene trees, i.e. including both salmonids and northern pike, a representative of Esociformes; the sister lineage to salmonids, which did not experience the salmonid-specific WGD (*19*). If both ohnologs have been retained in Atlantic salmon and rainbow trout, each clade will contain two genes per species, which we call '2:2 clades'. Using the 2:2 clades, we identified syntenic blocks by selecting the most commonly observed combinations of four chromosomes (>20 clades with the same combination of chromosomes) (Fig. S15). We assign the ortholog relationship of the four chromosomes using the most commonly observed phylogeny of clades in agreement with ancestral ohnolog resolution (AORe) (*17*) i.e. requiring two salmonid sister clades, each containing one Atlantic salmon and one rainbow trout ortholog. In the main text, we also call these Early rediploidization regions. In genomic regions that have undergone lineage-specific rediploidization, we observe two clades showing lineage-specific ohnolog resolution (LORe) (*17*) i.e. where ohnolog pairs from each species are monophyletic, because they diverged after speciation. In the main text, we also call these Late rediploidization regions. The syntenic relationship of 2:2 clades was used to verify the synteny of gene tree clades where two ohnologs were not retained in both species, including (among others): 1:1 clades (e.g. resulting from one ohnolog in a pair being lost ancestrally to Atlantic salmon and rainbow trout), 2:1 clades (where one ohnolog was lost in one species, and the other species retains both ohnologs) and 2:0 clades (e.g. when both ohnologs were lost in one species, and the other species retains both ohnologs). The number of genes belonging to all possible type of clade is shown in Fig. S15 and the full homology prediction is shared in Table S5 and available on SalmoBase.org (*34*).

**Whole genome alignment to explore regulatory element evolution**

Whole genome alignments including Atlantic salmon, rainbow trout, and northern pike were generated using progressive Cactus (*38*). As Cactus was not designed to align genomes affected by recent WGD events, we were concerned that running Cactus without adaptations using salmonid genomes would fail to effectively capture duplicated regions. As part of the European AQUA-FAANG project, a Cactus alignment was produced for six teleost species: Atlantic salmon, rainbow trout, along with turbot, European seabass, Gilthead seabream and common carp (available at http://ftp.ensembl.org/pub/misc/aqua-faang/cactus/). This six-species alignment was run in default mode, where the whole genome of each species was fed into Cactus. Inspection of the results revealed sub-optimal capture of duplicated regions, particularly in more divergent AORe regions (Fig. S16), which comprise much of salmonid genomes (*20*).

To confirm the sub-optimal capture of duplicated regions, we devised then tested an alternative strategy where collinear/syntenic regions retained from the salmonid WGD were extracted as separate regions for Atlantic salmon, rainbow trout (i.e., two ohnologous collinear regions from each species) and northern pike as an outgroup. Genome-wide collinear and syntenic blocks for the Atlantic salmon, rainbow trout, and northern pike genomes were identified by aligning the chromosomes of each species to the Atlantic salmon chromosomes using minimap2 (v 2.18-r1015) (*81*). Manual inspection of dot plots from the alignments facilitated the identification of conserved collinear blocks within the duplicated Atlantic salmon and rainbow trout genomes, and allowed for the determination of their orthologous regions in the northern pike genome. A final table, containing details of all the syntenic/collinear blocks with respect to each Atlantic salmon chromosome, is presented in

Table S6. As an initial test system, we extracted two collinear/syntenic regions associated with Atlantic salmon chromosome 3, namely Ssa03 and Ssa06 (homologous to regions of Omy12 and Omy13) and Ssa03 and Ssa14 (homologous to regions of Omy28 and Omy08) using the synteny data generated above (Table S5). We deliberately selected these regions as they represent AORe (Ssa03-14 and Omy28-08) and LORe (Ssa03-06 and Omy12-13) regions of similar sizes (*18*), meaning they are markedly more and less diverged as duplicated sequences, respectively (*17*). For each syntenic block, the extracted sequences were fed into Cactus as independent sequences, regardless of duplicated sequences originating within each species. Cactus was run with default parameters to generate a HAL alignment output for each syntenic block. The HAL output files for both syntenic blocks (Ssa03-Ssa06 and Ssa03-Ssa14) were converted to MAF alignment format using the hal2maf command *( --maxRefGap 20000 --noAncestors --noDupes and rest as default*) in the ComparativeGenomicsToolkit (*82*), facilitating downstream processing. Atlantic salmon chromosome 3 (Ssa03) was set as the reference chromosome for both tested syntenic blocks during the HAL to MAF conversion. The mafStats command within mafTools (*83*) was used to generate summary statistics for each MAF file, including the number of base pairs for each non-reference sequence (duplicated sequences) captured in the MAF alignment (Fig. S16). We found that the capture of duplicated regions in both species was markedly improved by alignment of the extracted syntenic regions compared to the standard Cactus approach (Fig. S16). The gain in recovered sequence was especially striking for the more divergent AORe (Early rediploidization) regions, and more modest, yet still substantial for highly similar LORe (Late rediploidization) regions (Fig. S16).

For our final Cactus alignment strategy, as described above, we segmented the Atlantic salmon and rainbow trout genomes into collinear/syntenic regions retained from the salmonid WGD and identified orthologous regions in the northern pike genome (Table S6). The resulting sequence regions were used for independent Cactus alignments for each Atlantic salmon chromosome. A custom workflow was developed to automate this process. Briefly, the pipeline uses each Atlantic salmon chromosome as a reference, denoted by the prefix Ssal_A. The duplicated salmon chromosome is assigned the prefix Ssal_B, while syntenic rainbow trout sequences are labelled Omyk_A and Omyk_B, and the orthologous northern pike sequence is denoted as Eluc. For each reference chromosome (Ssal_A), the syntenic/orthologous regions are extracted (extractBlocks.R) and aligned in Cactus (run_cactus.job.sh) with the assigned prefixes. The HAL output files were converted to MAF format using the hal2maf command in the Comparative Genomics Toolkit (*84*). A custom Python script (fix_maf.py) was used to adjust alignment coordinates from block-level to genome-wide coordinates, producing a final set of MAF alignment files, one per Atlantic salmon chromosome.

**Self-organizing maps and Uniform Manifold Approximation and Projection**

To determine how chromatin accessibility and gene expression were regulated across ontogeny, we clustered the Atlantic salmon and rainbow trout ATAC-seq and RNA-seq datasets using self-organizing maps (SOM). For the ATAC-seq data, we first extracted raw read counts for all robust open chromatin regions. To account for differences in sequencing depth, tags per million (TPM) values were calculated by normalizing read counts to the total number of mapped reads per sample while correcting for region length. To ensure comparability between samples, we calculated the fold change of observed over expected values for each open chromatin region, using the mean TPM value within the sample as the expected value. The resulting fold-change values were then scaled by dividing by the standard error across samples, ensuring comparability of both regions and conditions. For the RNA-seq data, we calculated the mean of biological replicates of transcripts per million

(TPM) values for all samples. We independently performed quantile normalization across DevMap and BodyMap samples using the preprocessCore R package (*85*) to correct for potential discrepancies in sequencing depth or experimental variability. As with ATAC-seq peaks, the standard error across samples was used to divide the gene expression values. Following this data pre-processing, using a hexagonal configuration, SOM clustering was performed using the kohonen R package (*86*). The described procedure was performed on the ATAC and RNA-seq DevMap and BodyMap datasets for Atlantic salmon and rainbow trout. The number of clusters was manually optimized to capture maximal variability in each dataset without incurring large differences in cluster size or redundant cluster profiles. The following SOM configurations were used: i) 4x4 for DevMap RNA-seq (i.e. 16 SOM clusters), ii) 6x6 for BodyMap RNA-seq (i.e. 36 SOM clusters), iii) 5x5 for DevMap ATAC-seq (25 clusters) and iv) 5x5 for BodyMap ATAC-seq (25 clusters). Cluster profiles were further visualized with heatmaps, after ordering clusters within each analysis based on a combination of maximally expressed samples, mean expression across samples and standard deviation across samples. This approach allowed us to segregate clusters into stage-specific and constitutive categories for DevMap samples and tissue-specific, multi-tissue expression and constitutive expression categories for BodyMap samples.

In addition, to visualize the dynamic developmental trajectory and to differentiate samples per tissue, we applied the Uniform Manifold Approximation and Projection (UMAP) algorithm (*87*) for dimensionality reduction on each processed matrix using the UMAP implementation in the uwot package (*88*) providing a two-dimensional visual representation of the data, highlighting underlying patterns and structures in our high-dimensional dataset.

*Cross-species SOM cluster comparisons*

To assess the conservation of gene expression and chromatin accessibility between Atlantic salmon and rainbow trout, we performed an enrichment analysis of orthologs across self-organizing map (SOM) clusters in both species. The analysis was conducted separately for the DevMap and BodyMap datasets, using both RNA-seq and ATAC-seq data. For this analysis, we used only 1-to-1 gene orthologs and robust ATAC-seq peaks located in syntenic regions within the same ancestral collinear block, defined by our Cactus whole-genome alignments. For each clustering, we compared all pairs of SOM clusters between Atlantic salmon and rainbow trout. We generated matrices representing the observed and expected number of orthologous genes for RNA-seq and syntenic ATAC-seq peaks for chromatin accessibility. The size of these matrices varied depending on the SOM configuration, reflecting the number of clusters in each dataset. The observed number of orthologs in each cluster pair was counted directly from the data, while expected values were estimated using a permutation-based approach, in which cluster assignments were randomized 10,000 times to generate a null distribution of orthologous counts. The mean number of orthologs across these permutations was used as the expected value for each cluster pair. Enrichment was calculated as the ratio of observed to expected orthologs, providing a measure of conservation between SOM clusters. To visualize the relationships between clusters, we generated heatmaps using the ComplexHeatmap R package (*89*), in which rows corresponded to SOM clusters in Atlantic salmon and columns to SOM clusters in rainbow trout, with colour intensity reflecting the observed-to-expected enrichment ratio. We generated Sankey plots using the ggalluvial R package (*90*) to illustrate shared orthologous genes and open chromatin region between species, with nodes representing SOM clusters and edges weighted according to the number of orthologs connecting each pair.

*Gene set enrichment for SOM clusters*

For SOM clusters produced using RNA-seq datasets, we investigated enrichment of particular biological functions among gene sets comprising each SOM cluster. This analysis was done separately per species and each major dataset (i.e. DevMap and BodyMap) using the gene ontology (GO) framework and the package clusterProfiler (*91*) restricting to 'biological process' terms and using Atlantic salmon and rainbow trout annotations produced by Ensembl, which were loaded into R using AnnotationForge (*92*) and biomaRt (*93*). For DevMaps, GO enrichment was performed on each SOM cluster. For BodyMaps, SOM clusters were manually curated according to tissue-specific expression, with SOM clusters showing constitutive expression used as an additional category. For each test, the background genes in the test were all genes expressed above 1 TPM in at least one sample for each dataset separately. Enriched GO terms were filtered using a custom script and the package GO.db (*94*) to retain only parent terms, therefore reducing the presentation of highly redundant GO terms. Shared GO terms among SOM clusters were visualized using the clusterProfiler function *compareCluster*.

**Validation of ChromHMM annotations using CAGE**

CAGE-seq followed a previously published protocol (*95*) with sequencing compatibility optimizations. To improve sequencing efficiency and demultiplexing accuracy on Illumina platforms, the legacy 3-bp barcode at the start of Read 1 was replaced with 8-bp TruSeq Unique Dual Indexes. Adapter ligation and reverse transcription were followed by an additional five to six cycles of PCR amplification using NEBNext Ultra II Q5 Master Mix (New England BioLabs, #M0544S) to incorporate Illumina P5/P7 flow cell adaptors and I5/I7 indexes. Library purification was performed using 1.4× AMPure XP beads (Beckman Coulter, #A63881) to remove adapter dimers and short fragments. Final libraries were quality-checked using a High Sensitivity DNA Bioanalyzer (Agilent, #5067-4626) and quantified with a Qubit dsDNA High Sensitivity Assay Kit (Thermo Fisher, #Q32854). Libraries were sequenced on an Illumina NextSeq 2000 with the flow cell P4-50 kit, generating 70 bp single-end reads. Following sequencing, raw reads were demultiplexed using standard Illumina processing pipelines. Adapter sequences and low-quality reads were trimmed and filtered using Trim Galore! with default settings. Quality-filtered reads were aligned to the Salmon and Trout reference genomes using Bowtie2 v2.5.0 (*96*) in end-to-end alignment mode, ensuring that the 5' end of each read was correctly mapped to its genomic location. Alignments were filtered to retain only reads with mapping quality greater than 20, ensuring only primary, uniquely mapped reads were retained before being converted to BAM format using SAMtools v1.5.

Mapped data were processed using CAGEr v2.11.4 (*97*) following the recommended workflow. First, the artificial G nucleotide added during reverse transcription was removed from the 5' end of each read to correct for CAGE-specific biases. A probabilistic approach was used to distinguish added matching Gs from non-added matching Gs. Signal normalization was performed using power-law normalization, and transcription start site (TSS) clusters were identified with a clustering threshold of 20bp. Clusters were further filtered to remove artefacts and low-expression peaks, and genomic coordinates were converted to a standardized format for quality validation of ChromHMM annotations.

**Ohnolog regulatory divergence analysis**

We calculated pairwise expression distances for defined classes of ohnolog and ortholog pairs (data shown in Fig. 3). We first calculated the mean of biological replicates of TPM values for each gene, separately for different samples. We then independently performed quantile normalization across DevMap and BodyMap samples using the preprocessCore R package (*98*). Next, the standard error across samples was divided by the gene expression values.

Ohnologs and orthologs were extracted from our homology prediction (Table S5) for comparison. Specifically, we compared expression data four non-overlapping gene-pair categories: (i) 8,757 2:2 AORe (Early rediploidization) pairs, which started diverging in the Atlantic salmon and rainbow trout ancestor; ii) 1,103 2:2 LORe (Late) rediploidization) pairs, where two ohnolog pairs are conserved in both Atlantic salmon and rainbow trout, but diverged independently in each species; (iv) 8,661 ortholog pairs representing AORe 2:2 ohnologs and (v) 5,099 ortholog pairs that represent singletons from AORe regions. Within each category, the orthogroup ID was used to segregate gene pairs into separate columns (named geneA and geneB). Expression divergence between the two members of a pair was measured using Jensen-Shannon distance (JSD). Prior to distance calculation, the two normalized TPM profiles were jointly quantile-normalized using the preprocessCore R package and re-scaled by dividing each value by the sum of all values. JSD values were then obtained using the JSD function of the phylentropy R package (*99*). To capture uncertainty, 100 bootstrap replicates were drawn for every embryonic stage (DevMaps) or tissue/maturity (BodyMaps) combination by resampling gene-pair indices with replacement. Within each replicate we recalculated JSD for the sampled set and subsequently recorded the mean and standard deviation across replicates. To assess the correlation of divergence patterns between categories, we calculated Pearson's correlation between the sample-wise mean JSD values; scatter plots, regression lines and 95% confidence intervals were produced with the ggpubr R package (*100*).

**Evolution of regulatory elements**

Genomic coordinates of the unified set of 319,718 Atlantic salmon robust open chromatin regions were provided as a BED file to the mafExtractor command within mafTools, allowing us to extract each robust open chromatin region within our whole genome alignment in MAF format. A custom R script was used to process each resulting alignment file to extract the start and end coordinates for sequences (Ssal_A, Ssal_B, Omyk_A, Omyk_B, Eluc) present in the MAF file. Tabular data from individual chromosomes was merged into a single dataset and curated to remove spurious alignments, including nested alignments where Ssal_B coordinates were identical for multiple robust open chromatin region (9,250 regions), and instances where a single robust open chromatin region overlapped two regions on the same chromosome (4,996 regions). De-duplication was performed to eliminate duplicate representation of robust open chromatin regions across the two duplicated chromosomes. The final output captures the genomic coordinates of 225,801 reference robust open chromatin regions (Ssal_A) along with the coordinates of their syntenic/orthologous regions (Ssal_B, Omyk_A, Omyk_B, Eluc) (Supplementary Dataset 1).

We grouped the robust open chromatin regions into three categories: 1) "Shared", representing robust open chromatin regions conserved in syntenic sequences located on two duplicated chromosomes represented in the whole genome alignment; 2) "Alignable only", representing syntenic sequences conserved on two duplicated chromosomes represented in the whole genome alignment, but with a robust open chromatin region found on only one of these duplicated regions; and 3) "Exclusive", representing robust open chromatin regions specific to a single chromosome (i.e. no evidence that the region is retained as a duplicated sequence from the salmonid WGD). A schematic of these regions is provided in Fig. 4 of the main text.

To identify "Shared" regulatory elements, robust open chromatin regions overlapping both duplicated regions (i.e., Ssal_A and Ssal_B) were selected, and the percentage overlap between duplicated robust open chromatin regions calculated. The summit regions for robust open chromatin regions (defined from ATAC-seq peaks) were overlaid onto these overlapping regions to filter for high-confidence regulatory elements, where the summit coordinates fell within the overlapping regions for both duplicates (Ssal_A and Ssal_B). For "Exclusive" regulatory elements, the alignments were filtered to retain regions where both the sequence and robust open chromatin regions were confined to Ssal_A or Ssa_B. Similarly, "Alignable only" regions were identified by filtering for alignments where sequence conservation was observed across both duplicated chromosomes (Ssal_A and Ssal_B), but a robust open chromatin region was present only in Ssal_A or Ssa_B.

We next overlapped the robust open chromatin regions from each category with our ChromHMM annotation to identify those representing active promoters (45,802/225,801) or active enhancers (78,424/225,801) across the unified dataset spanning all samples. This identified 16,938/14,836, 6,845/19,546 and 7,873/10,232 active promoters/enhancers belonging to the “Shared”, “Alignable only”, and “Exclusive” categories, respectively (Supplementary Dataset 2). The promoter and enhancer elements within these three categories were further split into Early and Late rediploidization categories using the genomic coordinates across the Atlantic salmon genome (see Table S9). The activity of categorized regulatory elements (presence/absence), was assessed for individual sample types in the BodyMap and DevMap atlases. Specifically, for active promoters and enhancers, we calculated the proportion of each within the “Shared”, “Alignable only”, and “Exclusive categories” across each DevMap stage of embryogenesis and across different BodyMap tissue/maturation stage samples, independently for Early and Late rediploidization regions (results shown in Fig. 4; Table S12). Fisher's exact tests (with Bonferroni correction) were performed to assess statistical significance across different sample types for all possible combinations within each alignment category, and separately for early and late rediploidization regions (Table S13). Pearson’s correlation was performed across all sample types and per rediploidization category to compare the proportions of active promoters or enhancers within the “Shared,” “Alignable only,” and “Exclusive” categories) vs. JSD values for 2:2 ohnologue pairs in salmon (see previous section) (Table S14).

*Transposable element content within regulatory elements*

A curated TE dataset for Atlantic salmon (*53*) was used as input to RepeatMasker to annotate TE regions across the salmon genome. TE annotations were filtered to remove low complexity, satellite, and simple repeats. TE content was calculated as the percentage of active enhancer bases overlapping TE sequences out of the total sum of active enhancer bases. This was calculated separately for active enhancers found in each sample type, and those classified into the “Shared”, “Alignable only” and “Exclusive” categories. Pearson’s correlation was performed across all sample types and per rediploidization region to compare the proportion of active enhancers within each alignment category vs. the proportion of TEs overlapping active enhancers (Table S15).

*Evolutionary fate of enhancer-CNEs*

CNEs were defined in the northern pike genome (fEsoLuc1.pri; GCA_011004845.1) based on a whole genome alignment of 46 species including human alongside a broad selection of actinopterygian lineages (teleosts and non-teleosts), inclusive of Atlantic salmon and rainbow trout. The approach to call CNEs and identify their phylogenetic age followed Naville et al. (2015) (*101*). Using the northern pike genome as a reference allowed us to i) identify northern pike CNEs in our whole genome alignment and ii) leverage their co-orthology to putative Atlantic salmon CNE ohnologs. The approach in full is as follows.

Starting from a 46-species Multiz alignment with fEsoLuc1 as the reference genome, we used GERP++ gerpcol (*102*) to compute the GERP conservation score (i.e. number of rejected substitutions) at all positions, then filter out aligned columns to keep only those fulfilling all the following conditions: 1) belong to a conserved element with significant GERP score (FDR: 0.2); (2) do not overlap with any annotated exon for any of the aligned genomes, nor with any annotated pike repeat sequence; (3) still belong to an aligned block of at least 10 consecutive bases after filtering steps 1 and 2. For each of these blocks, we attempt to assign putative target gene(s) using synteny conservation: among all pike genes located within a fixed radius around the block, we choose as putative target(s) the one(s) which are most often also found in the vicinity of the orthologous block in other genomes. Then, blocks less than 100bp away and targeting the same genes in the same species were merged into single units that we consider CNEs. Therefore, one CNE can be made of several conserved non-coding blocks. After this merging process, we obtained 416,589 CNEs in the northern pike genome (Supplementary Dataset 3), which spanned 21,136,463 bp. Each of these CNEs was assigned a putative age, which corresponds to the most recent ancestor of all the species which possess an orthologous CNE: Euteleostomi (ancestral to ray-finned fish and mammals); Actinopteri (ancestral to ray-finned fish, excluding Polypteriformes); Neopterygii (ancestral to holostean and teleost fish); Teleostei (ancestral to teleost fish); Clupeocephala (ancestral to clupeocephalan teleosts, excluding Osteoglossomorpha and Elopomorpha); Euteleosteomorpha (ancestral to euteleost fish, excluding Otocephala) and Protacanthopterygii (ancestral to protacanthopterygians, including Esociformes and Salmonidae).

We categorized and extracted CNEs captured in our whole genome alignment, requiring that 100% of the northern pike CNE sequence was captured and located within an alignment belonging to the Shared, Alignable only, or Exclusive categories, previously categorized as representing active enhancers in Atlantic salmon. These extractions were performed using Maftools to generate maf files capturing individual CNEs. At this stage, we identified 21,928, 14,040, and 2,565 individual enhancer-CNEs belonging to the Shared, Alignable only, and Exclusive categories, respectively. Each MAF alignment was subsequently processed through an R-based filtering pipeline applying the following criteria to retain highly credible CNEs: 1) At least 90% of the CNE sequence length was present among any of the northern pike or Atlantic salmon sequences within each MAF alignment block; 2) The alignment block contains zero gaps for any northern pike and Atlantic salmon sequences captured; 3) At least one Atlantic salmon co-ortholog must exhibit >80% sequence similarity to the pike ortholog. Final counts and proportions of Atlantic salmon enhancer-CNEs belonging to each alignment category were provided across the genome and separated by the Early and Late rediploidization regions (Table S17), with information provided for individual CNEs in Supplementary Dataset 4.

Nucleotide sequence percentage identity was separately calculated between Atlantic salmon enhancer-CNE ohnolog pairs (Atlantic salmon A vs. B) and compared between the Shared and Alignable only categories. To test if the rate of sequence evolution in enhancer-CNE ohnologs depends on enhancer-CNE activity, we firstly calculated nucleotide sequence percentage identity separately for each enhancer-CNE within an ohnolog pair in comparison to their shared pike CNE ortholog (i.e., Atlantic salmon A vs. pike and Atlantic salmon B vs. pike). We then compared differences in sequence divergence against pike for salmon A and B ohnologs for the Shared (A active, B active) and Alignable only (A active, B inactive) categories. Each of the above analyses was done globally, then stratified by each phylogenetic age of CNEs, with differences between groups of interest assessed using

Wilcoxon's rank-sum test, using the Bonferroni-Holm method to obtain adjusted P values accounting for multiple comparisons.

*Enhancer-CNE activity across ontogeny*

To explore the impact of ontogeny, tissue-context and rediploidization on the evolution of enhancer-CNEs following the salmonid WGD, we calculated the number and length of enhancer-CNEs per: i) each sample-type defined within our DevMap and BodyMap atlases (averaged across sex for the BodyMap data), ii) each alignment category (Shared, Alignable only, Exclusive) and iii) each rediploidization region (Early and Late). The results are shown in Supplementary Dataset 5. To normalize the above comparisons, CNE counts were divided by the total length of CNEs across the three alignment categories for each phylogenetic age across all sample types. The normalized proportion of enhancer-CNEs in the Shared, Alignable only and Exclusive categories was separately plotted for Early and Late rediploidization regions per phylogenetic age and sample type. Fisher's Exact test was used to test for enrichment of enhancer-CNEs in Early vs. Late rediploidization regions, comparing the length of enhancer-CNEs overlapping each region, against the total length for Early (2,023,914,877 bp) and Later regions (649,544,050 bp) (Table S18). This analysis was repeated after splitting enhancer-CNEs within each age category into the Shared, Alignable and Exclusive alignment categories (Table S19).

Fisher's exact test was also used to compare 1) all possible CNE phylogenetic age combinations per sample-type, alignment category and rediploidization region (Table S20). 2) all possible Alignment categories per sample type, rediploidization region and CNE phylogenetic age category (Table S21) and 3) all possible sample-types per enhancer-CNE phylogenetic age class, rediploidization region and alignment category (Table S22).

**Linking enhancer evolution to ohnolog regulatory divergence**

A series of steps were performed to investigate the extent to which divergence in the expression of salmon ohnolog pairs was coupled to divergence of *cis*-enhancer elements proximal to these genes.

The first step was to define regulatory domains for ohnolog genes, following Berthelot et al. (2018) (*46*). Each gene's regulatory domain was defined by an 'exclusive' region 5kb upstream and 1kb downstream of its transcription start site (TSS). Next, an 'overlapping' region extended up and downstream of the exclusive region, until meeting another gene's exclusive region, or a maximum distance of 1 Mb from the TSS. The exclusive and overlapping regions together defined the regulatory domain. Exclusive regions can overlap each other, and overlapping regions can overlap each other, but overlapping regions cannot overlap exclusive regions.

The second step was to annotate *cis*-enhancers by overlapping robust open chromatin regions with active enhancer annotations from our ChromHMM annotation across all sample types. Gene regulatory domain regions were used to assign the enhancers falling within the domains to the respective gene. With this approach, enhancers can be assigned to more than one gene when domains overlap. All enhancer ohnolog pairs were assigned to the Shared, Alignable only, or Exclusive category using our whole genome alignment, as described above. When enhancers were located outside a gene's regulatory domain, yet aligned to an enhancer located within the other ohnolog's regulatory domain, they were taken forward in the analysis.

The third step was to delineate ohnolog pairs into two extreme categories of expression conservation: "conserved" and "diverged". This was done using Pearson's correlation with scaled TPM values averaged across biological replicates for all sampled conditions (i.e., all

DevMap stages of embryogenesis, plus the different BodyMap tissues from the two maturation stages), for both ohnolog pairs of the same species, and between 1:1 ortholog pairs between species, using our homology prediction (Table S5). We used Pearson correlation value thresholds to classify 10,366 ohnolog groups (2:2 ohnolog pairs from both species) into the conserved and diverged groups. A set of 3,146 ohnolog groups were defined as "conserved", based on Pearson correlation values > 0.7 for the 2:2 ohnolog pairs of both salmonid species and > 0.5 between the equivalent ortholog pairs across species. A set of 1,346 ohnolog groups were defined as "diverged", based on Pearson correlation values ≤ 0.3 for the 2:2 ohnolog pairs of both salmonid species.

Regulatory landscape differences between 2:2 ohnologs were compared between the "diverged" and "conserved" expression groups from Atlantic salmon. For statistical analysis, unless stated otherwise, the Wilcoxon rank sum test was chosen for tests of differences between the distribution of values of "conserved" and "diverged" ohnologs, with 95% confidence intervals computed from median values using bootstrapping with 10,000 replicates. The number of enhancers per gene was compared between "conserved" and "diverged" ohnologs. Differences in enhancer membership to the alignment categories (Exclusive, Shared, Alignable only) between "conserved" and "diverged" ohnolog pairs were tested using binomial generalized linear models (GLMs), fitted for enhancer proportions with ohnolog conservation status as the predictor, reporting odds ratios with 95% confidence intervals. Multiple comparisons were Holm-adjusted, and global differences across categories additionally tested using a 3×2 Chi-square test. Similarity in the activity of Shared enhancers was compared by dividing the number of times Shared enhancers of an ohnolog enhancer group were annotated as active by ChromHMM in the same sample type, over the total unique number of samples the shared enhancers were active. We compared differences between ohnologs in their 'enhancer-usage' across conditions. For this purpose, we calculated, per sample type, the proportion of enhancers active as a product of the total number of active enhancers for each gene. At least one active enhancer for each gene was required to test an ohnolog pair. The Euclidean distance between the two vectors of proportions was calculated between ohnolog pairs, termed 'enhancer-usage distance'. Distances of "diverged" ohnologs were then compared to "conserved" ohnologs.

Next, we sought evidence for a direct relationship between conservation/divergence in gene expression and regulatory landscape differences between ohnolog pairs. For this purpose, both enhancer-usage Euclidean distance and gene expression Euclidean distances were calculated as described above for all 10,366 ohnolog pairs, separately for DevMap and BodyMap samples. SOM cluster identifications, assigned to genes based on gene co-expression maps, were used to subset ohnolog groups into specific developmental stages or tissue types. At least one gene of the ohnolog pair needed to be assigned to a specific stage or tissue. Ohnolog pairs were allowed to have two assignments if each gene belonged to difference SOM clusters. Finally, ohnolog pair expression distance was plotted against enhancer-usage distance separately by developmental stage or tissue type, with density levels visualizing clusters of ohnolog pairs showing the relationships between active enhancer-usage and expression divergence. We used a global Kruskal-Wallis rank sum test to compare overall differences in dispersion of expression and enhancer distances between sample types. To test if ohnologs clustered around a single or multiple peaks (representing regulatory conservation or divergence), we used a two-component mixture model (Mclust R package (*103*)) to split ohnologs into two clusters and compared between sample types the proportion of ohnolog pairs fitting into the first or second cluster.

Finally, we compared TE content in active enhancer sequences of "conserved" and "diverged" ohnologs. The proportions TE content, calculated as base pairs of TE regions overlapping active enhancers regions, out of total enhancers base pairs, was determined per ohnolog pair separately by enhancer categorization (Shared, Alignable, Exclusive). TE content in exclusive enhancers was then broken down by four major TE orders: DNA, LINE, LTR, and SINE, and specifically by the Tc1/mariner (TcMar) class of DNA transposons, the most abundant class of TE in the Atlantic salmon genome. TE content for TcMar transposons were then broken down by sample types i.e., different developmental stages and tissues. Means of values per ohnolog pairs were used for bootstrapping 95% confidence intervals and for Wilcoxon tests between "conserved" and "diverged" ohnologs.

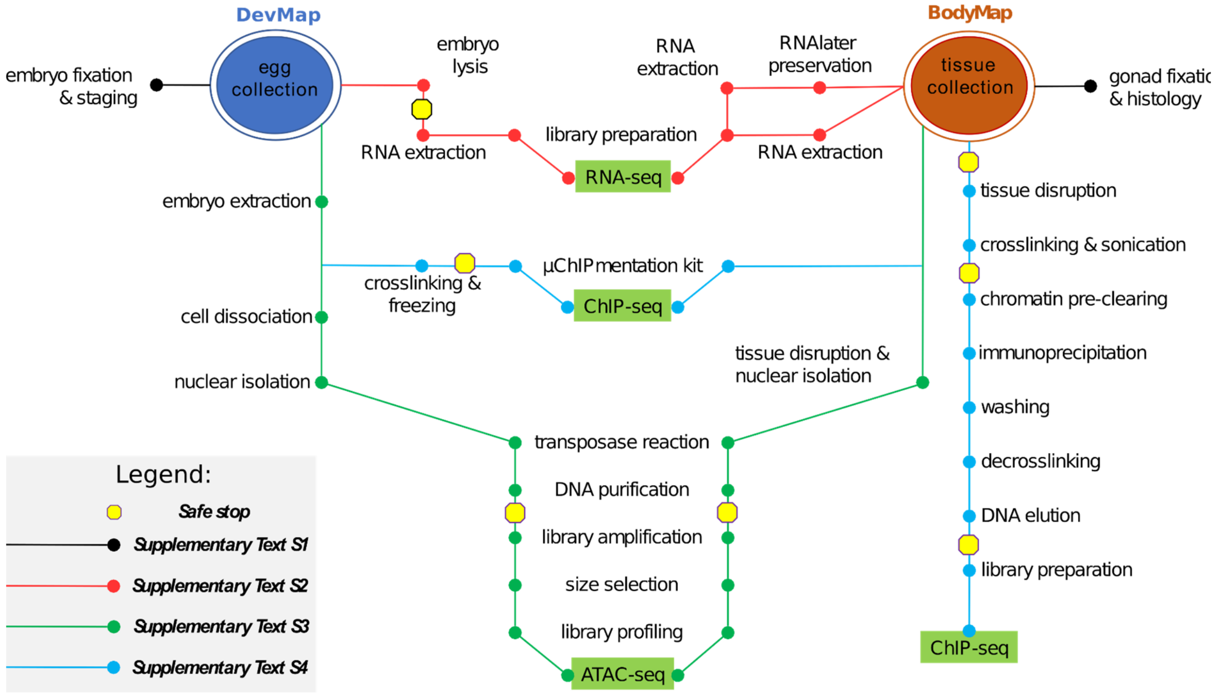

**Fig. S1. Graphic visualization of protocol development for sequencing assays used to functionally annotate two salmonid genomes**. The map illustrates interconnections between different protocols fully outlined in Text S1-S4.

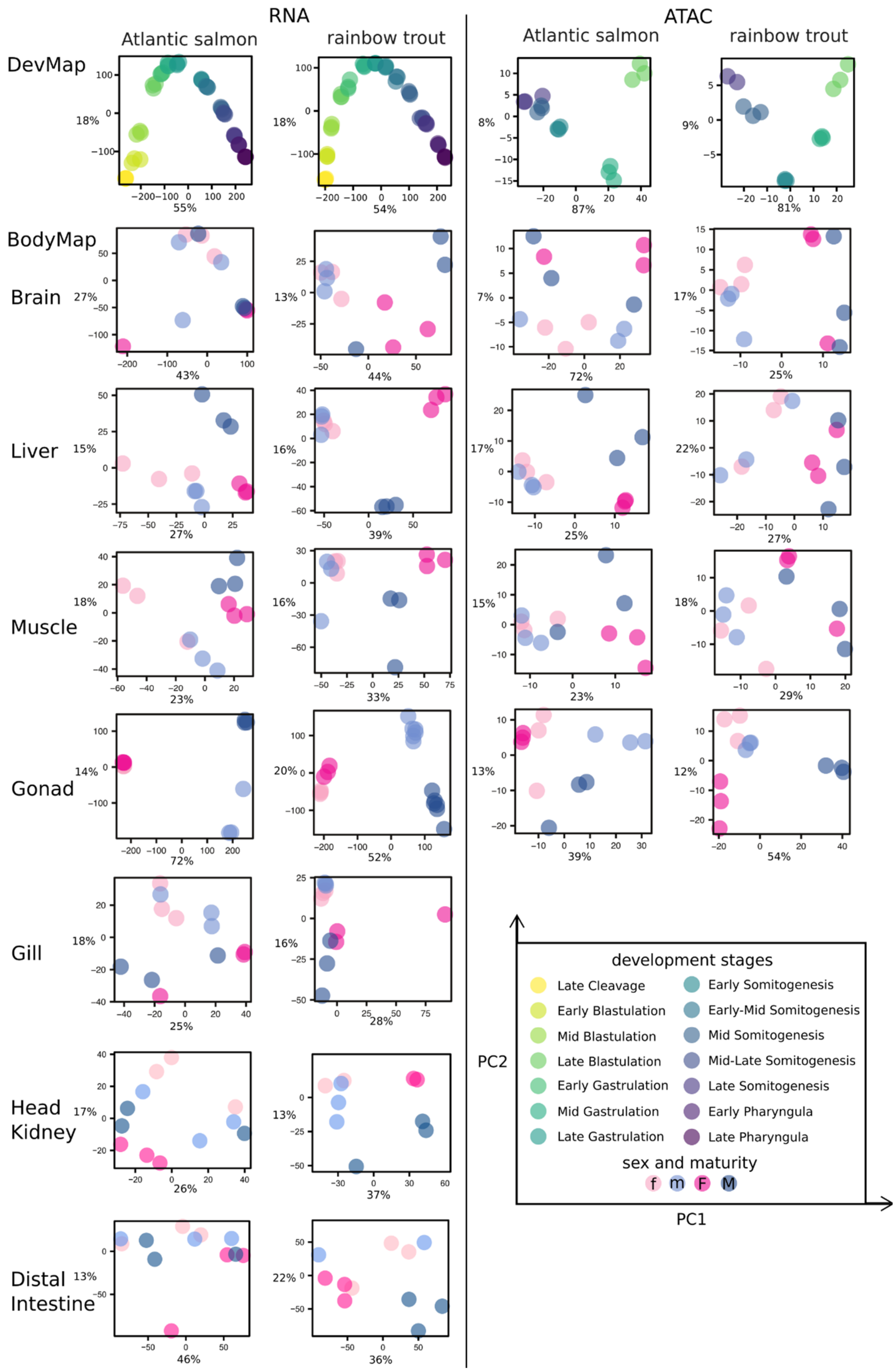

**Fig. S2. Principal component analysis (PCA) of RNA-Seq and ATAC-seq datasets for DevMap and BodyMap from Atlantic salmon and rainbow trout**. The key at the bottom right highlights different DevMap stages or symbols used for sexes in BodyMap (tissues separated into immature = lowercase ‘f’ or ‘m’ and mature = uppercase ‘F’ and ‘M’). The proportion of variation explained by PC1 and PC2 is shown on each plot.

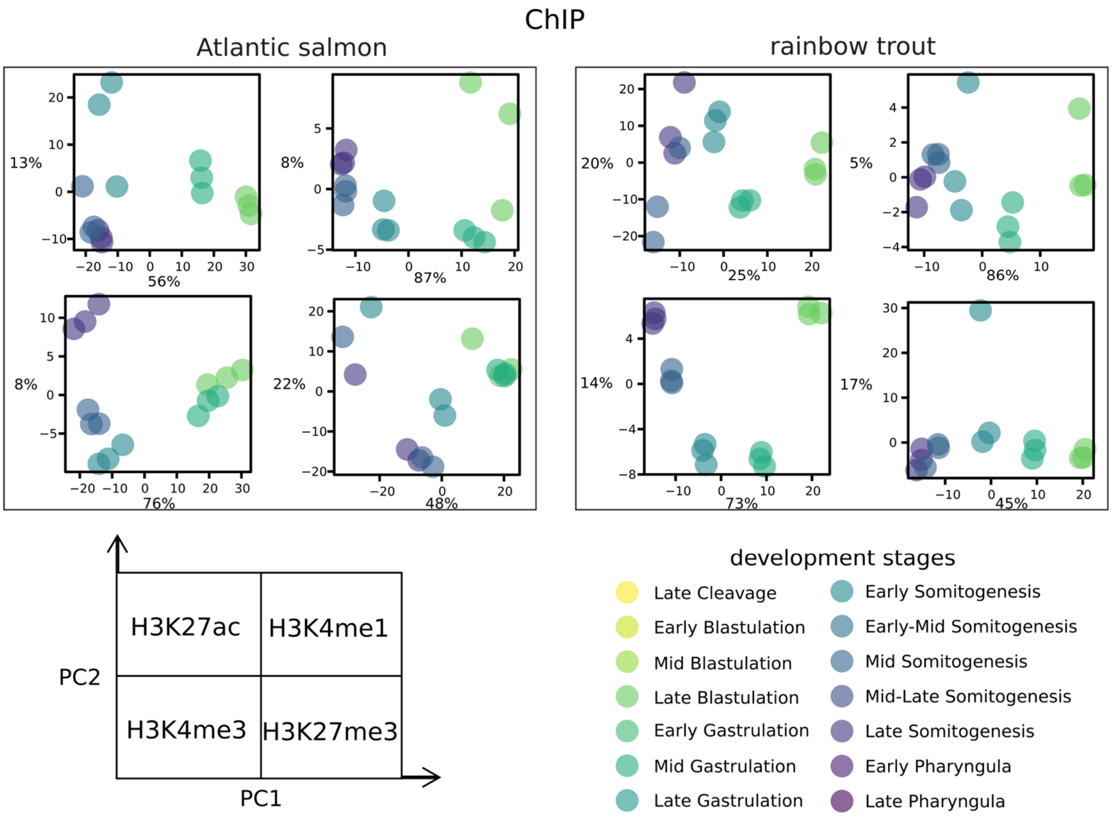

**Fig. S3. PCA of ChIP-seq datasets for DevMap from Atlantic salmon and rainbow trout**. The key at the bottom right highlights different DevMap stages. The proportion of variation explained by PC1 and PC2 is shown on each plot.

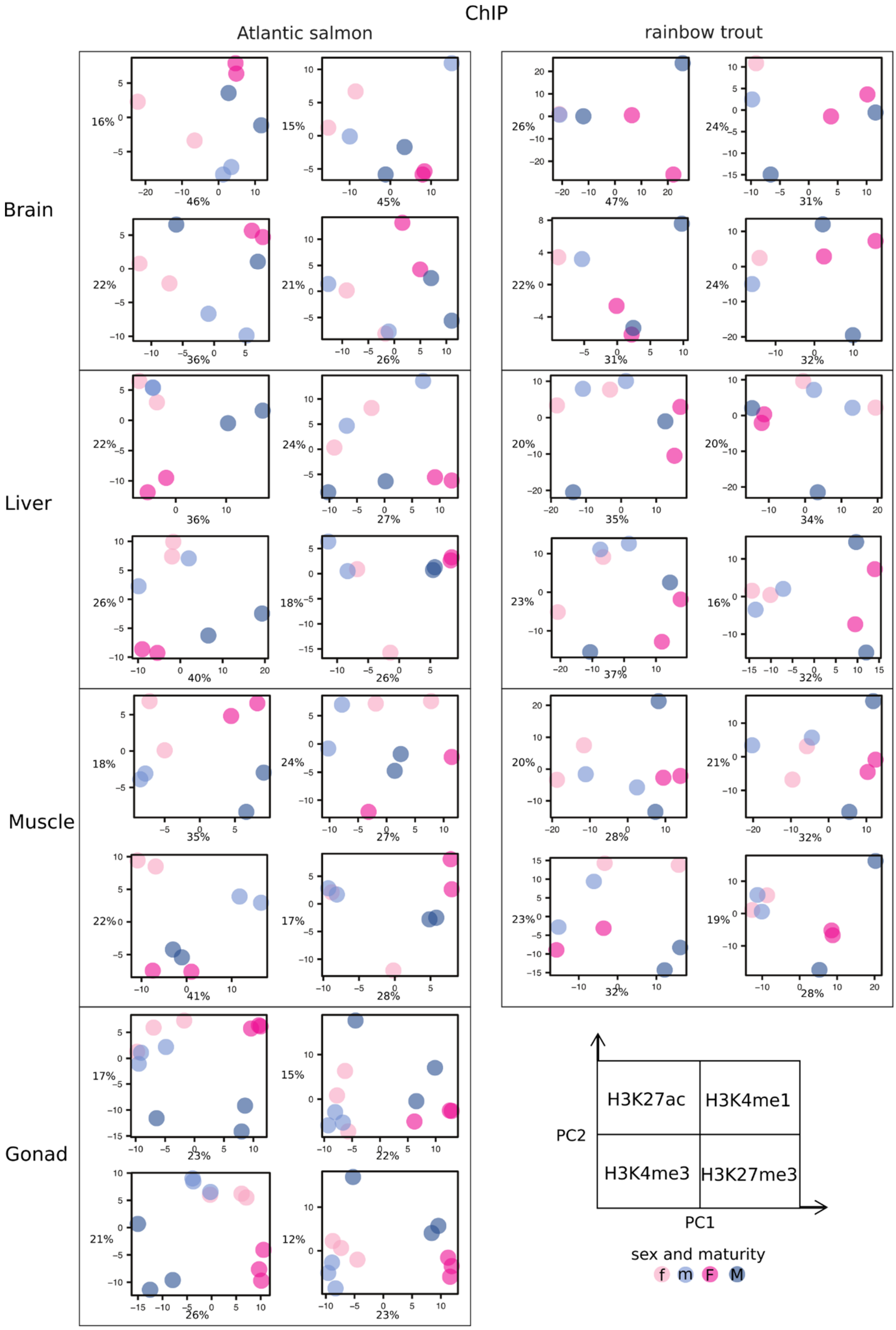

**Fig. S4. PCA of ChIP-seq datasets for BodyMap from Atlantic salmon and rainbow trout.** The key at the bottom right highlights different symbols used for sexes in BodyMap (tissues separated into immature = lowercase ‘f’ or ‘m’ and mature = uppercase ‘F’ and ‘M’). The proportion of variation explained by PC1 and PC2 is shown on each plot.

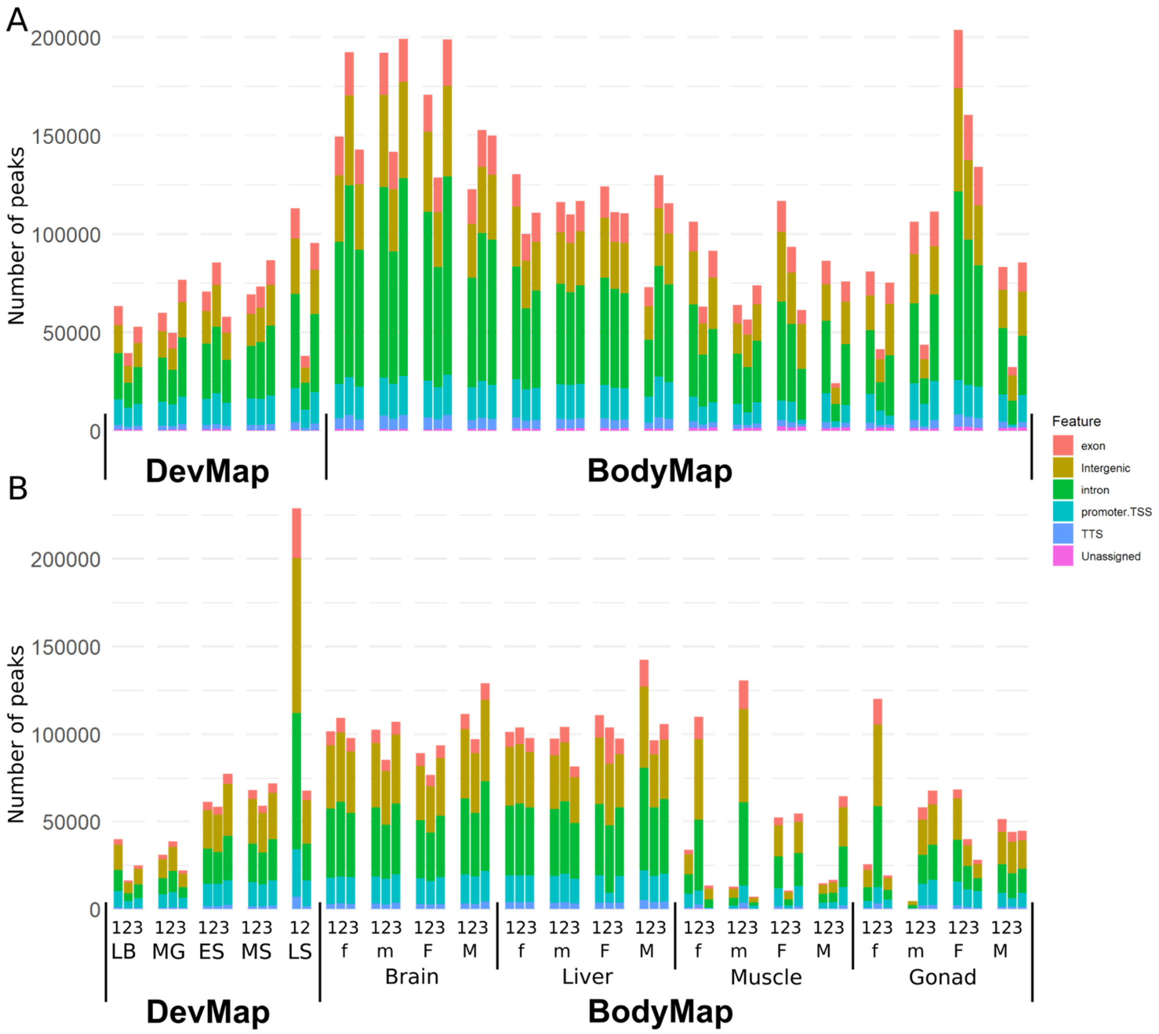

**Fig. S5. Overlap of robust open chromatin regions with annotated genome features.** Data are shown for DevMap (stages: LB = late blastulation; MG = mid gastrulation; ES = early somitogenesis; MS = mid-somitogenesis; LS = late somitogenesis) and BodyMap (tissues separated into immature = lowercase 'm' or 'f' and mature = uppercase 'M' and 'F') samples for females and males, respectively) for Atlantic salmon (**A**) and rainbow trout (**B**).

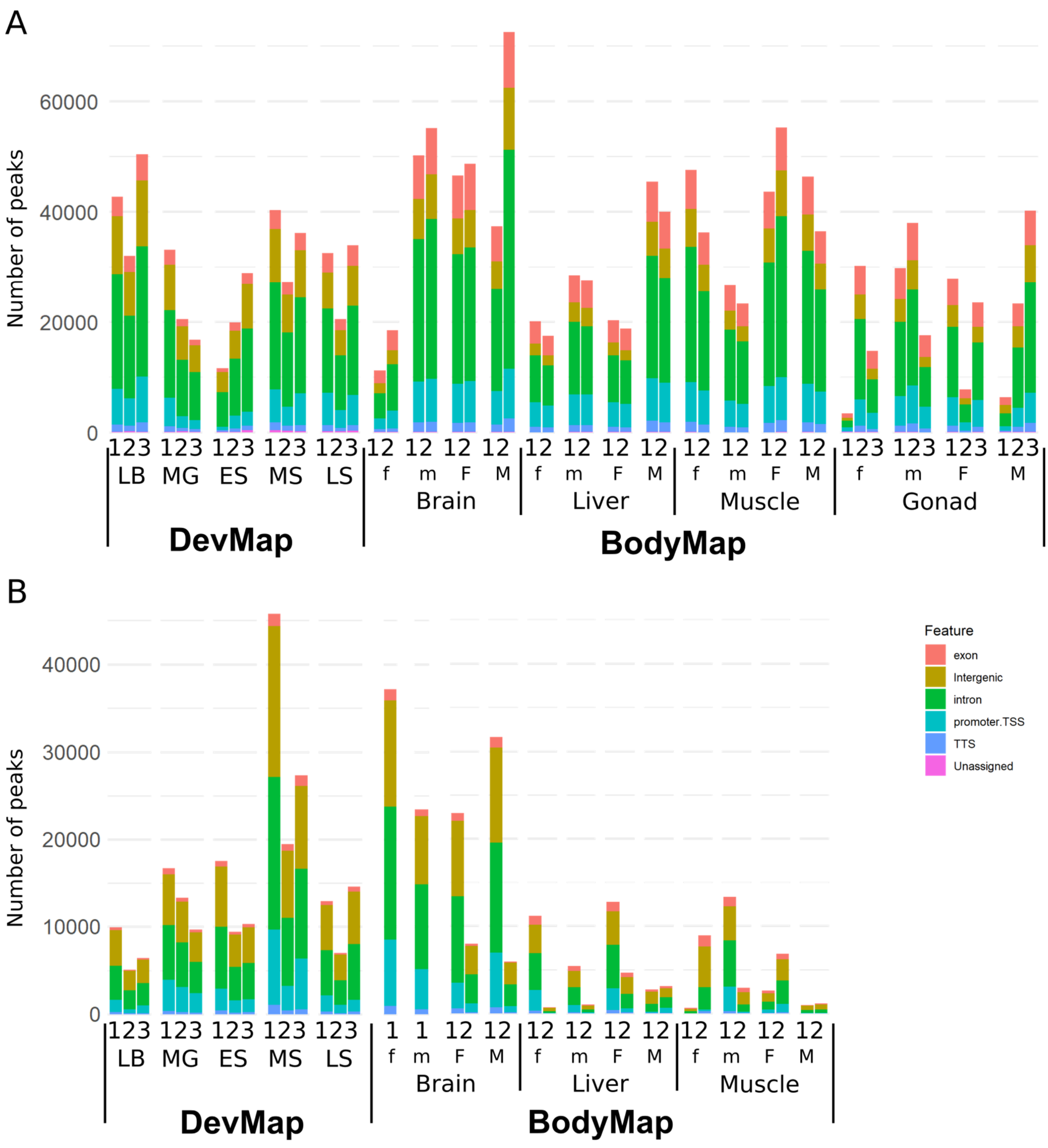

**Fig. S6. Overlap of ChIP-seq peaks with annotated genome features for H3K27ac.** Data are shown for both DevMap (stages: LB = late blastulation; MG = mid gastrulation; ES = early somitogenesis; MS = mid-somitogenesis; LS = late somitogenesis) and BodyMap (tissues separated into immature = lowercase 'f' or 'm' and mature = uppercase 'F' and 'M') samples for females and males, respectively) for Atlantic salmon (**A**) and rainbow trout (**B**).

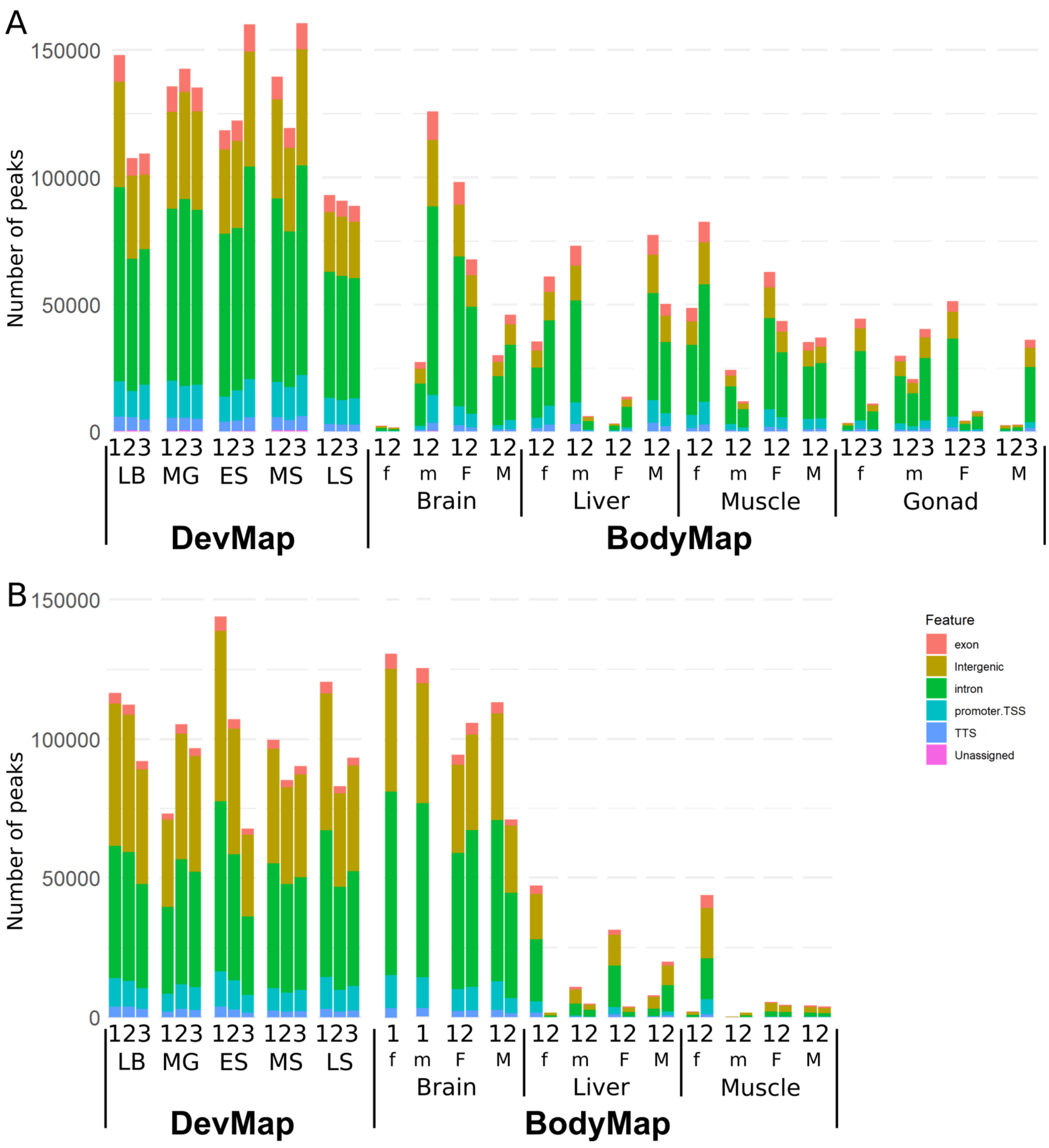

**Fig. S7. Overlap of ChIP-seq peaks with annotated genome features for H3K4me1.** Data are shown for DevMap (stages: LB = late blastulation; MG = mid gastrulation; ES = early somitogenesis; MS = mid-somitogenesis; LS = late somitogenesis) and BodyMap (tissues separated into immature = lowercase 'f' or 'm' and mature = uppercase 'F' and 'M') samples for females and males, respectively) for Atlantic salmon (**A**) and rainbow trout (**B**).

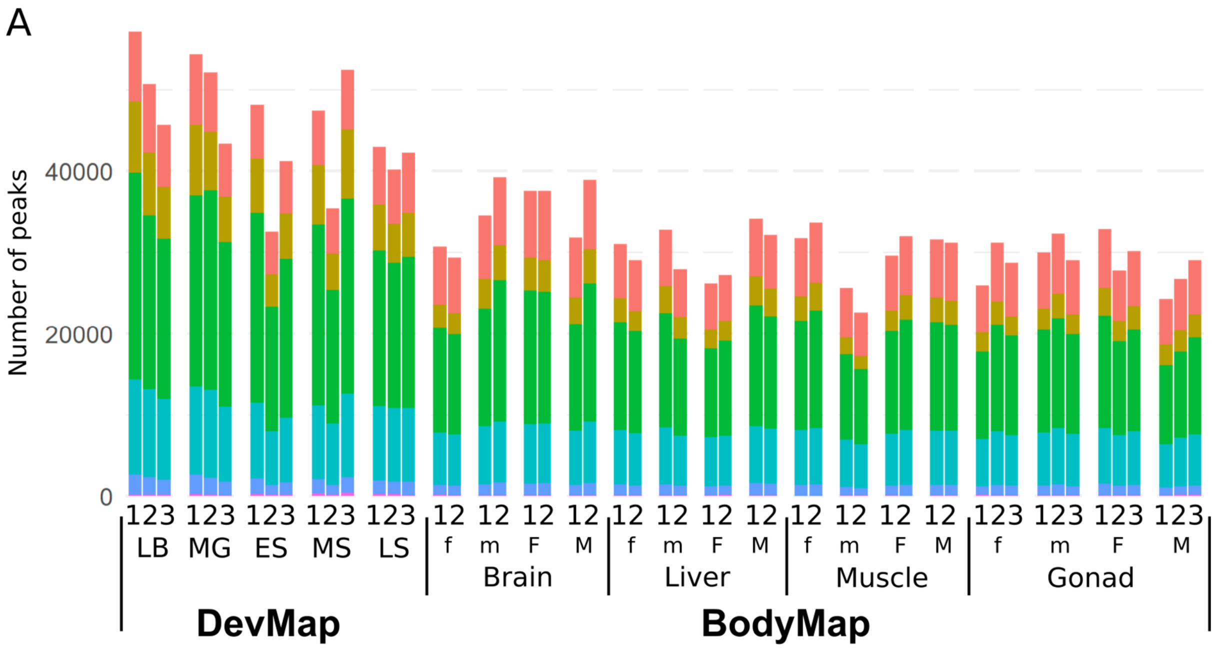

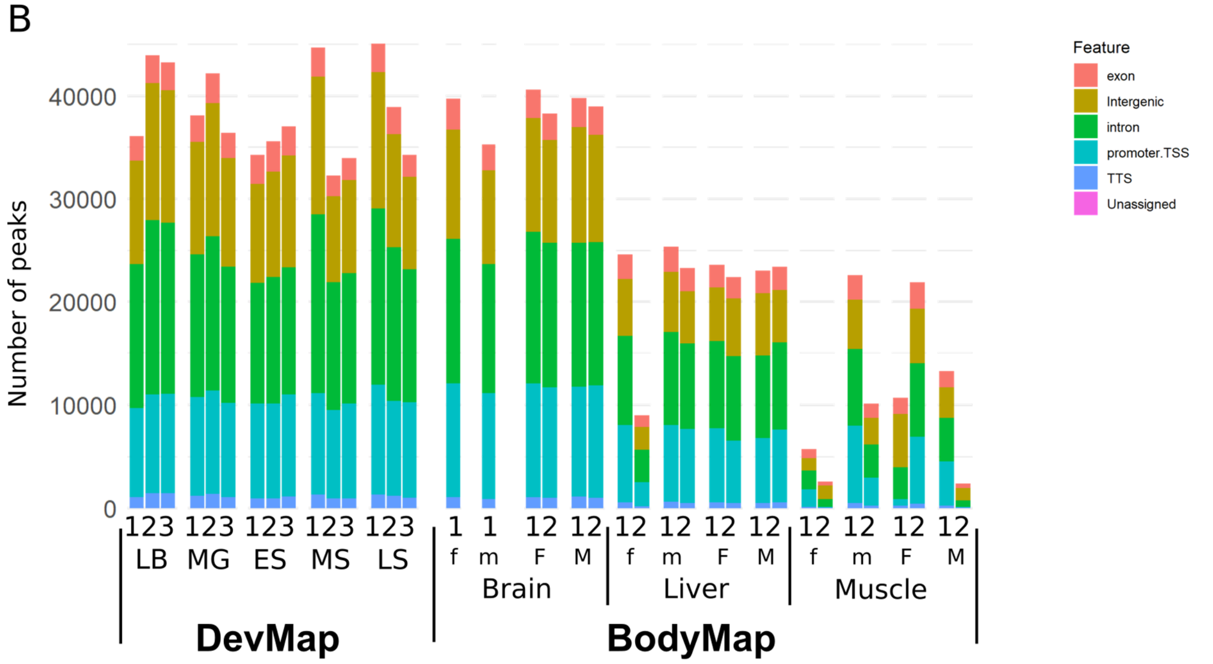

**Fig. S8. Overlap of ChIP-seq peaks with annotated genome features for H3K4me3.** Data are shown for DevMap (stages: LB = late blastulation; MG = mid gastrulation; ES = early somitogenesis; MS = mid-somitogenesis; LS = late somitogenesis) and BodyMap (tissues separated into immature = lowercase 'f' or 'm' and mature = uppercase 'F' and 'M') samples for females and males, respectively) for Atlantic salmon (**A**) and rainbow trout (**B**).

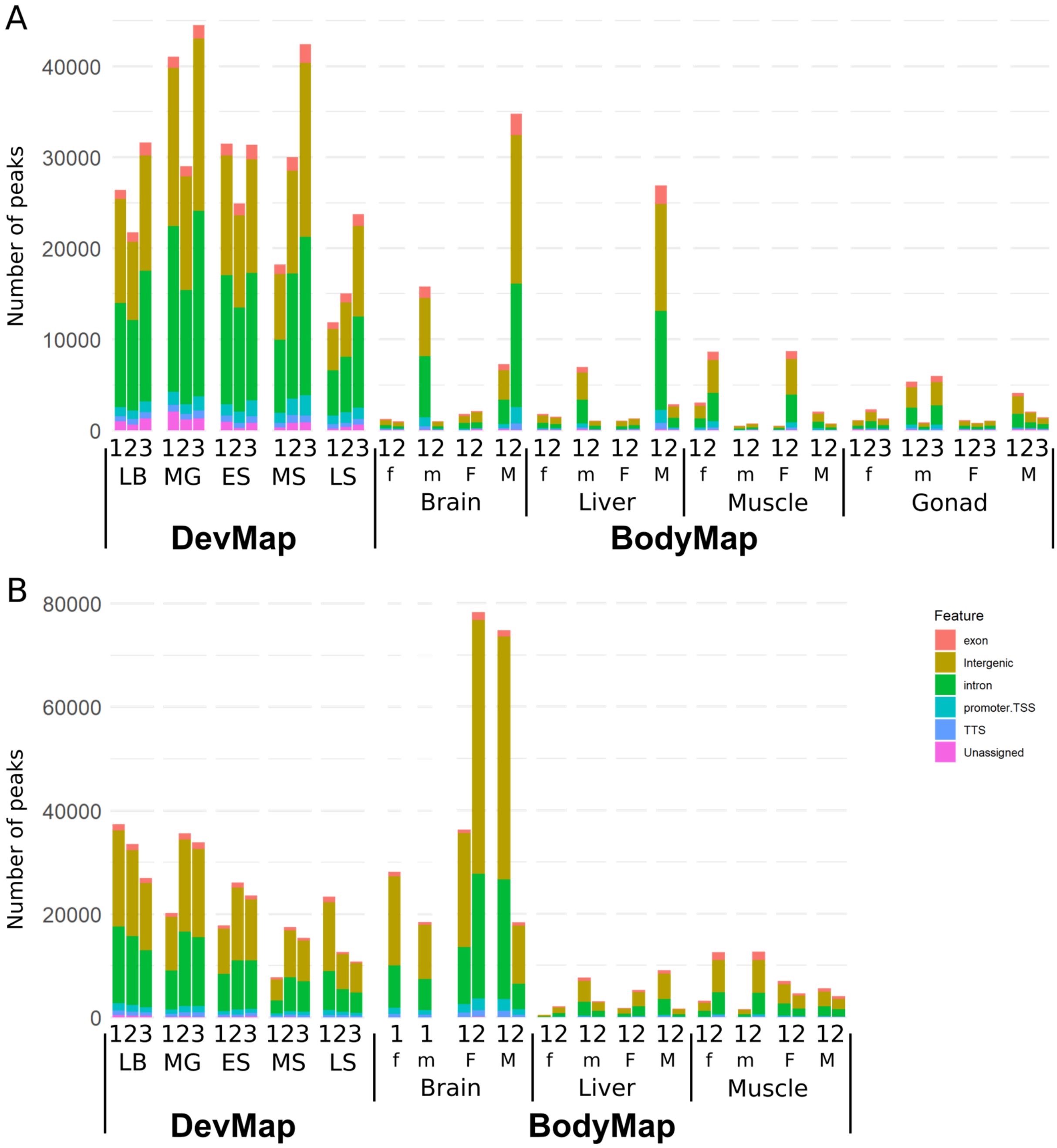

**Fig. S9. Overlap of ChIP-seq peaks with annotated genome features for H3K27me3.** Data are shown for DevMap (stages: LB = late blastulation; MG = mid gastrulation; ES = early somitogenesis; MS = mid-somitogenesis; LS = late somitogenesis) and BodyMap (tissues separated into immature = lowercase ‘f’ or ‘m’ and mature = uppercase ‘F’ and ‘M’) samples for females and males, respectively) for Atlantic salmon (**A**) and rainbow trout (**B**).

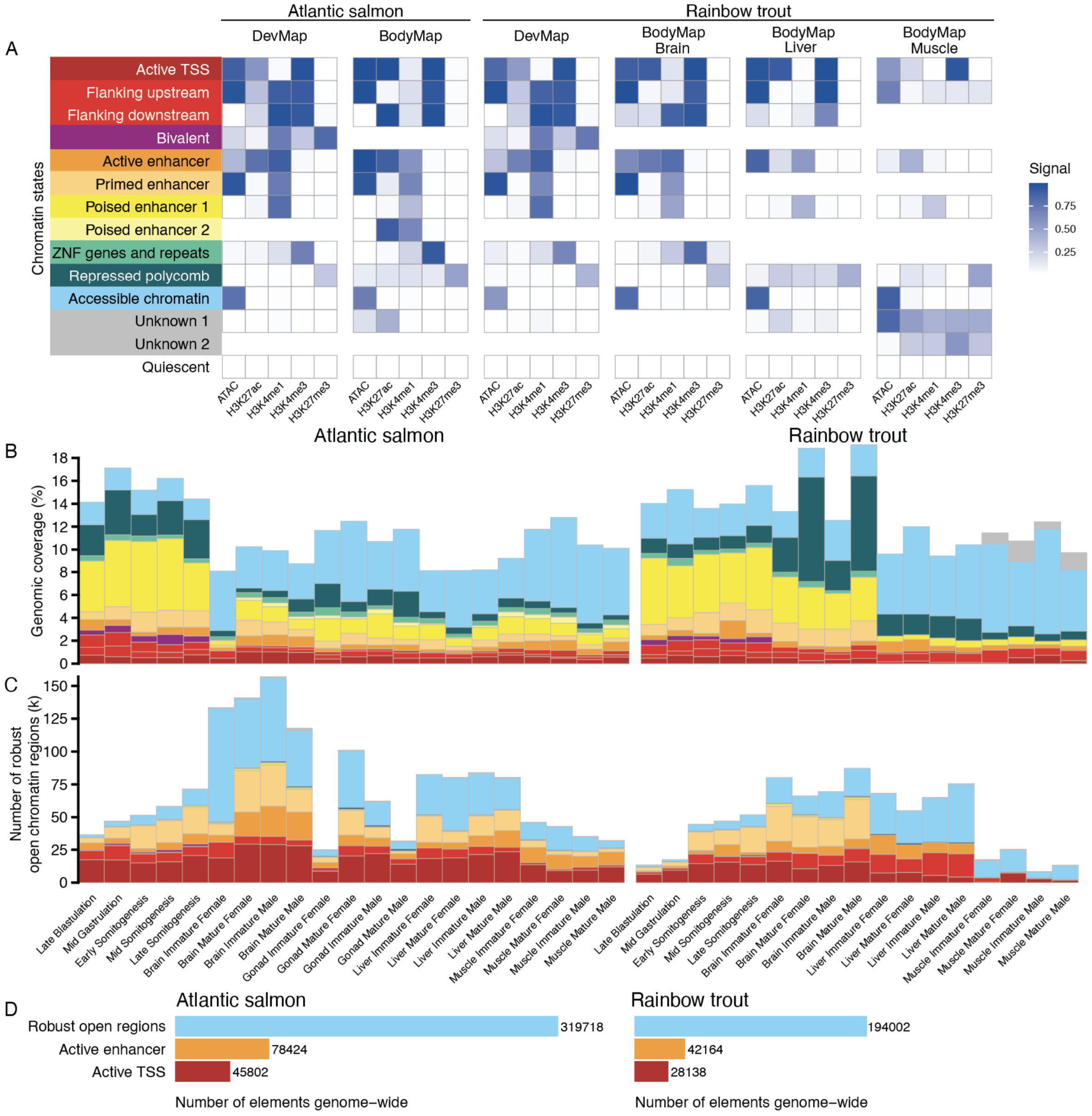

**Fig. S10. ChromHMM models and robust open chromatin regions defined across two salmonid genomes to predict regulatory elements**. (**A**) Comparison of selected ChromHMM chromatin state models for Atlantic salmon and rainbow trout DevMap and BodyMap datasets. (**B**) Sample-specific proportions (values shown are means of biological replicates) of each salmonid genome covered by different ChromHMM chromatin states, colored according to part **A**. (**C**) Sample-specific counts (values shown are means of biological replicates) of robust chromatin regions representing different chromatin states for each salmonid species. (**D**) Counts of genome-wide robust open chromatin regions, including those comprising active enhancer and active promoters (i.e., active TSS) for Atlantic salmon and rainbow trout, combined across DevMap and BodyMap.

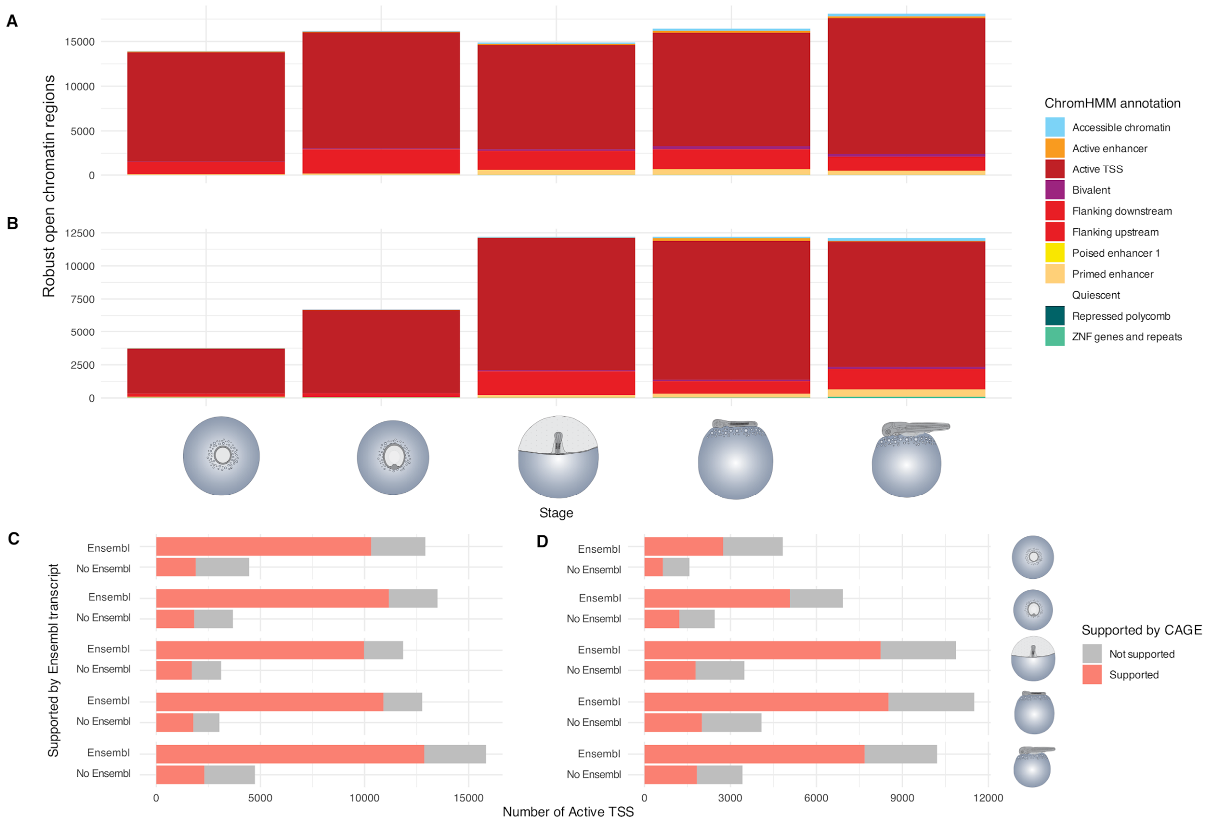

**Fig. S11. Validation of annotated regulatory elements using CAGE data with DevMap samples**. (**A, B**) Overlap between ChromHMM annotated regulatory elements and CAGE promoters in Atlantic salmon (**A**) and rainbow trout (**B**). The majority of overlapping peaks are associated with promoter or promoter-proximal functions, confirming the specificity and accuracy of ChromHMM annotations when independently validated with CAGE. (**C, D**) Bar charts depicting the number of regulatory elements with Ensembl transcript annotation support and without Ensembl transcript annotation support, colored by their CAGE support status, in Atlantic salmon (**C**) and rainbow trout (D). While most Ensembl-supported elements are also supported by CAGE data, a subset of elements without Ensembl support show CAGE support, indicating the potential presence of novel, previously unidentified transcripts.

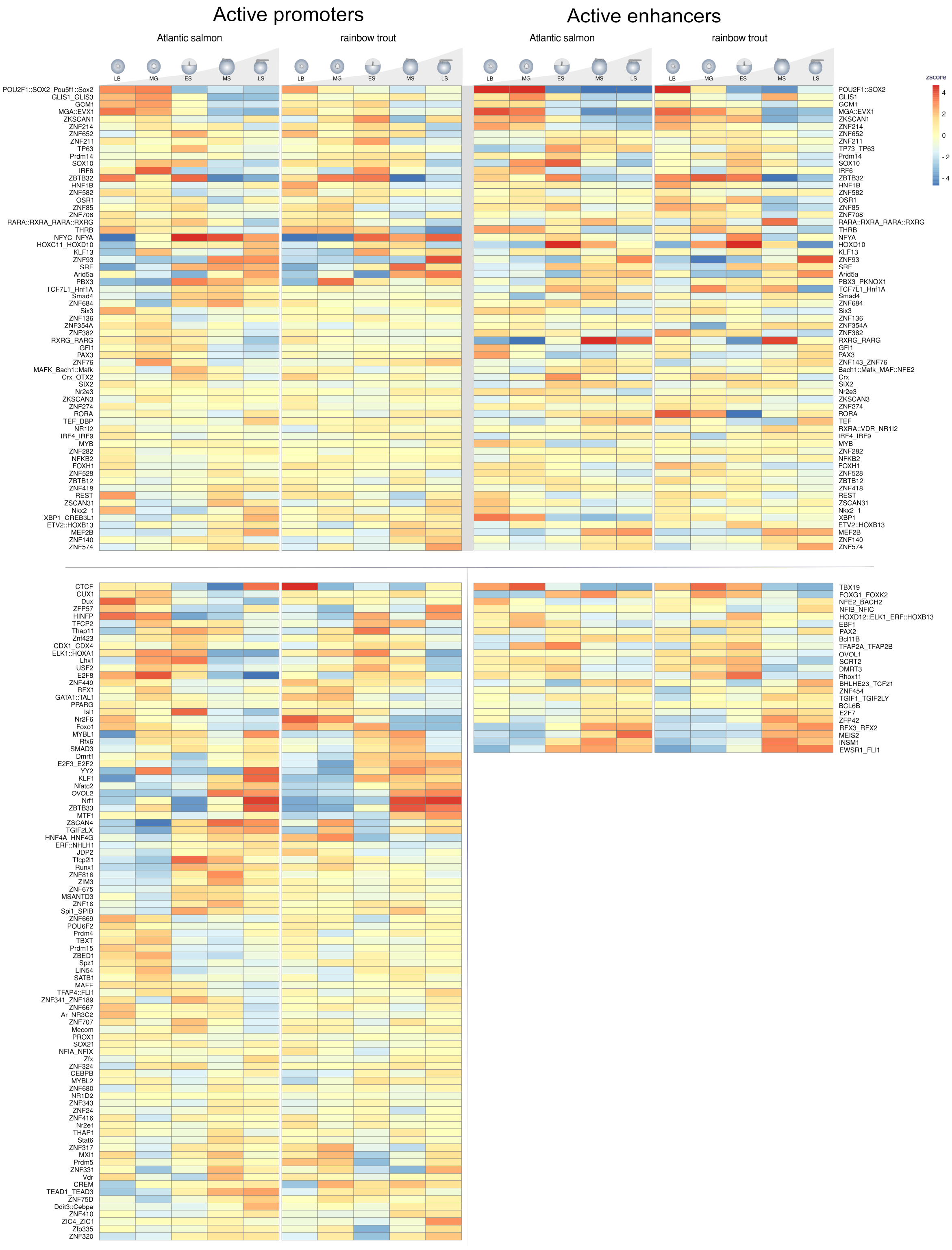

**Fig. S12**. **Transcription factor binding sites (TFBS) enrichment analysis across DevMap stages in Atlantic salmon and rainbow trout**. Enrichment of TFBSs within robust open chromatin regions annotated by ChromHMM as active promoters (**A**) or enhancers (**B**). TFBSs showing significant differences across stages are shown on a heatmap, with colours representing z-score values to visualize relative overrepresentation or underrepresentation according to the key on the right side. Text S5 includes a summary of key results.

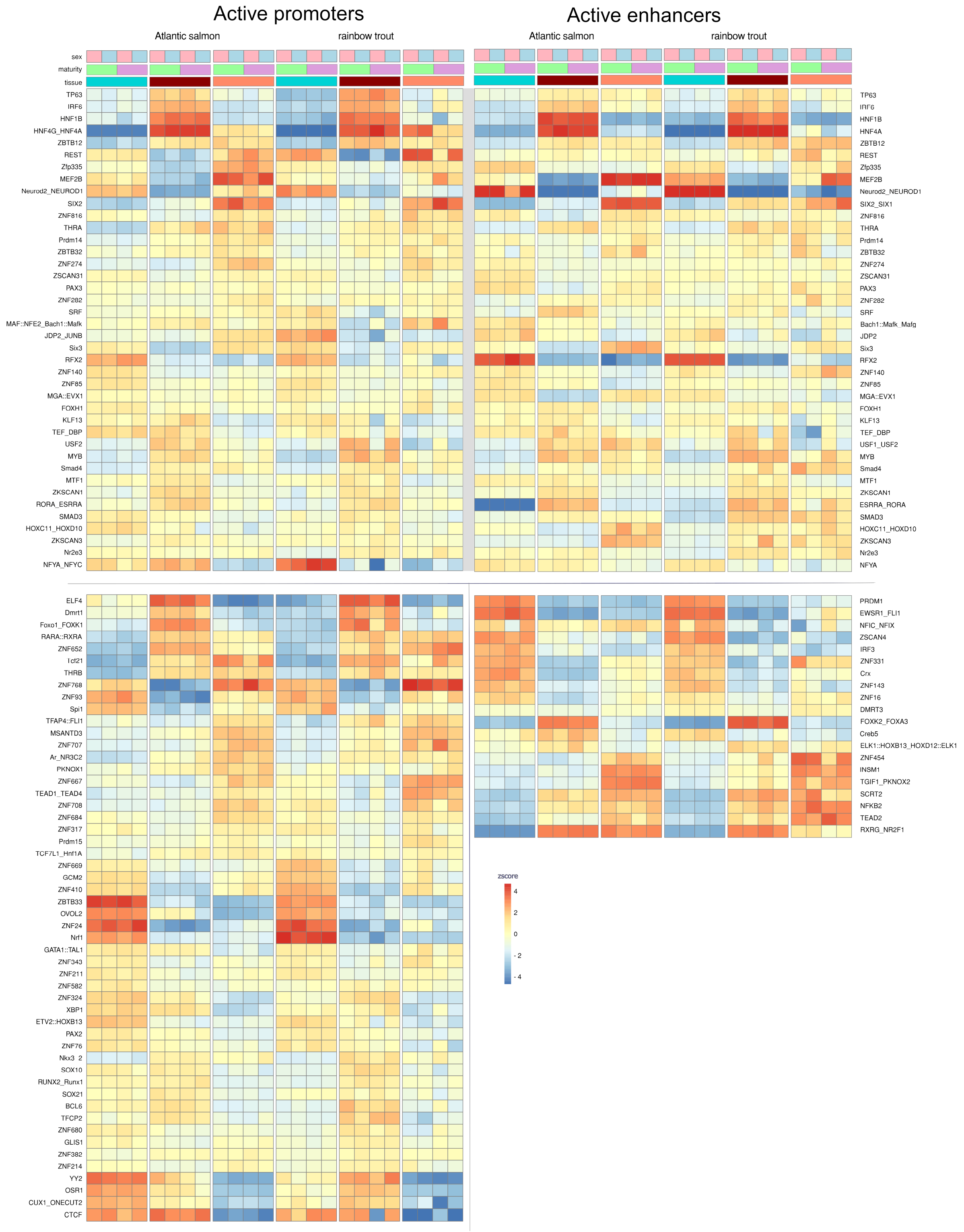

**Fig. S13**. TFBS enrichment analysis across BodyMap tissues in Atlantic salmon and rainbow trout. Enrichment of TFBSs within robust open chromatin regions annotated by ChromHMM as active promoters (**A**) or enhancers (**B**). TFBSs showing significant differences across stages are shown on a heatmap, with colours representing z-score values to visualize relative overrepresentation or underrepresentation according to the key on the right side. Text S5 includes a summary of key results.

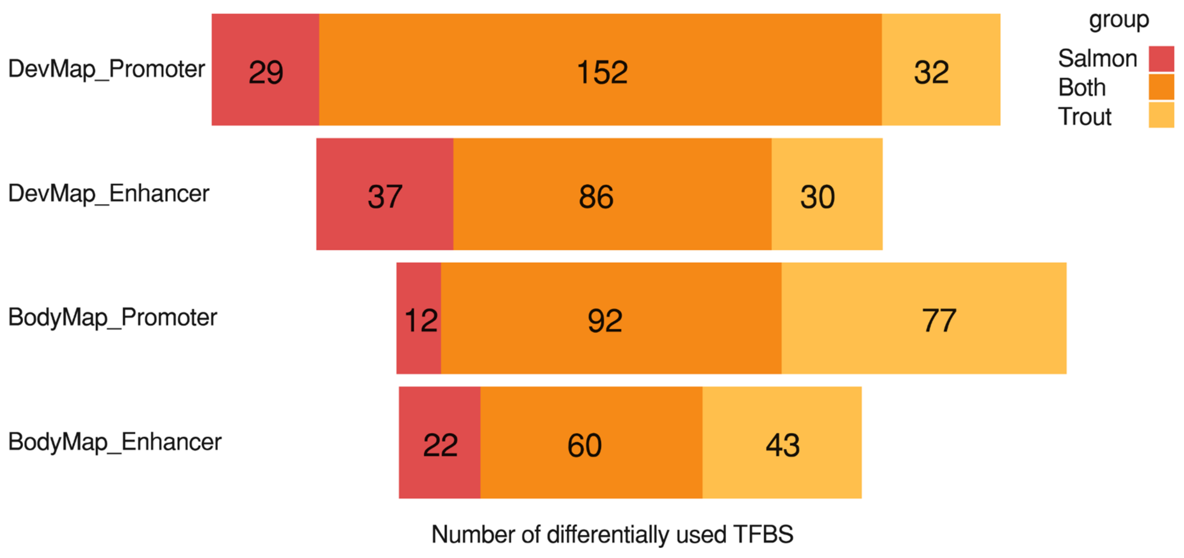

**Fig. S14. Overlap between enriched TFBS for Atlantic salmon and rainbow trout in DevMap and BodyMap sample types**. For DevMap or BodyMap, the number of overlapping TFBS found enriched within robust open chromatin regions annotated by ChromHMM as active TSS or active enhancers for both species is presented in orange, while the number of unique TFBS is depicted in red for Atlantic salmon and in yellow for rainbow trout. Text S5 includes a summary of key results.

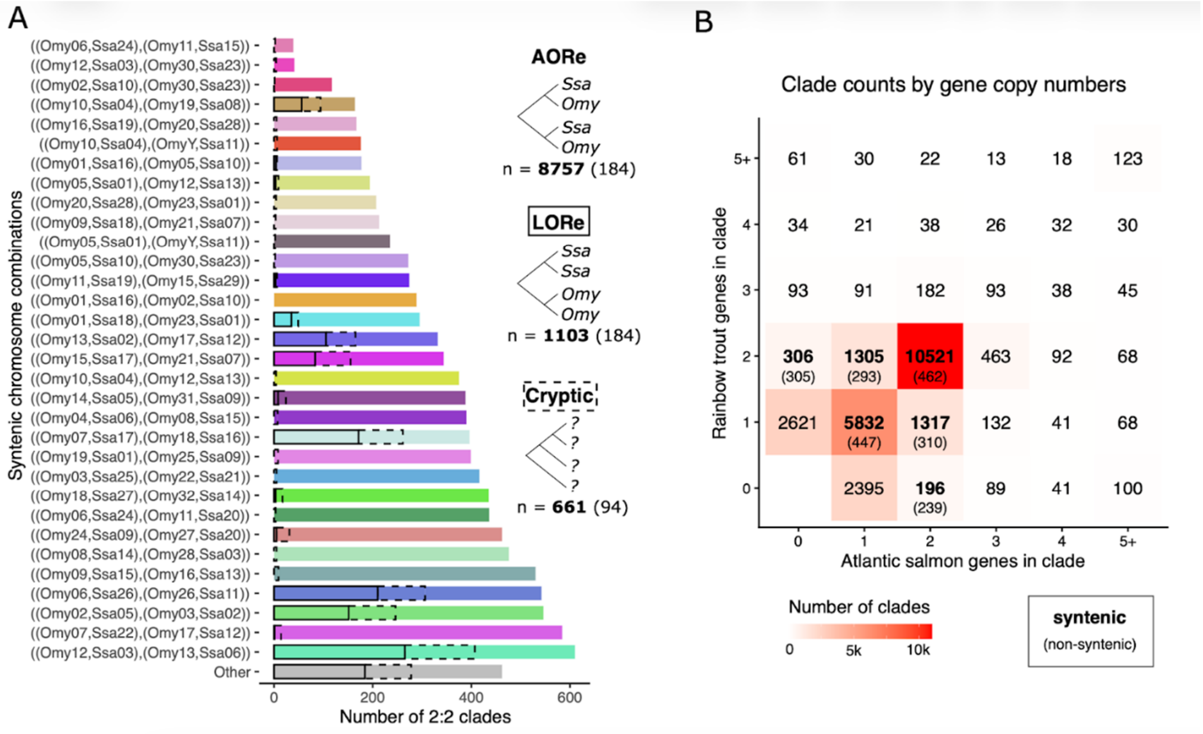

**Fig. S15. Homology prediction capturing ohnologs (within) and orthologs (between) for the Atlantic salmon and rainbow trout genomes** (see Materials and Methods). (**A**) Identification of homologous/syntenic chromosome regions represented by genes captured within 2:2 clades in phylogenetic analysis (2:2 clades are those where both salmonid species retain two duplicated genes). Inserts show the number of AORe (i.e., Early rediploidization; ohnolog divergence shared by both salmonid species), LORe (lineage-specific rediploidization; ohnolog divergence post-speciation of Atlantic salmon and rainbow trout; also referred to in this study as Late rediploidization) and other (cryptic) phylogenetic topologies. Numbers in parenthesis represent clades not matching syntenic chromosomes, i.e "Other". (**B**) Number of clades capturing different numbers of genes in the two salmonid species. The bold numbers show how many of the clades capture genes located within syntenic chromosomes, as expected by a WGD event, i.e., representing our definition of high-confidence ohnologs (numbers in parenthesis represent clades not in syntenic chromosomes). The plot highlights the dominance of 2:2 clades (i.e., ohnolog pairs retained in both species), 1:1 clades (e.g., resulting from one ohnolog in a pair being lost ancestrally to Atlantic salmon and rainbow trout), as well as 2:1 clades (e.g., where one ohnolog was lost in one species, and the other species retains both ohnologs).

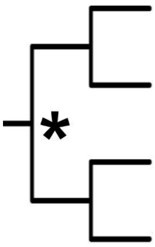

| Ssa03:53,495,240 – 100,966,100 | | | Ssa03:2,160,013 – 45,141,280 | | |
|---|---|---|---|---|---|
| LORe | Ensembl | AQUA-FAANG | AORe | Ensembl | AQUA-FAANG |
| *Ssa03* | 47,623,004 | 47,470,893 | *Ssa03* | 43,083,082 | 42,981,345 |
| *Ssa06* | 26,638,611 | 25,604,383 | *Omyk28* | 26,638,736 | 26,837,948 |
| *Omyk12* | 23,530,163 | 26,143,366 | *Ssa14* | 9,597,080 | 15,716,672 |
| *Omyk13* | 23,137,724 | 25,833,768 | *Omyk08* | 9,204,883 | 14,845,254 |

**Fig. S16**. **Comparison of effectiveness of two whole genome alignment strategies for capture of duplicated regions from salmonid WGD**. The table shows data derived from the alignment of a chromosome selected from the Atlantic salmon genome that contains two distinct rediploidization histories (*17*, *18*) including a large LORe (Late rediploidization) region (Ssa03-06) that underwent rediploidization after the split of *Salmo* and *Oncorhynchus* and a large AORe (Early rediploidization) region that underwent rediploidization early in salmonid evolution (Ssa03-14). We compared the capture of these regions, along with their homologous regions in rainbow trout (Omyk12-13 and Omyk28-08) in two Cactus alignments (see Methods). The first was an alignment that was prepared by the Ensembl Genome Browser ("Ensembl") run using a standard approach that is not aware of WGD, and the second the alignment prepared in this study ("AQUA-FAANG") with our WGD-aware approach. The WGD-aware approach recovered around 37-38% more aligned sequence in AORe regions for both species, and 10-11% in LORe (Late rediploidization) regions.

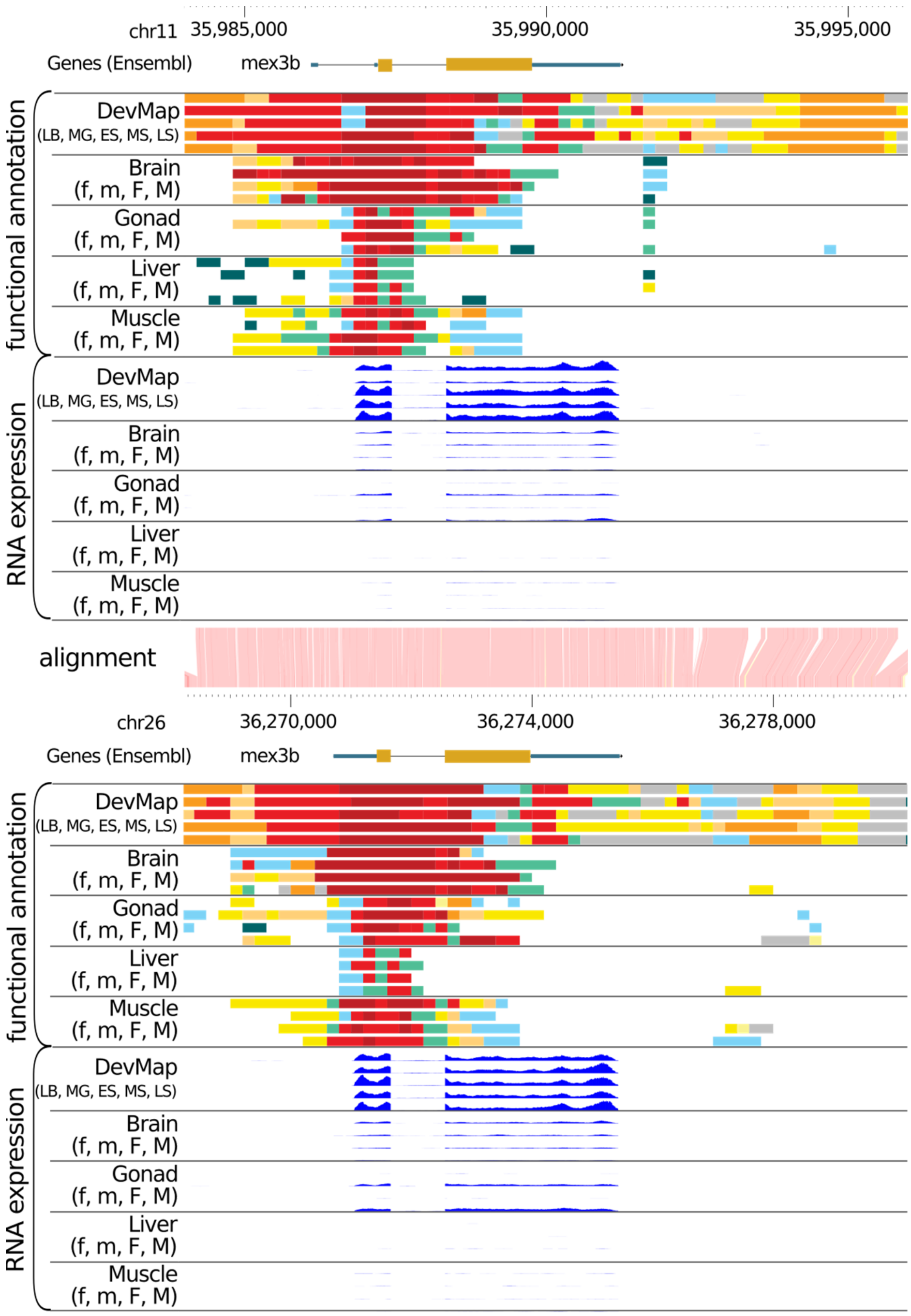

**Fig. S17. Integration of functional annotation data for a pair of rainbow trout ohnologs for *mex3b*.** The genes shown are found on Chromosome 6 (ENSOMYG00000040028) and 26 (ENSOMYG00000077707). Predicted chromatin states (colours as per Fig. S10) and corresponding gene expression is visualized for both DevMap and BodyMap annotations. Equivalent figure for Atlantic salmon is provided in Fig. 1.

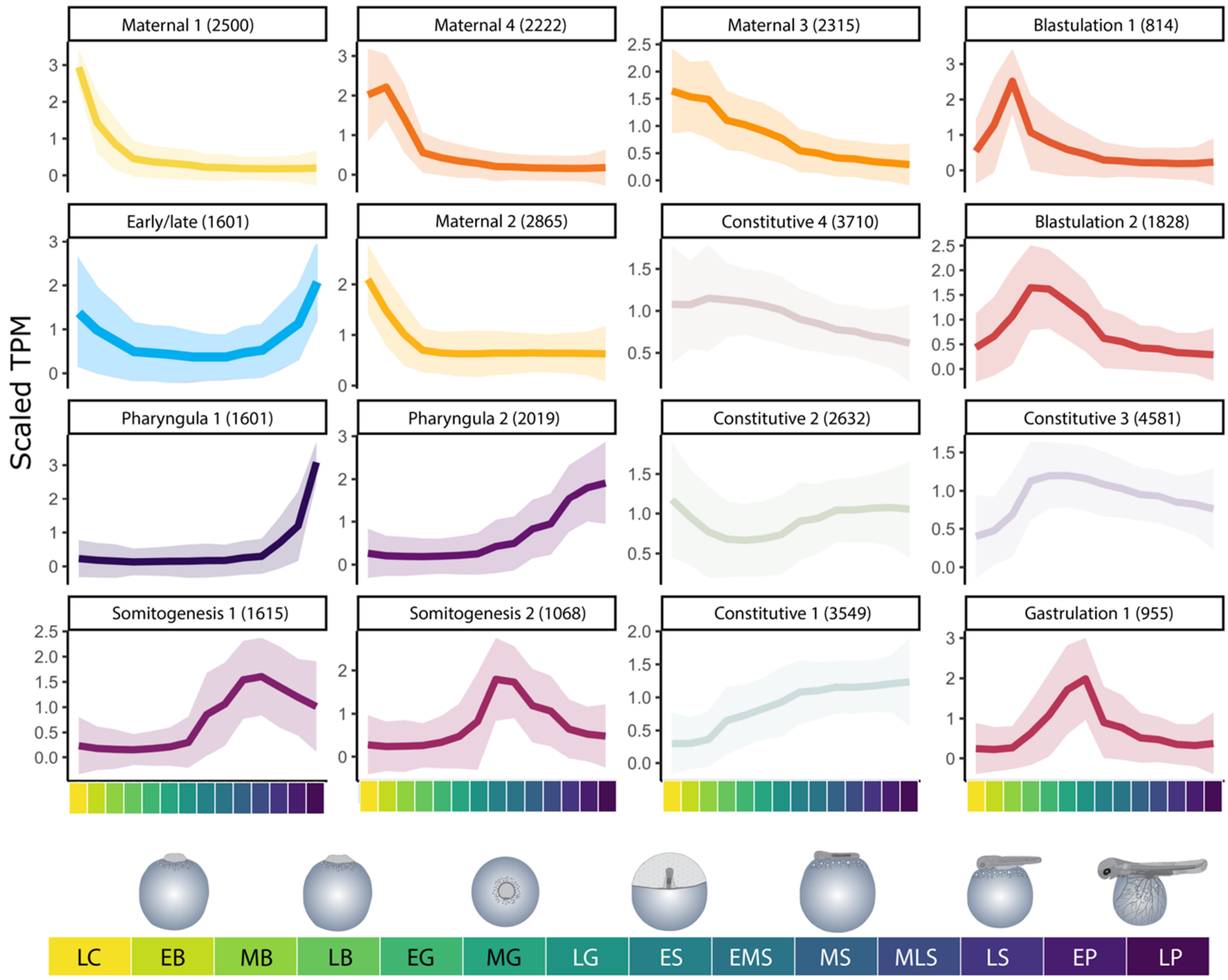

**Fig. S18. Atlantic salmon DevMap transcriptome SOM clusters**. Each individual line plot represents one SOM cluster and shows average normalised gene expression, with the position on the panel assigned by the SOM algorithm based on the inferred topology of the data. The colours indicate the order of expression throughout embryogenesis, based on colours used in Fig. 2 in the main text. Colored areas around each line plot show 95% confidence intervals. The top of each line plot indicates the name assigned to each SOM cluster based on gene expression pattern, along with the number of genes captured per cluster. Stages are indicated using the colours shown in the bottom annotation.

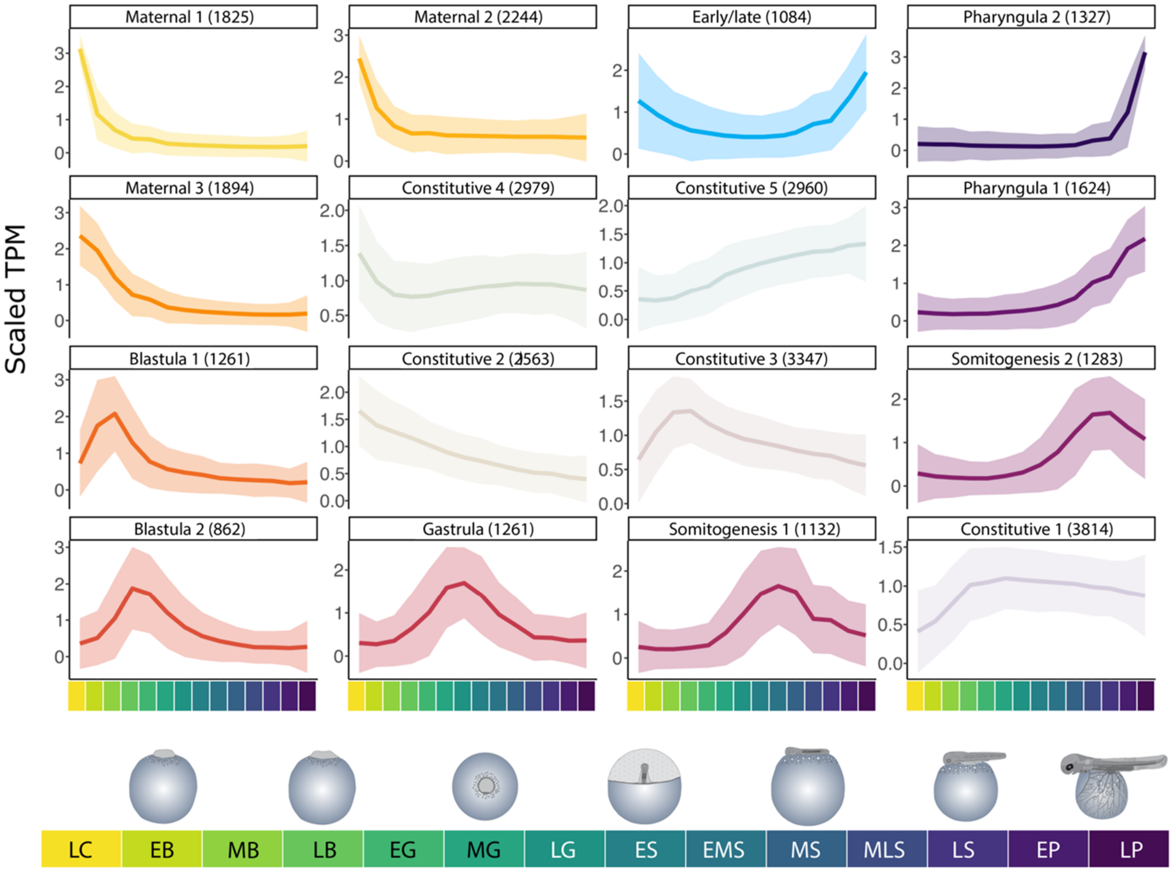

**Fig. S19. Rainbow trout DevMap transcriptome SOM clusters**. Each individual line plot represents one SOM cluster and shows average normalised gene expression, with the position on the panel assigned by the SOM algorithm based on the inferred topology of the data. The colours indicate the order of expression throughout embryogenesis, based on colours used in Fig. 2 in the main text. Colored areas around each line plot show 95% confidence intervals. The top of each line plot indicates the name assigned to each SOM cluster based on gene expression pattern, along with the number of genes captured per cluster. Stages are indicated using the colours shown in the bottom annotation.

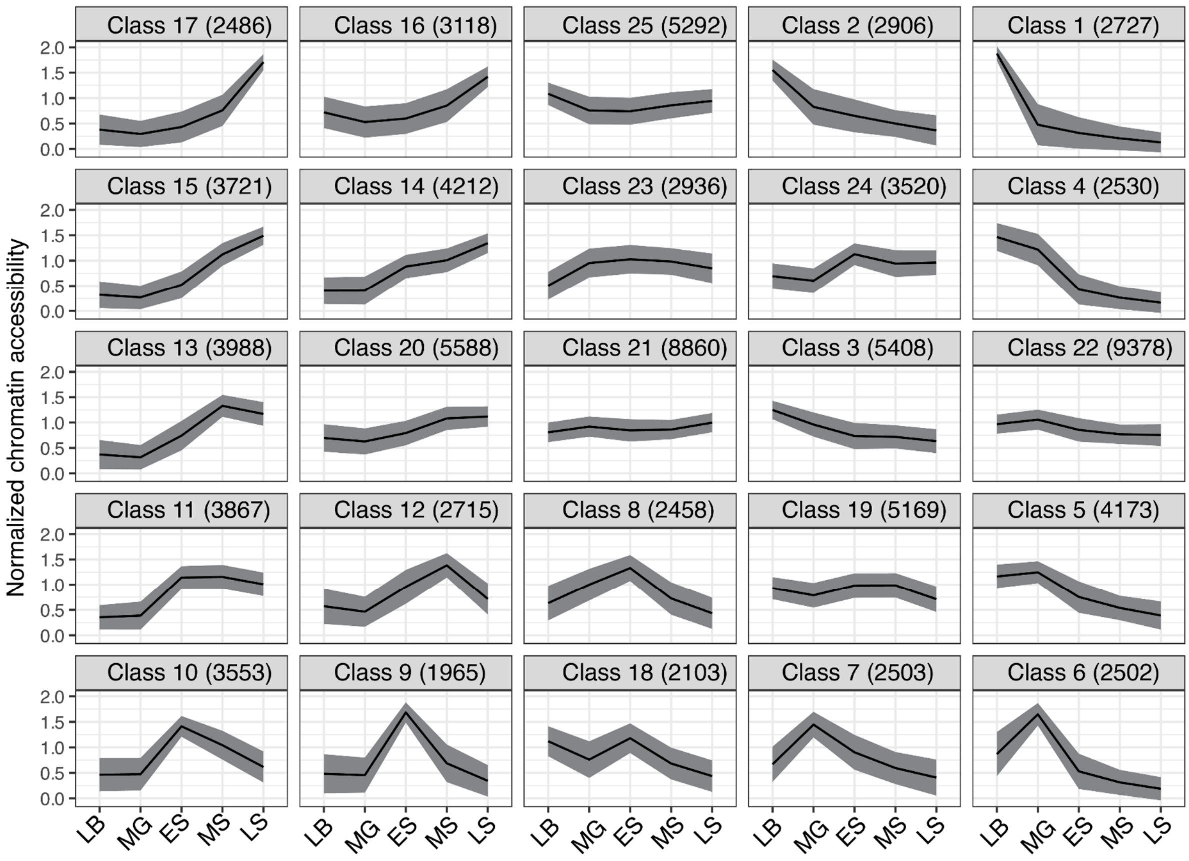

**Fig. S20. Atlantic salmon DevMap chromatin openness SOM clusters**. Each individual line plot represents one SOM cluster and shows average normalised chromatin openness/accessibility, with the position on the panel assigned by the SOM algorithm based on the inferred topology of the data. Shaded areas around each line plot show 95% confidence intervals. The top of each line plot indicates the SOM class number, along with the number of robust open chromatin regions captured per class. Stages are indicated using abbreviations as x-axis labels: LB, Late blastula; MG, Mid gastrula; ES, Early somitogenesis; MS, Mid somitogenesis; LS, Late somitogenesis.

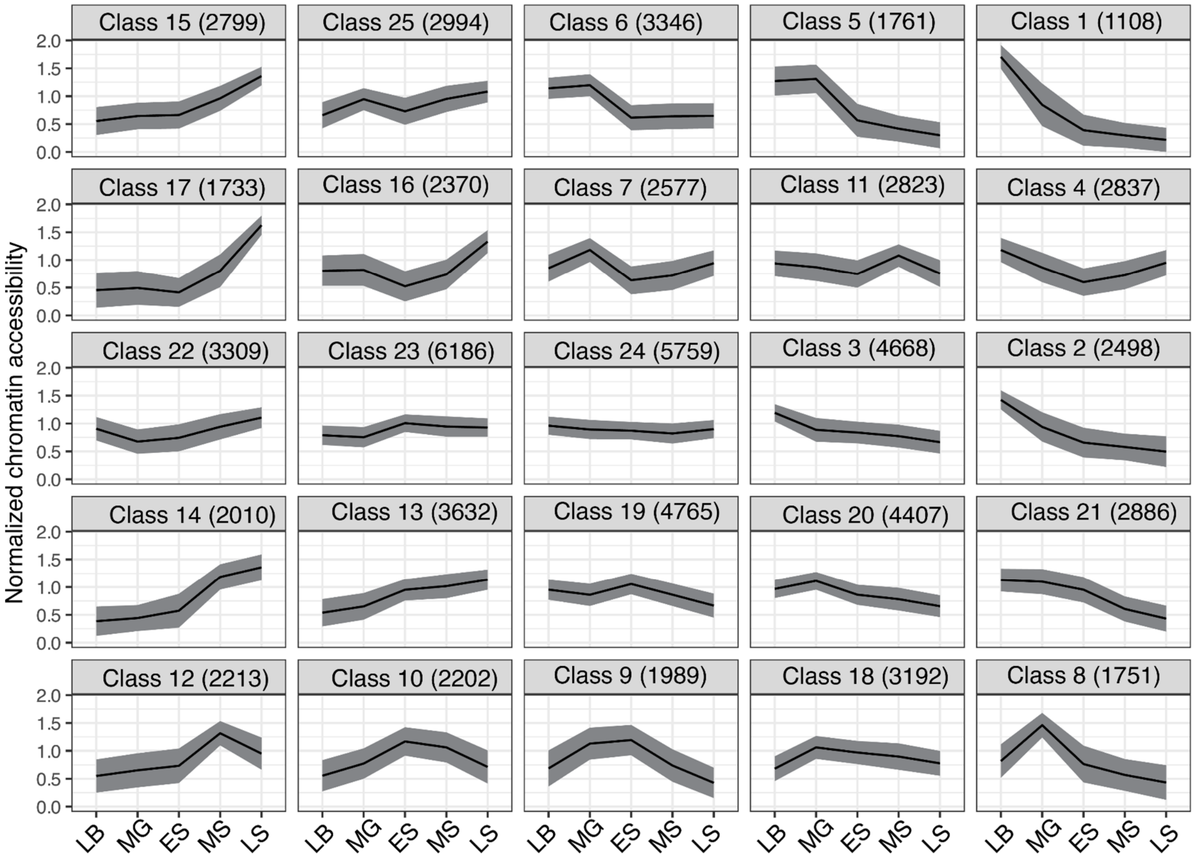

**Fig. S21. Rainbow trout DevMap chromatin openness SOM clustering.** Other details are as per the Fig. S20 legend.

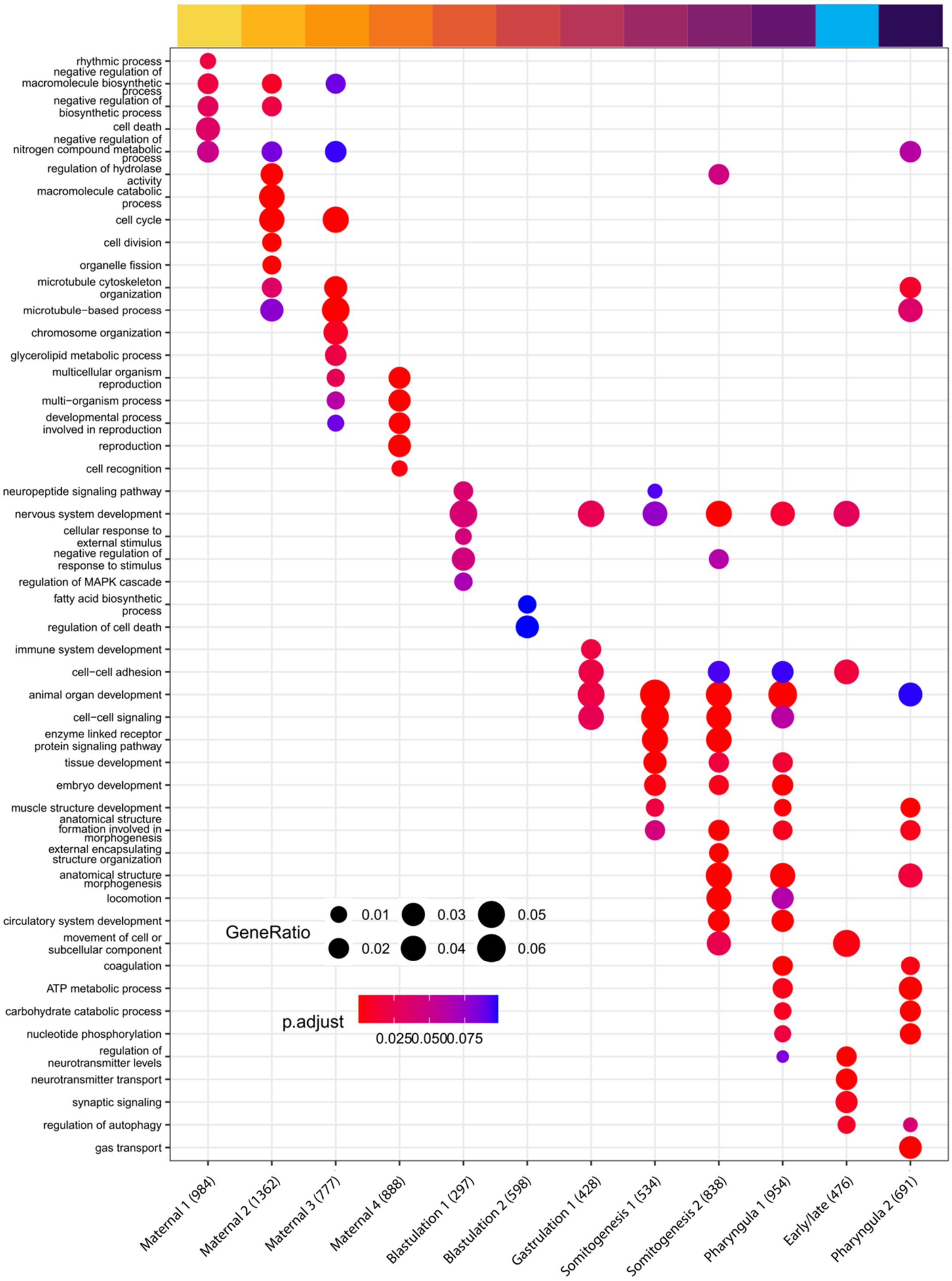

**Fig. S22. Enriched gene ontology (GO) terms for DevMap transcriptome of Atlantic salmon.** The dot plot compares the most enriched GO terms across SOM clusters shown on the x-axis (shown in Fig. S18), restricted to 'Biological Process' terms. Each cluster shows a maximum of five distinct terms, based on FDR adjusted p-value, and additional terms also enriched in other clusters. The y-axis provides GO terms, with the size of each dot indicating gene ratio and colours indicating FDR adjusted p-value according to the shown scales. Numbers in labels indicate the number of genes in the SOM cluster matched to at least one GO term. The cluster named "early/late" was not included in Fig. 2 (main text).

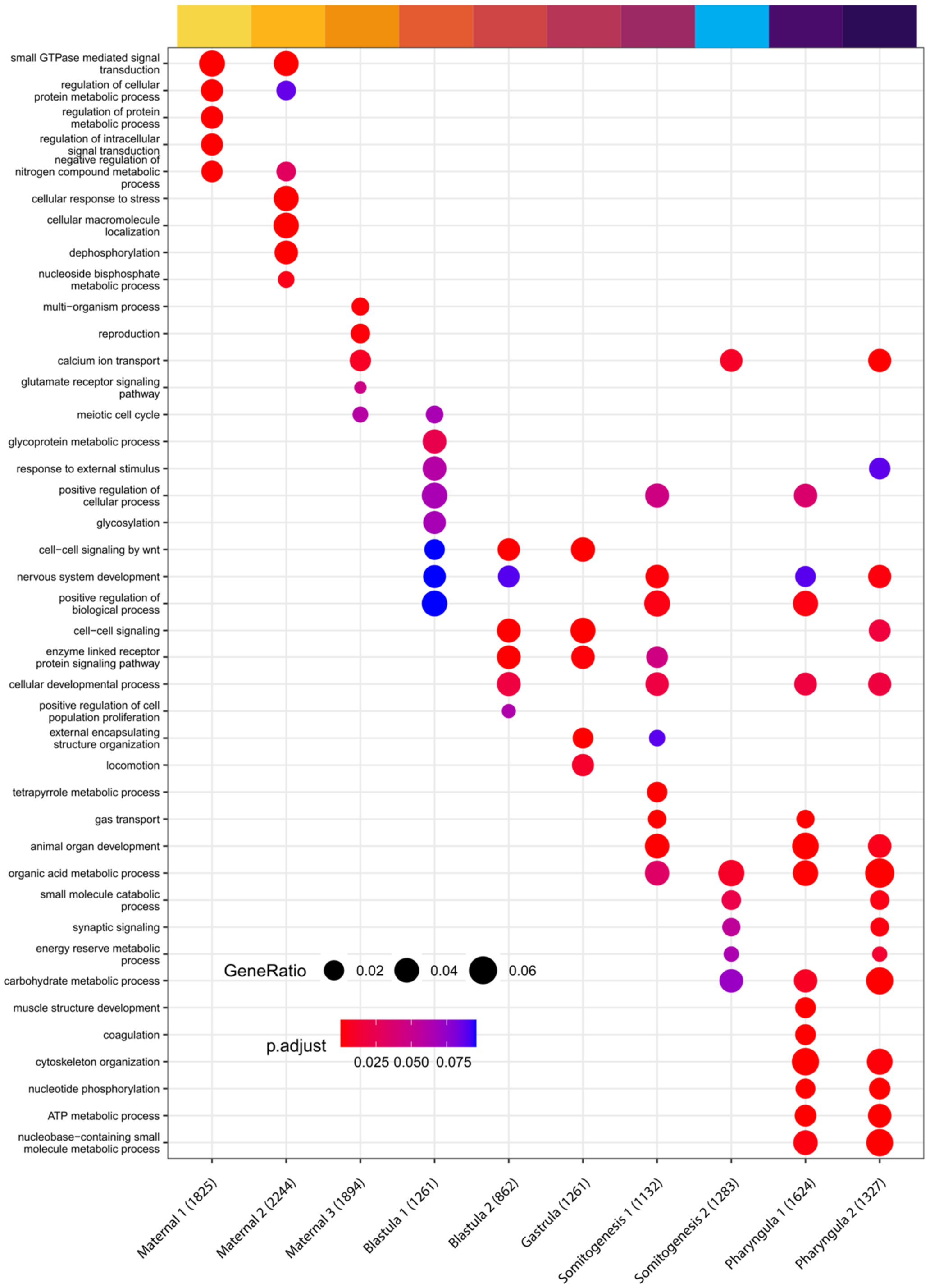

**Fig. S23. Enriched gene ontology (GO) terms for DevMap transcriptome of rainbow trout.** The dot plot compares the most enriched GO terms across SOM clusters shown on the x-axis (shown in Fig. S19), restricted to 'Biological Process' terms. Each cluster shows a maximum of five distinct terms, based on FDR adjusted p-value, and additional terms also enriched in other clusters. The y-axis provides GO terms, with the size of each dot indicating gene ratio and colours indicating FDR adjusted p-value according to the shown scales. Numbers in labels indicate the number of genes in the SOM cluster matched to at least one GO term.

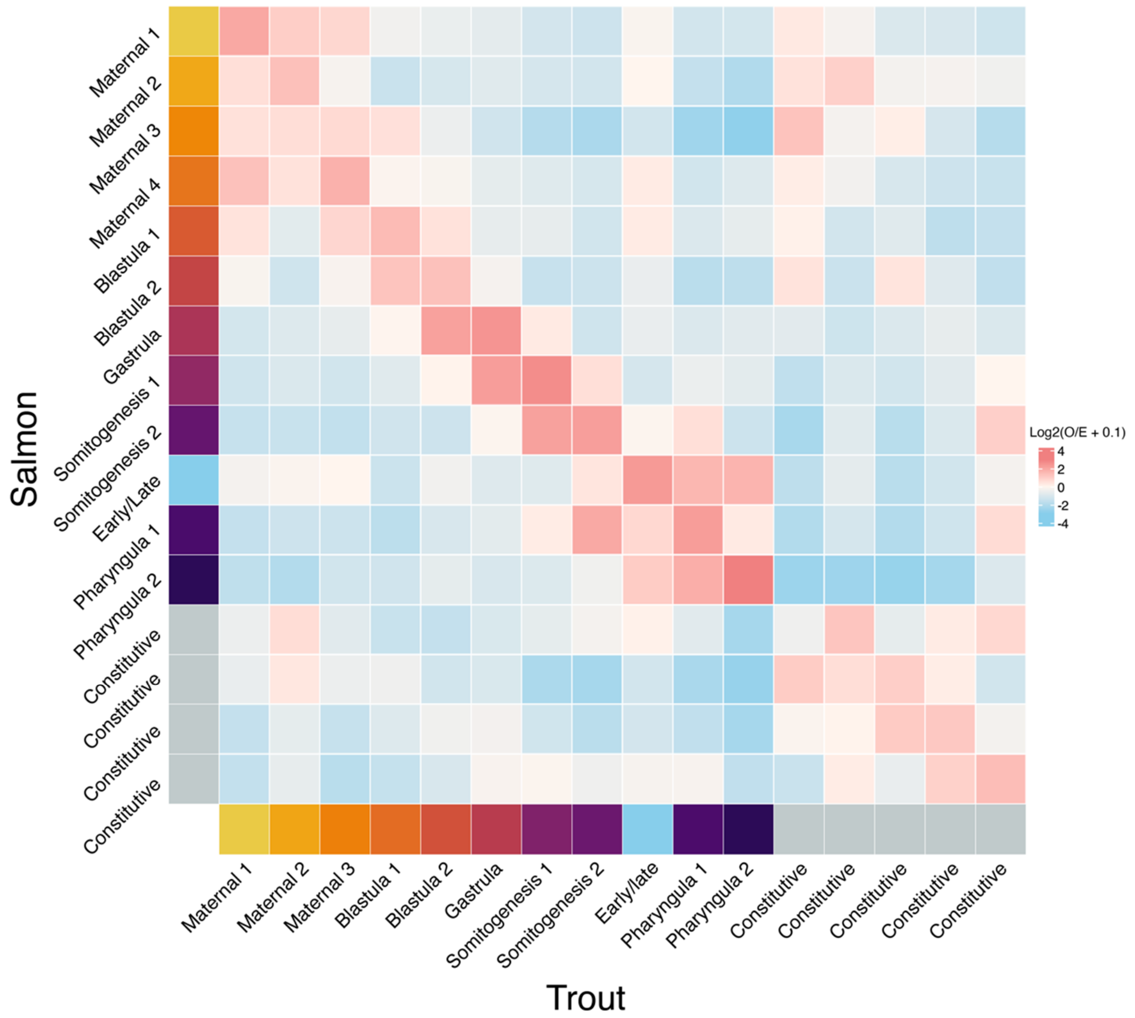

**Fig. S24. Matrix of observed to expected ratio of orthologs shared across all combinations of Atlantic salmon and rainbow trout DevMap SOM clusters**. Expected values were calculated as the mean of 10,000 random distributions of trout genes across SOM clusters. Colours represent log-2 of observed to expected ratio, such that positive and negative values represent ortholog enrichment and depletion, respectively. Axis labels indicate SOM cluster names, as shown in Fig. S18 (Atlantic salmon) and Fig. S19 (rainbow trout).

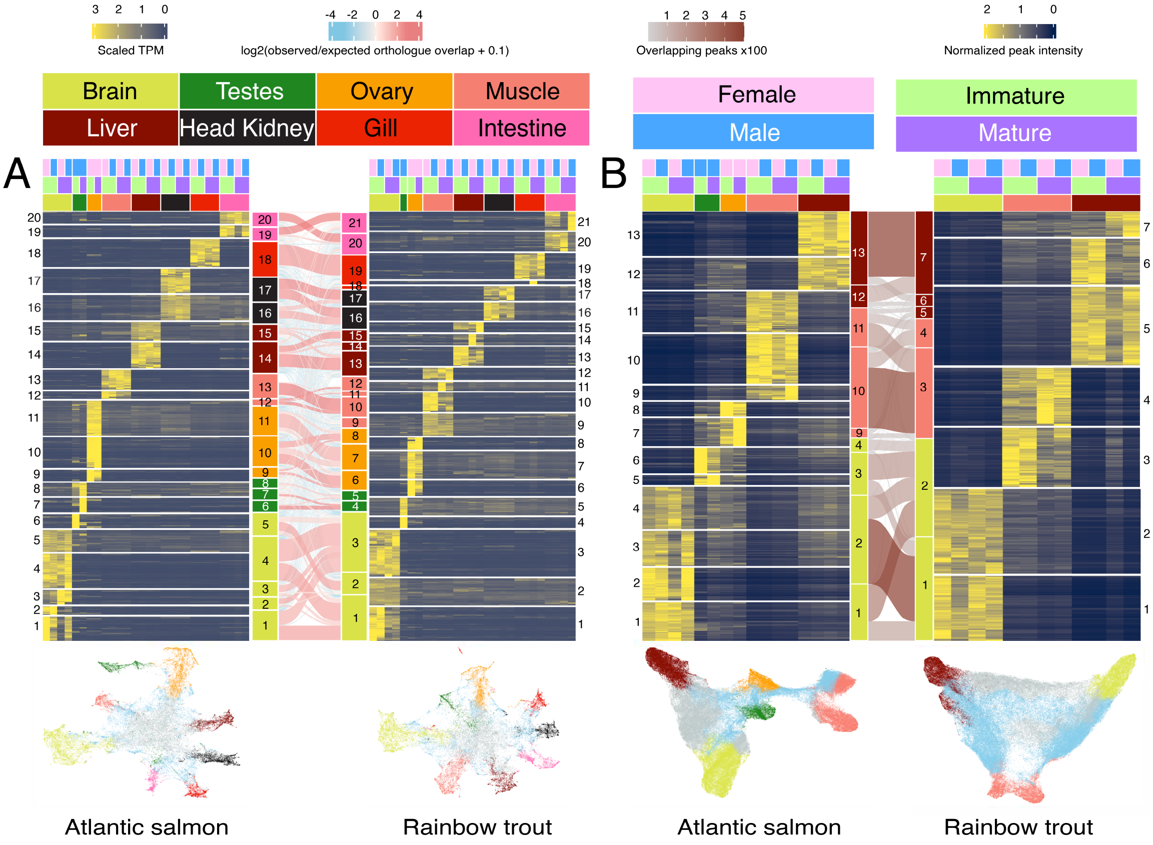

**Fig. S25**. **Functional activity of salmonid genomes in adult tissues at two stages of sexual maturation**. (**A**) Transcriptome and (**B**) chromatin openness dynamics in adult tissues from both sexes at two stages of sexual maturation (BodyMap). Respective data shown are SOM clusters for mean normalised transcripts per million values (TPM, RNA-Seq) and mean normalised read intensity values in robust open chromatin regions (ATAC-seq). Full BodyMap SOM clusters are shown in Figs S26-29. For panel **A**, light red bands illustrate the observed-to-expected ratio of orthologous gene expression between the two species across all cluster combinations. For panel B, brown bands between clusters show the number of syntenic open chromatin regions based on whole genome alignment. On the UMAP plots shown under each heatmap, each dot represents mean normalised TPM values of individual genes (panel **A**) or mean normalised ATAC-seq read intensity values in robust open chromatin regions (panel **B**). Grey dots and light blue dots on each UMAP are genes or chromatin regions captured within constitutive and multi-tissue expression SOM clusters (Figs S26-S29), respectively. This analysis captures tissue-specific clusters of expression and chromatin openness, alongside clusters capturing genes expressed in multiple tissues (Figs S26-29). We also note clusters showing various levels of specificity by sex or maturation stage (Figs S26-29). We found that tissue-specific clusters were enriched for GO terms consistent with organ functions, including immunity (head kidney, gill, liver), hematopoiesis (head kidney), metabolism (liver, muscle), oogenesis (ovary), meiosis (ovary, testis), spermatogenesis (testis), the nervous system (brain), the circulatory system (gill, brain) and ion transport (gill, distal intestine) (Figs S30-S31; Tables S7-S8; Text S7). The UMAPs evidences heterogeneity consistent with the differentiated tissues profiled. The central UMAP region is populated with genes or chromatin regions captured by constitutive or multi-tissue clusters (Figs S26-29), while features captured by tissue-specific clusters presented as protruding arms. Enrichment of orthologous gene expression and open chromatin regions was observed across tissue-specific clusters for both species (Fig. S32).

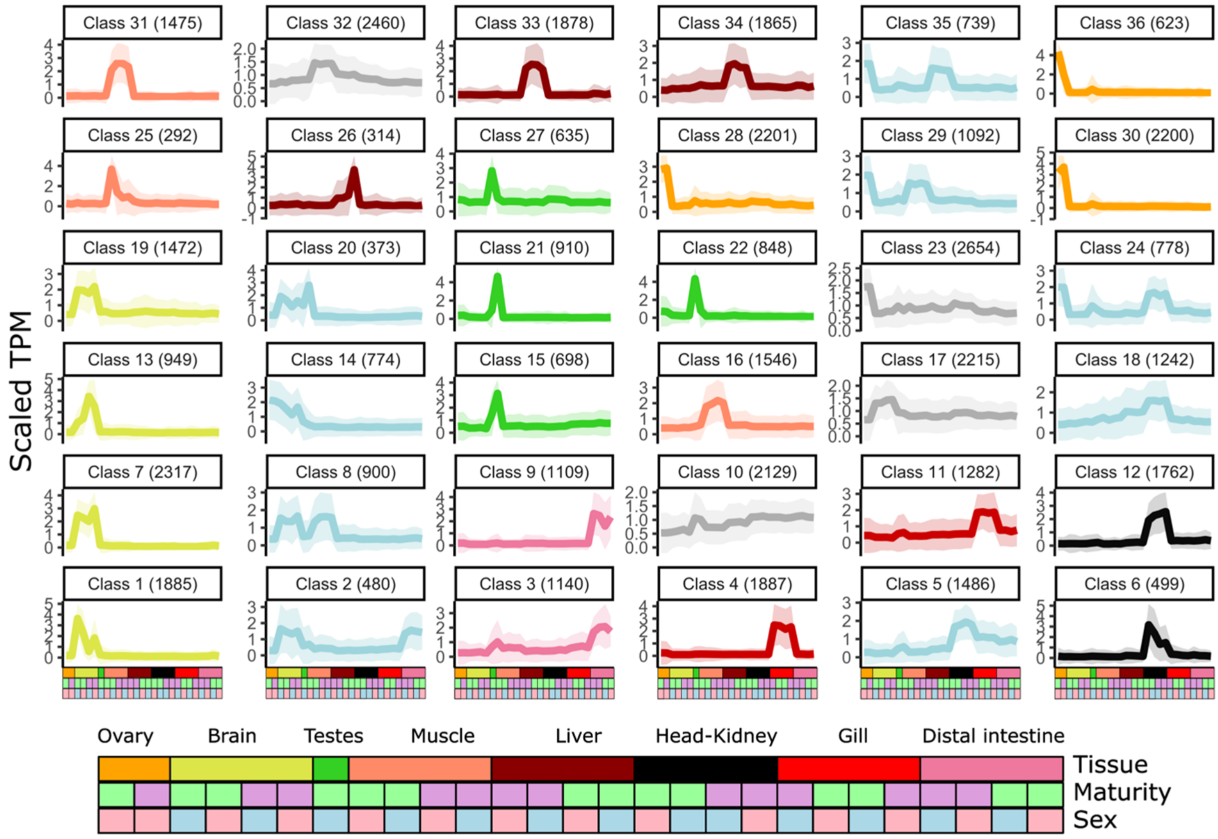

**Fig. S26. Atlantic salmon BodyMap transcriptome clusters**. Each individual line plot represents one SOM cluster and shows average normalised gene expression, with the position on the panel assigned by the SOM algorithm based on the inferred topology of the data. The colours indicate the tissue-specific expression of each cluster (based on colours used in Fig. 2), with additional clusters presented here (In grey, constitutive clusters, and in light blue, clusters expressed across several tissues). Coloured areas around each line plot show 95% confidence intervals. The top of each line plot indicates the number of genes captured per cluster. Each sample is indicated using the colours shown in the bottom annotation.

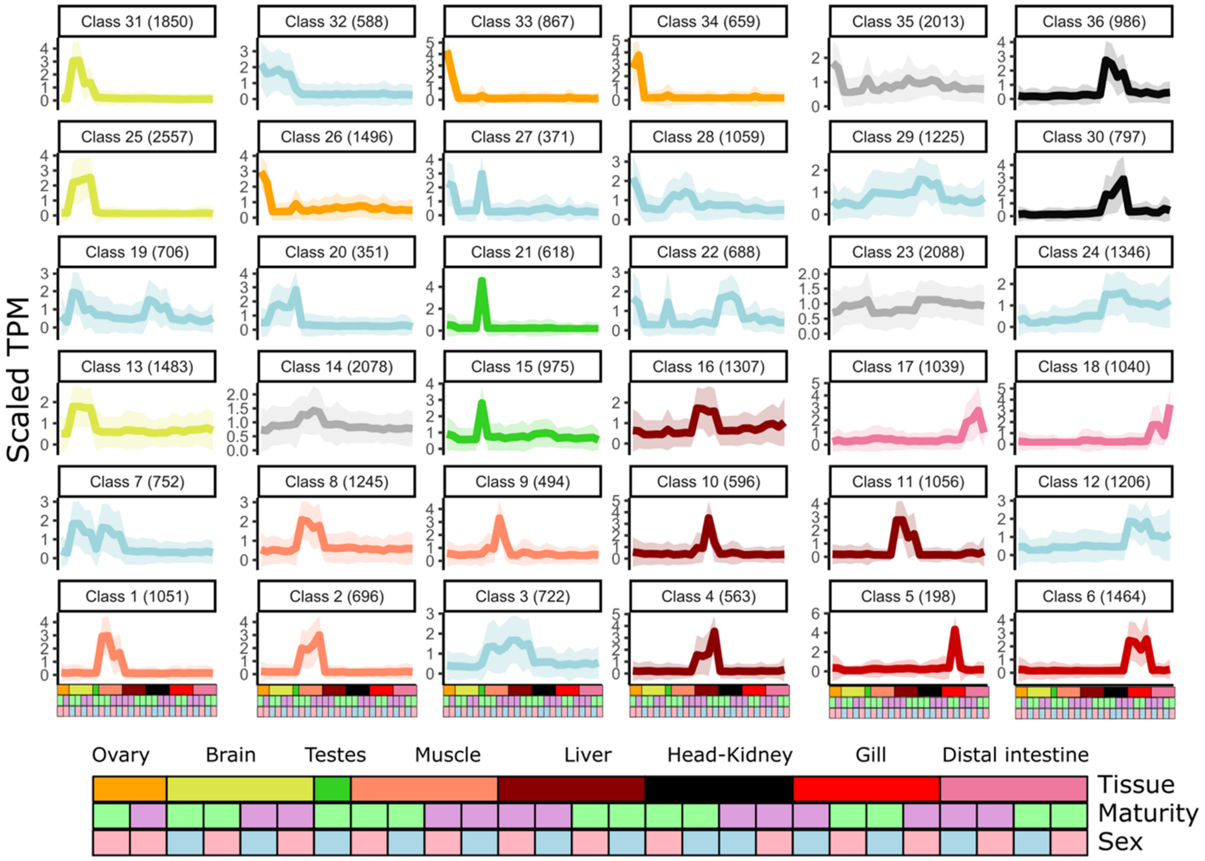

**Fig. S27. Rainbow trout BodyMap transcriptome clusters**. Other details are as per the Fig. S26 legend.

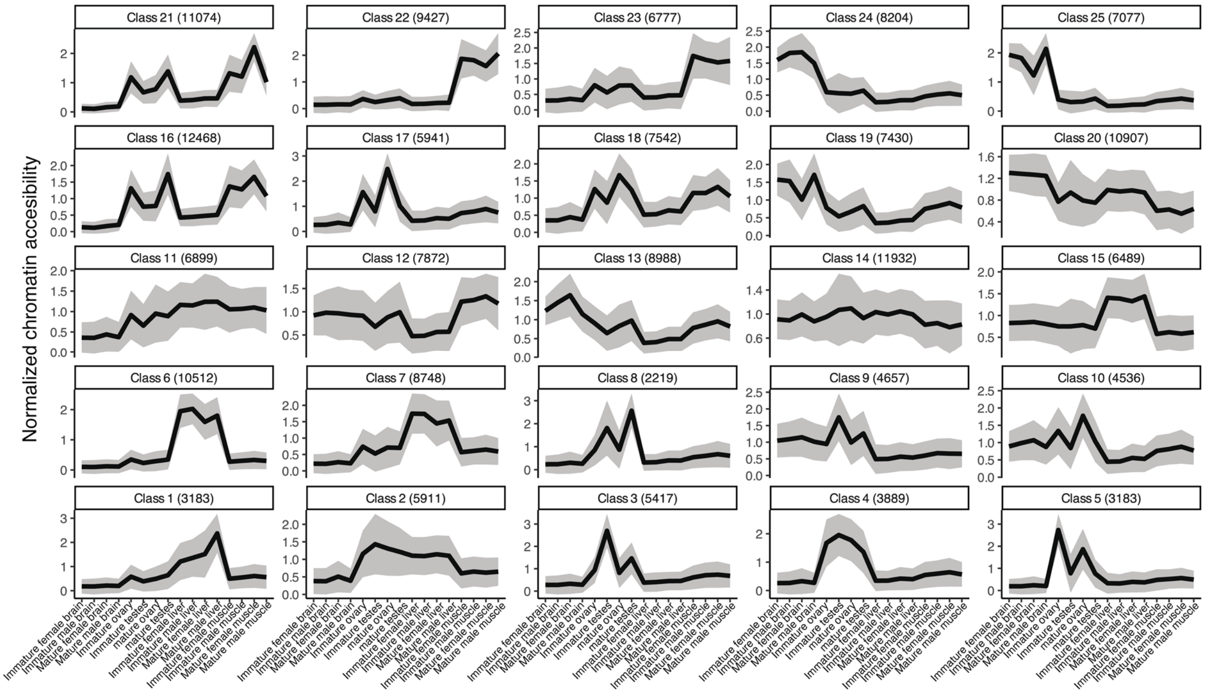

**Fig. S28. Atlantic salmon BodyMap chromatin openness SOM clusters**. Each individual line plot represents one SOM cluster and shows average normalised chromatin openness, with the position on the panel assigned by the SOM algorithm based on the inferred topology of the data. Shaded areas around each line plot show 95% confidence intervals. The top of each line plot indicates the SOM class number, along with the number of robust open chromatin regions captured per class. The x-axis labels represent sample information, including tissue type, sex and stage of maturity

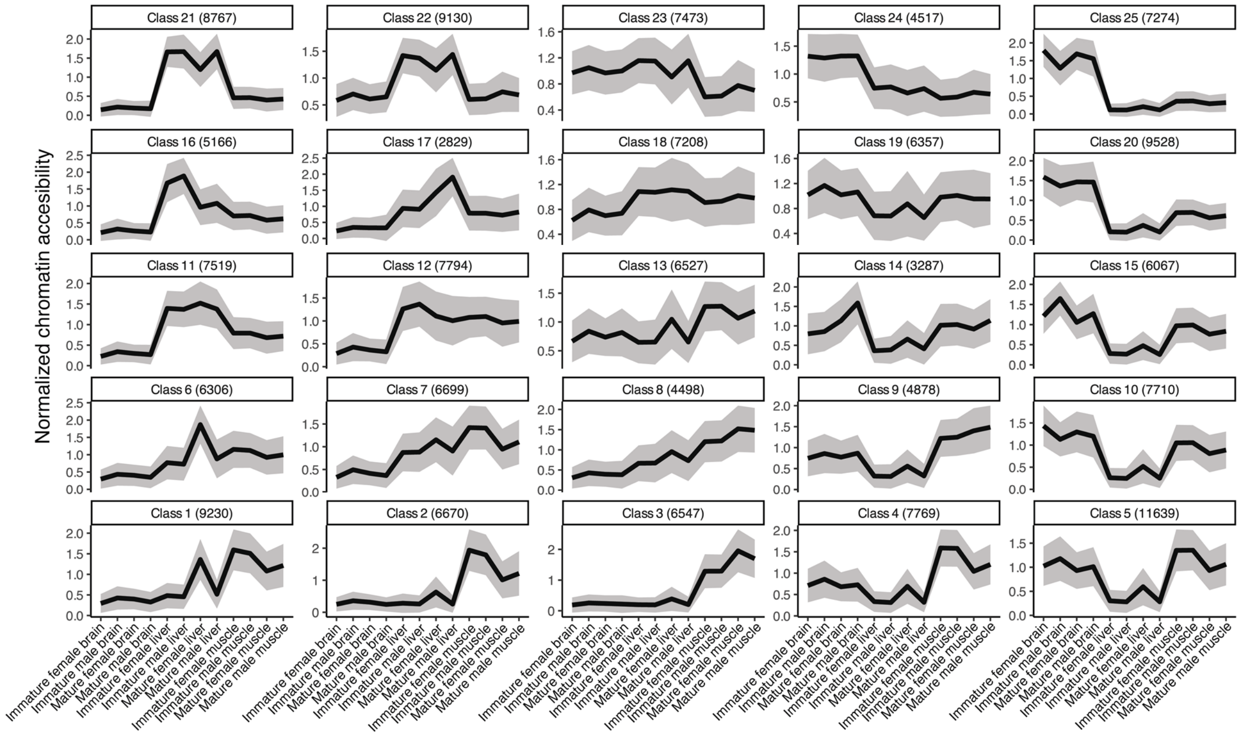

**Fig. S29. Rainbow trout BodyMap chromatin openness SOM clusters.** Other details are as per the Fig. S28 legend.

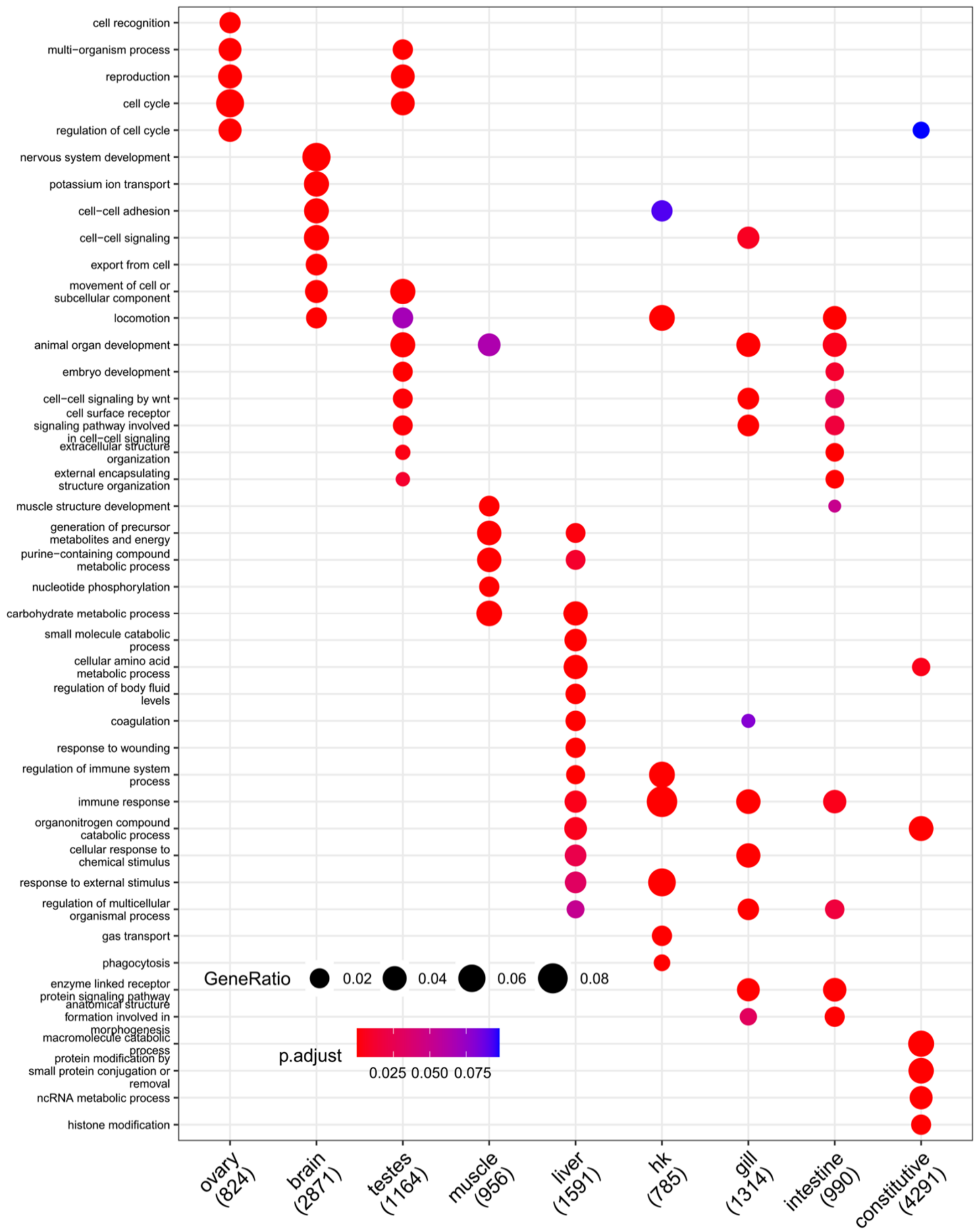

**Fig. S30**. **Enriched gene ontology (GO) terms for BodyMap transcriptome of Atlantic salmon.** The dot plot compares the most enriched GO terms across SOM clusters shown on the x-axis (shown in Fig. S25), restricted to 'Biological Process' terms. Each cluster shows a maximum of five distinct terms, based on FDR adjusted p-value, and additional terms also enriched in other clusters. The y-axis provides GO terms, with the size of each dot indicating gene ratio and colours indicating FDR adjusted p-value according to the shown scales. Numbers in labels indicate the number of genes in the SOM cluster matched to at least one GO term. 'hk' = head kidney, 'intestine' = distal intestine.

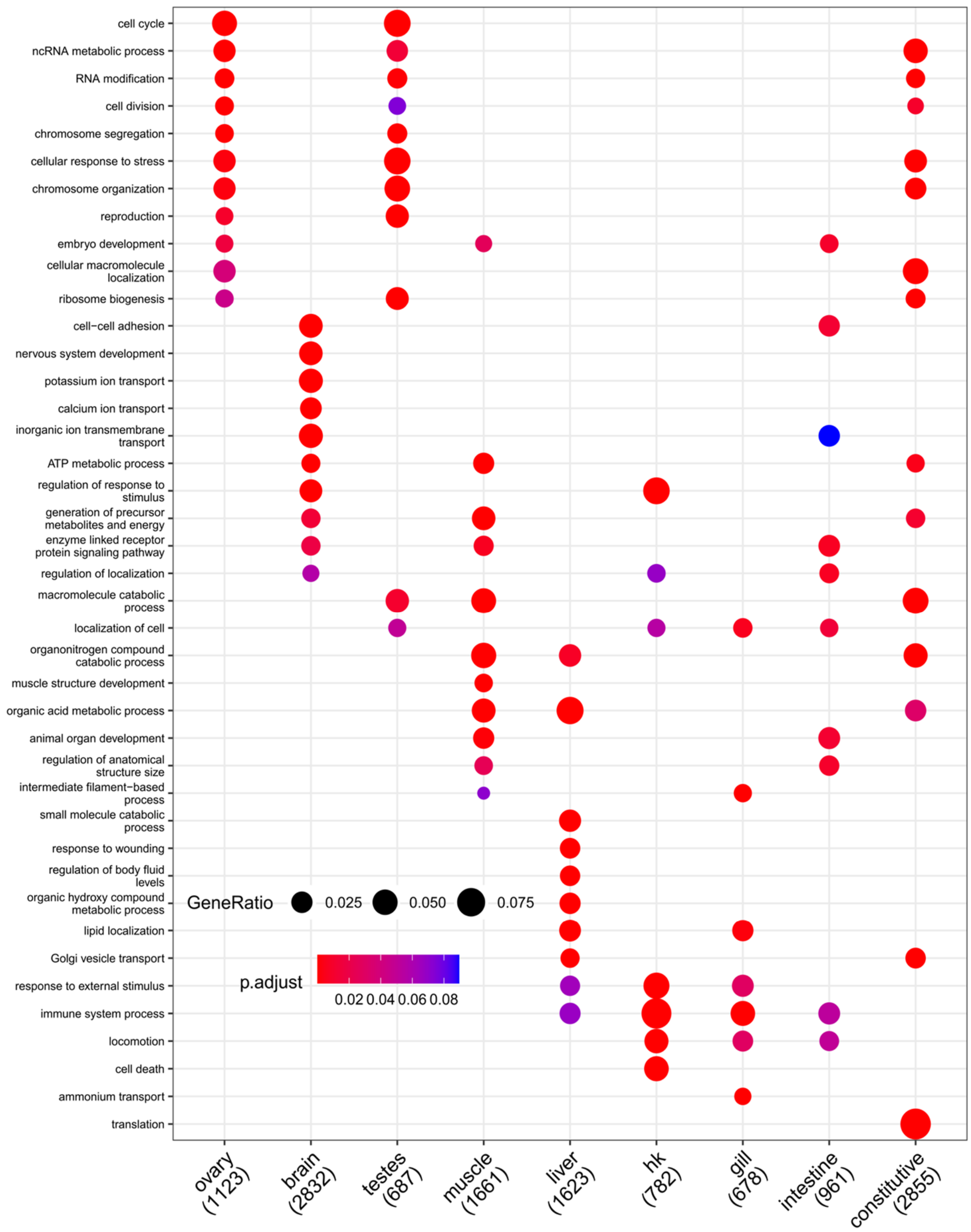

**Fig. S31**. **Enriched gene ontology (GO) terms for BodyMap transcriptome of rainbow trout.** The dot plot compares the most enriched GO terms across SOM clusters shown on the x-axis (shown in Fig. S26), restricted to 'Biological Process' terms. Each cluster shows a maximum of five distinct terms, based on FDR adjusted p-value, and additional terms also enriched in other clusters. The y-axis provides GO terms, with the size of each dot indicating gene ratio and colours indicating FDR adjusted p-value according to the shown scales. Numbers in labels indicate the number of genes in the SOM cluster matched to at least one GO term. 'hk' = head kidney, 'intestine' = distal intestine.

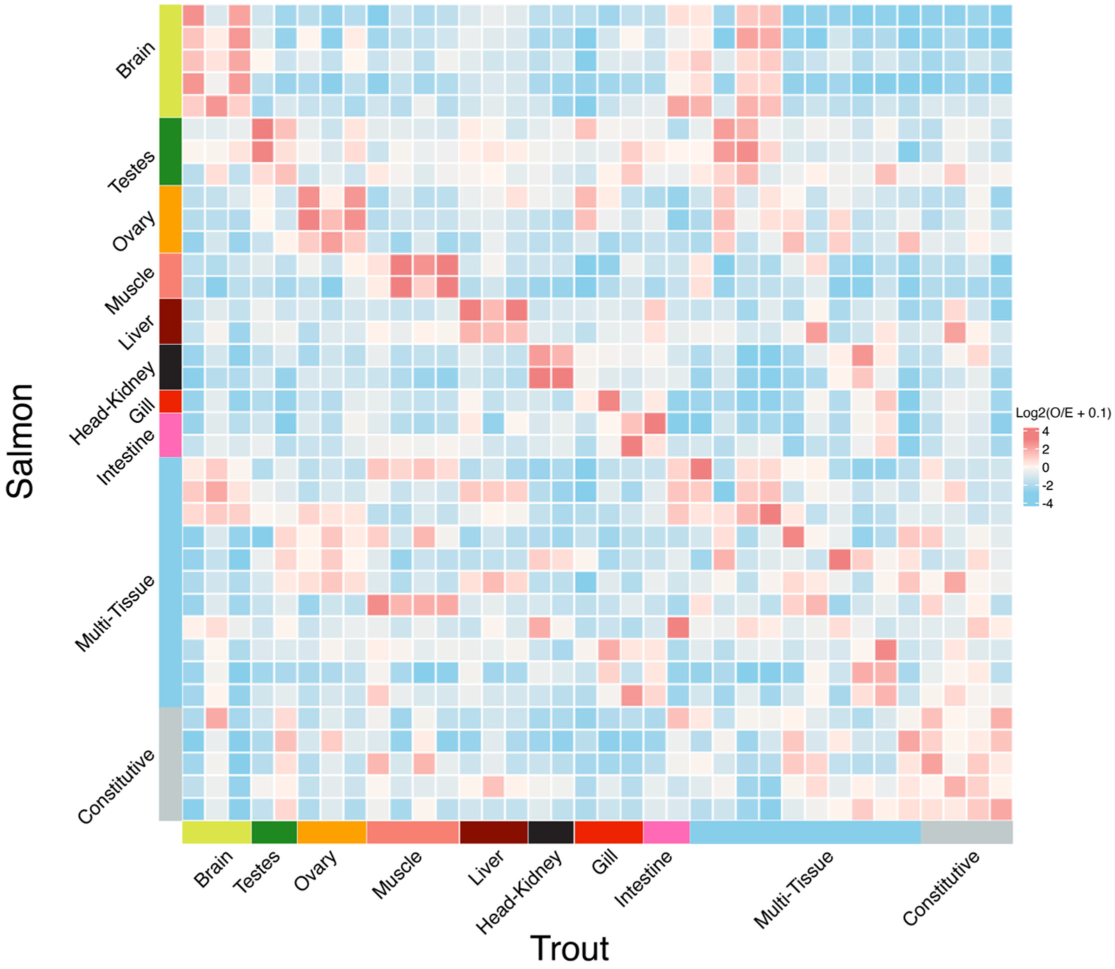

**Fig. S32. Matrix of observed to expected ratio of orthologs shared across all combinations of Atlantic salmon and rainbow trout BodyMap SOM clusters.** Expected values were calculated as the mean of 10,000 random distributions of trout genes across SOM clusters. Colours represent log-2 of observed to expected ratio, such that positive and negative values represent ortholog enrichment and depletion, respectively. Axis labels indicate SOM cluster names, as shown in Fig. S26 (Atlantic salmon) and Fig. S27 (rainbow trout). ‘Intestine’ = distal intestine.

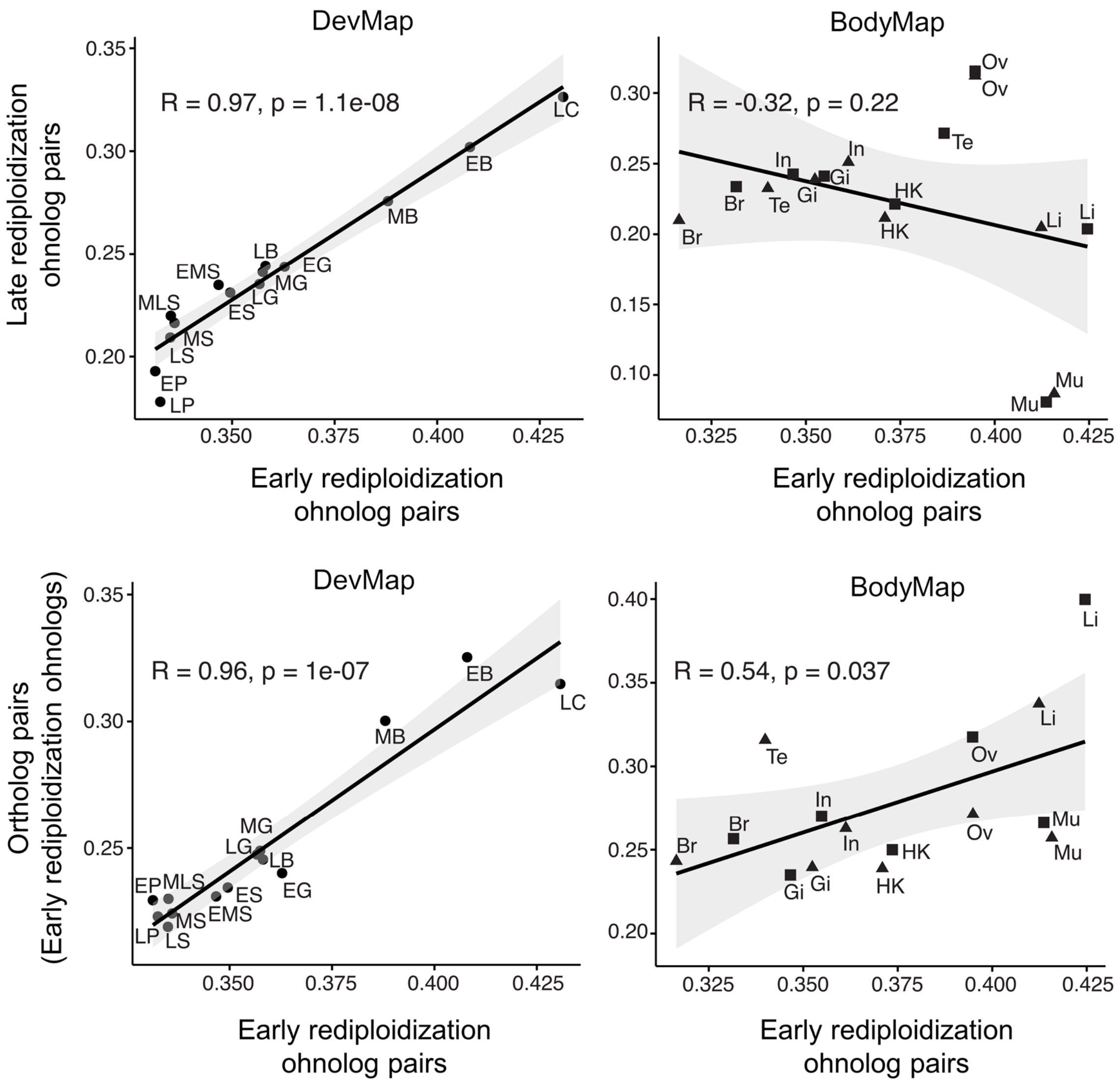

**Fig. S33. Scatterplots comparing mean JSD values between pairs of ohnologs and orthologs across ontogeny and tissues**. The solid lines represent the least-squares fit and the shaded grey bands are 95 % confidence intervals. Pearson's R and p values are shown for each comparison.

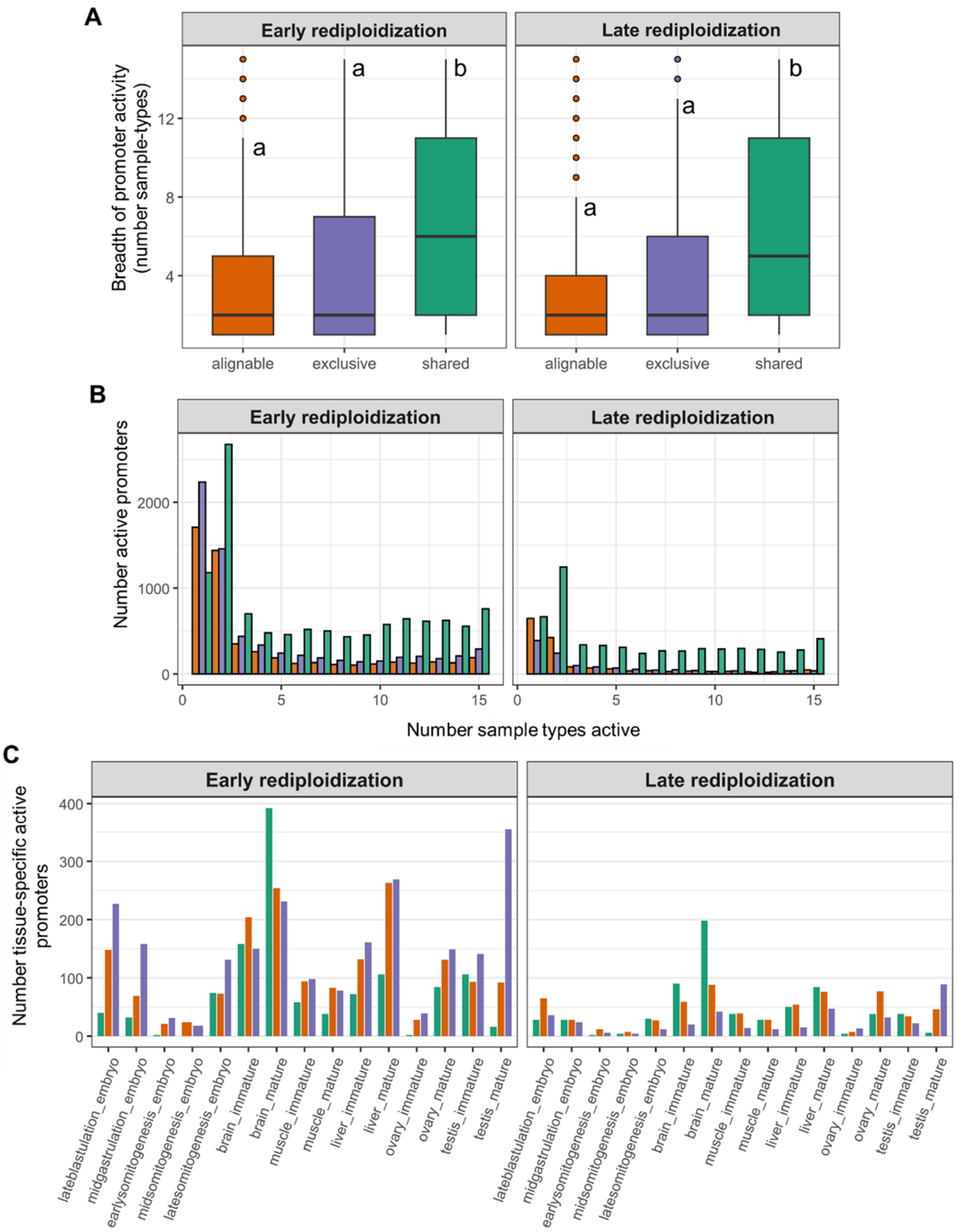

**Fig. S34. Dynamics of Atlantic salmon promoter activity across sample types, rediploidization regions and WGD alignment categories**. (**A**) Breadth of promoter activity summarized across samples types and stages in the DevMap and BodyMap atlases (embryo stages, adult tissues at immature and mature stages) compared between the Exclusive, Alignable only and Shared categories. Shared promoters have the broadest activity across sample types (unique lowercase letters indicate significant differences according to Wilcoxon's rank test; [Left] Early regions: Alignable only vs. Exclusive: *p*-adj=0.006; Alignable only vs. Shared: *p*-adj=9.96e-287; and Exclusive vs. Shared: *p*-adj=4.64e-261; [right] Late regions: Alignable only vs. Exclusive: *p*-adj=1.63e-12; Alignable only vs. Shared: *p*-adj=1.26e-157; and Exclusive vs. Shared: *p*-adj=2.61e-61. (**B**) Consistent with part **A**, many more active promoters from the Shared category (retained as active ohnolog pairs) are active in a higher number of distinct sample types. (**C**) The number of tissue-specific active promoters varies extensively across sample types and alignment categories. For both Early and Late rediploidization regions, mature brain has the highest number of active ohnolog pairs that are inactive in other sample types, whereas mature testis has the highest number of sample-type unique Exclusive promoters.

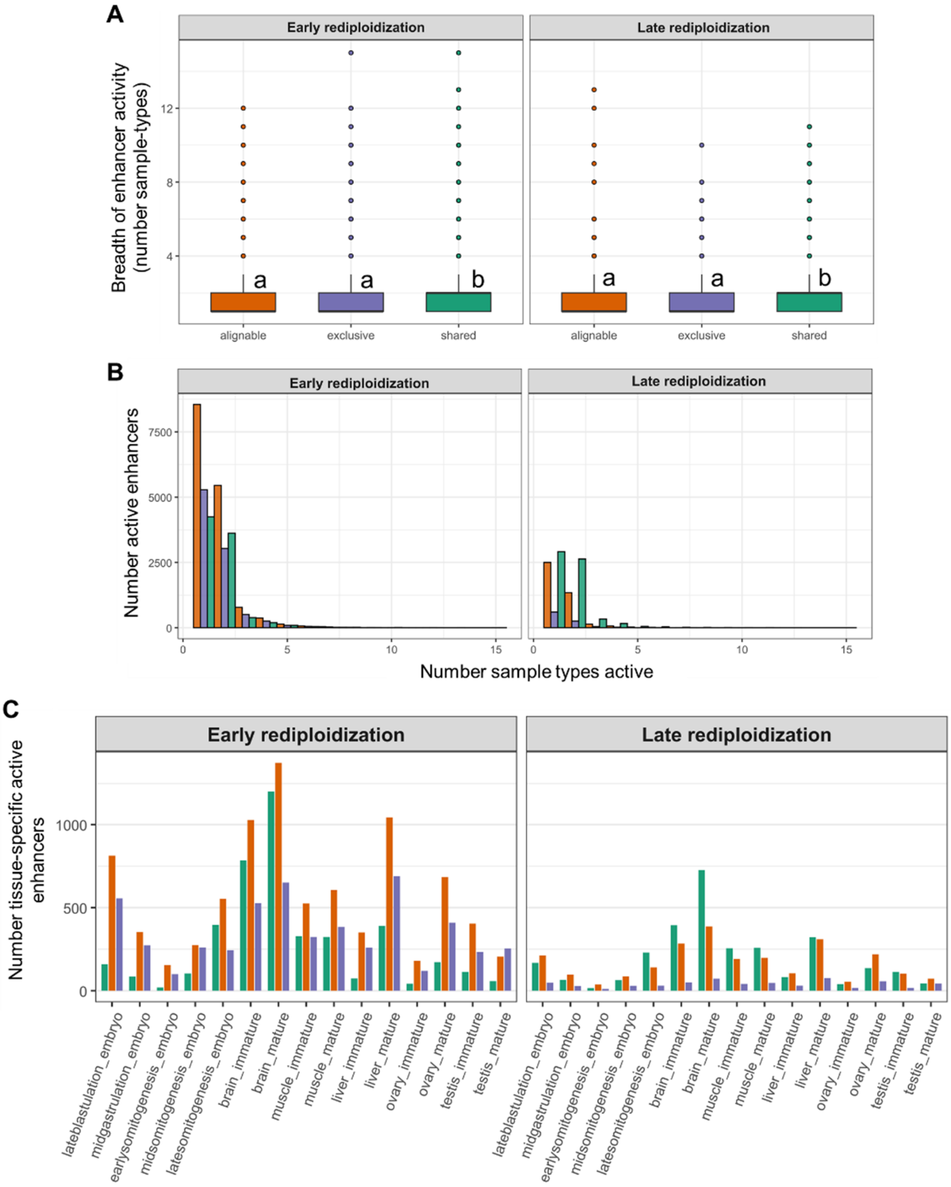

**Fig. S35. Dynamics of Atlantic salmon enhancer activity across sample types, rediploidization regions and WGD alignment categories.** (**A**) Breadth of enhancer activity summarized across samples types and stages in the DevMap and BodyMap atlases (embryo stages, adult tissues at immature and mature stages) compared between the Exclusive, Alignable only and Shared categories. There is no difference in the breadth of enhancer activity in different sample types across the Exclusive and Alignable only categories. However, the Shared enhancers retained as active ohnolog pairs show significantly higher activity breadth than both other alignment categories (unique lowercase letters indicate significant differences according to Wilcoxon's rank test; [Left] Early regions: Alignable only vs. Exclusive: *p*-adj=0.213; Alignable only vs. Shared: *p*-adj=1.16e-15; and Exclusive vs. Shared: *p*-adj=4.92e-16; [right] Late regions: Alignable only vs. Exclusive: *p*-adj=0.260; Alignable only vs. Shared: *p*-adj=2.66e-43; and Exclusive vs. Shared: *p*-adj=3.40e-18. (**B**) Consistent with **A**, the majority of enhancers from all alignment categories show activity in a small breadth of different tissue types. (**C**) The number of tissue-specific active enhancers varies across sample types and alignment categories.

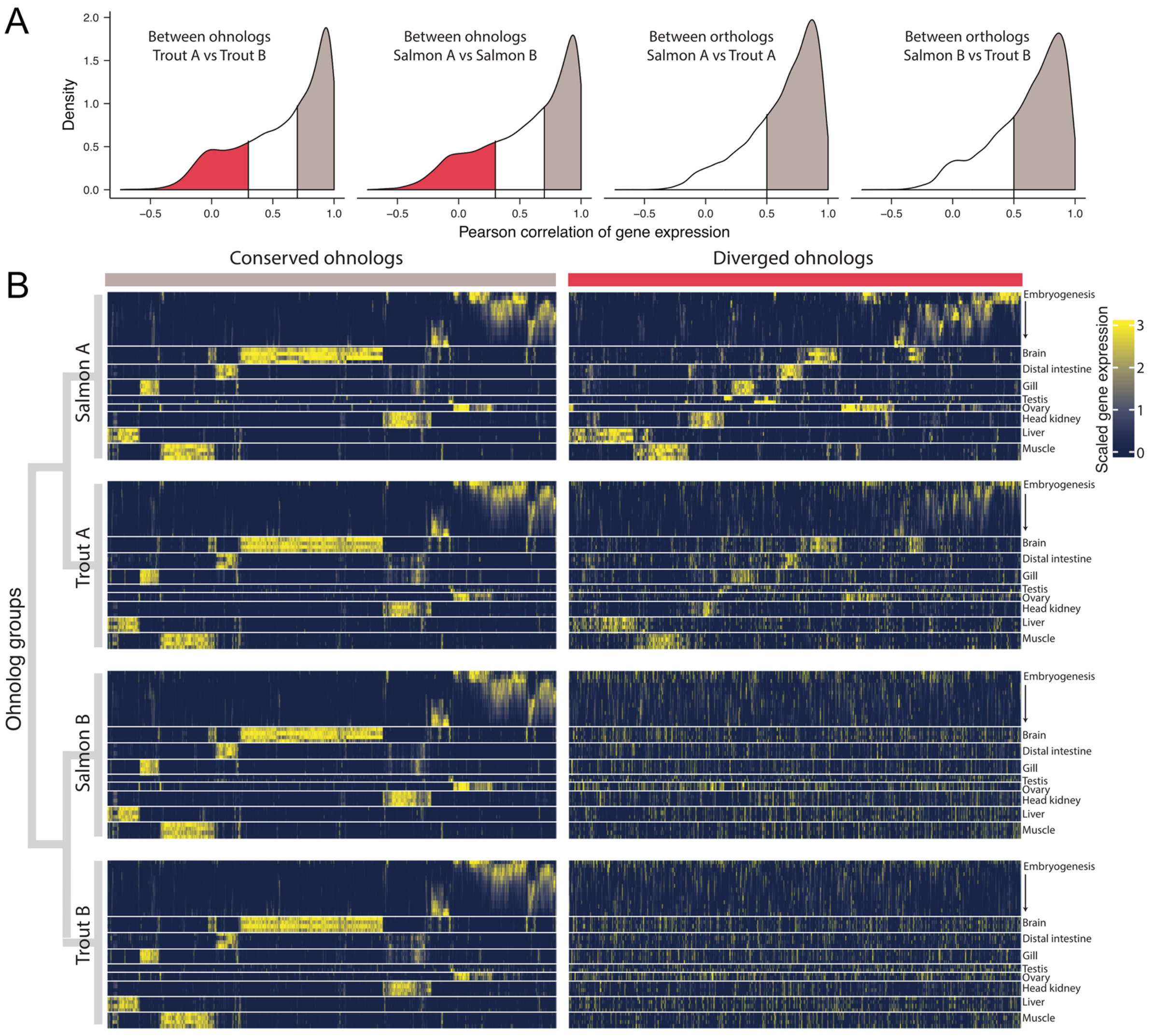

**Fig. S36. Defining 'conserved' and 'diverged' expression conservation classes for 2:2 ohnolog pairs**. (**A**) Distribution of Pearson's correlation values calculated between expression profiles between 2:2 ohnolog pairs within species, and orthologs of the same from Atlantic salmon and rainbow trout. Threshold for "conserved" ohnologs = R > 0.7 between 2:2 ohnolog pairs (both species) and R > 0.5 between ortholog. Threshold for "diverged" ohnologs = R ≤ 0.3 between ohnolog pairs (both species). (**B**) Heatmap of scaled gene expression across sample types for "conserved" and "diverged" ohnologs in both species.

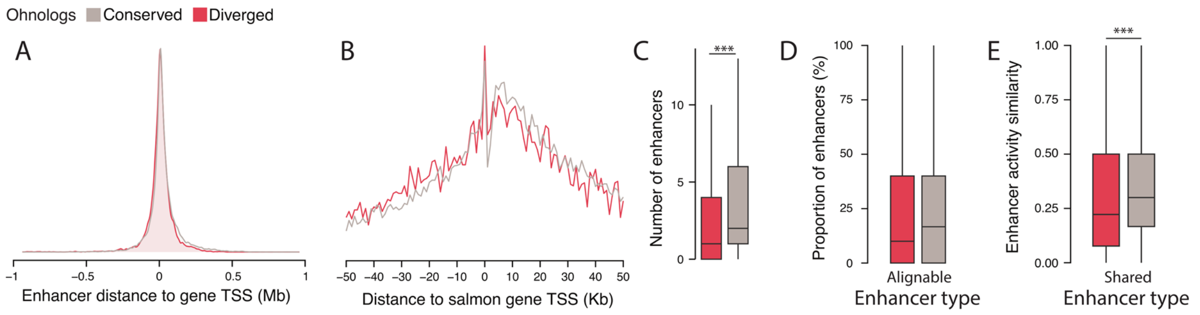

**Fig. S37. Comparing Atlantic salmon ohnolog enhancers within the "conserved" (grey) and "diverged" (red) expression categories**. (**A**) Density distribution of enhancer distances to gene TSS within regulatory domains. (**B**) Frequency of enhancer distances within +/-50 Kb of gene TSS. (**C**) We observed a difference in the number of enhancers assigned to regulatory domains of "conserved" and "diverged" ohnologs (Wilcoxon rank sum test: $p = 2.2 \times 10^{-16}$). (**D**) However, there was no difference in the proportion of enhancers (24.9% vs 23.4%) belonging to the Alignable only category (binomial GLM, $p = 0.058$, odd ratio = 0.92, 95% CI 0.85–1.00). (**E**) We observed a difference in Shared enhancer activity similarity between the expression classes, as measured by the presence or absence of activity across all sample types (respective median similarity for conserved and diverged expression classes = 0.30 vs 0.22; Wilcoxon rank sum test: $p = 1.2 \times 10^{-6}$).

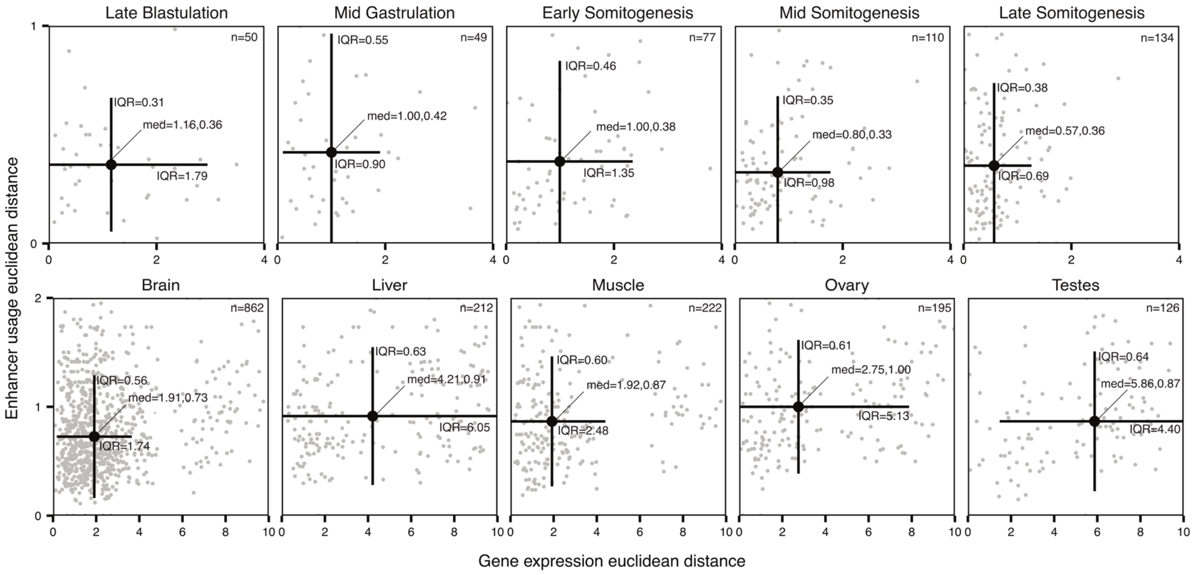

**Fig. S38. Ohnolog expression distances against enhancer distances in Atlantic salmon**. Euclidean distances between 2:2 ohnolog pairs are compared, for scaled gene expression profiles across sample types against enhancer-usage (proportions of total enhancers active in each sample type). Ohnologs were separated by stage of embryogenesis (top row, DevMap) or tissue type (bottom row, BodyMap) using SOM clustering (main text, Fig. 2). Number of ohnolog pairs per cluster is shown in the top right (n=x). Median values are shown ('med') for expression and enhancer distances respectively, with lines showing upper and lower interquartile ranges (IQR).

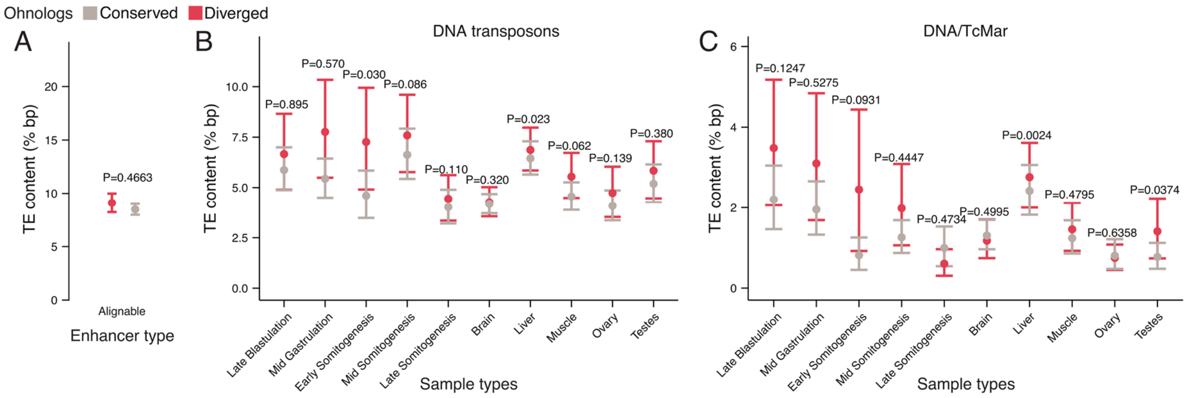

**Fig. S39. TE content in ohnolog enhancers in Atlantic salmon**. (**A**) Mean percentage of all TE bp overlapping Alignable only enhancers of conserved (grey) and diverged (red) ohnologs. Also shown is the content of: (**B**) DNA transposons and (**C**) TcMar class DNA transposons, in Exclusive enhancers based on activity in specific sample types. Probability values are from Wilcoxon rank sum tests between "conserved" and "diverged" ohnologs. 95% confidence intervals were calculated from bootstrapping with 10,000 replicates.

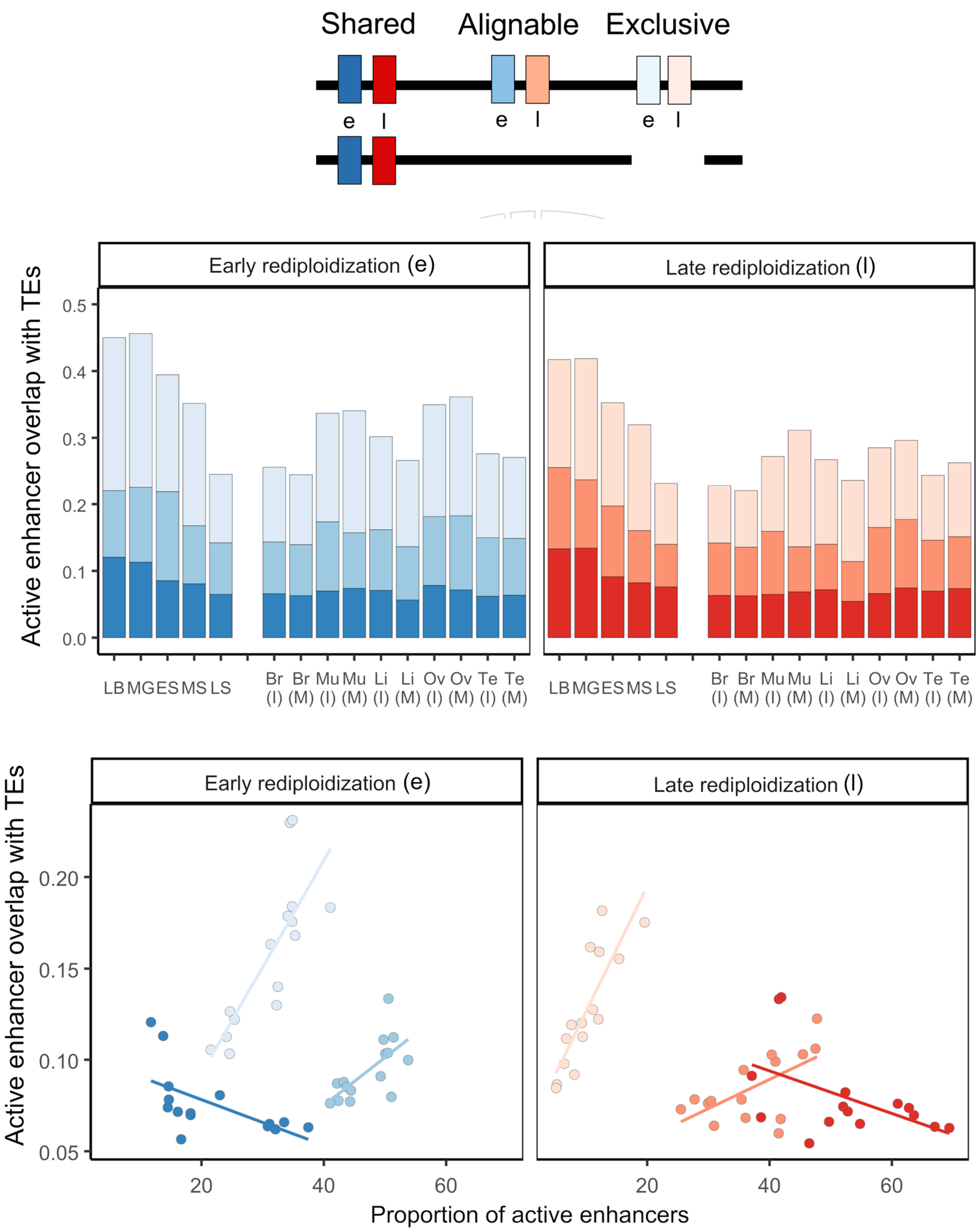

**Fig. S40. Ontogeny and tissue context constrain the overlap between TEs and active enhancers**. The top graph shows the percentage overlap of TEs across different DevMap and BodyMap sample types for each alignment category, and both rediploidization regions. The bottom graph visualises the correlation between percentage overlap of TEs vs. proportion of active enhancers across different DevMap and BodyMap sample types.

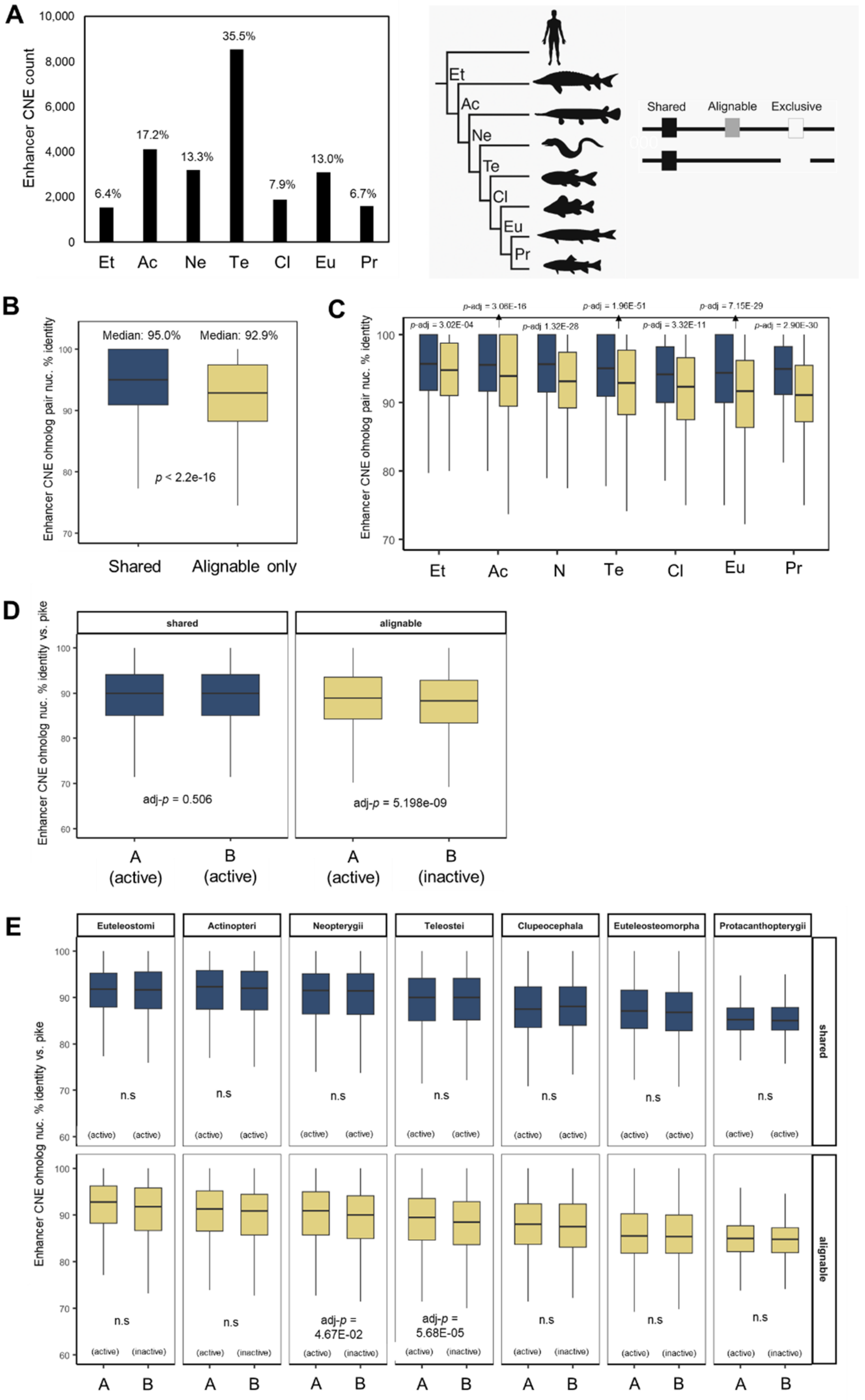

**Fig. S41. Overview of enhancer-CNEs evolution in the Atlantic salmon genome. (A) Counts of enhancer-CNEs stratified by phylogenetic age**. (**B** and **C**) Shared enhancer-CNEs, active at both ohnologs, show significantly higher sequence identity than Alignable only enhancer-CNEs, where enhancer activity is restricted to one ohnolog in a pair (Wilcoxon's rank test, done globally [**B**] and per CNE age category [**C**]). (**D** and **E**) Shared enhancer-CNE ohnologs from the same pairs show no difference in respective sequence-level divergence with their shared northern pike ortholog. However, for enhancer-CNEs in the Alignable category, the inactive enhancer-CNE ohnolog shows significantly greater sequence divergence with the pike ortholog, globally and in two CNE age categories (Wilcoxon's rank test, done globally [**D**] and per CNE age category [**E**]).

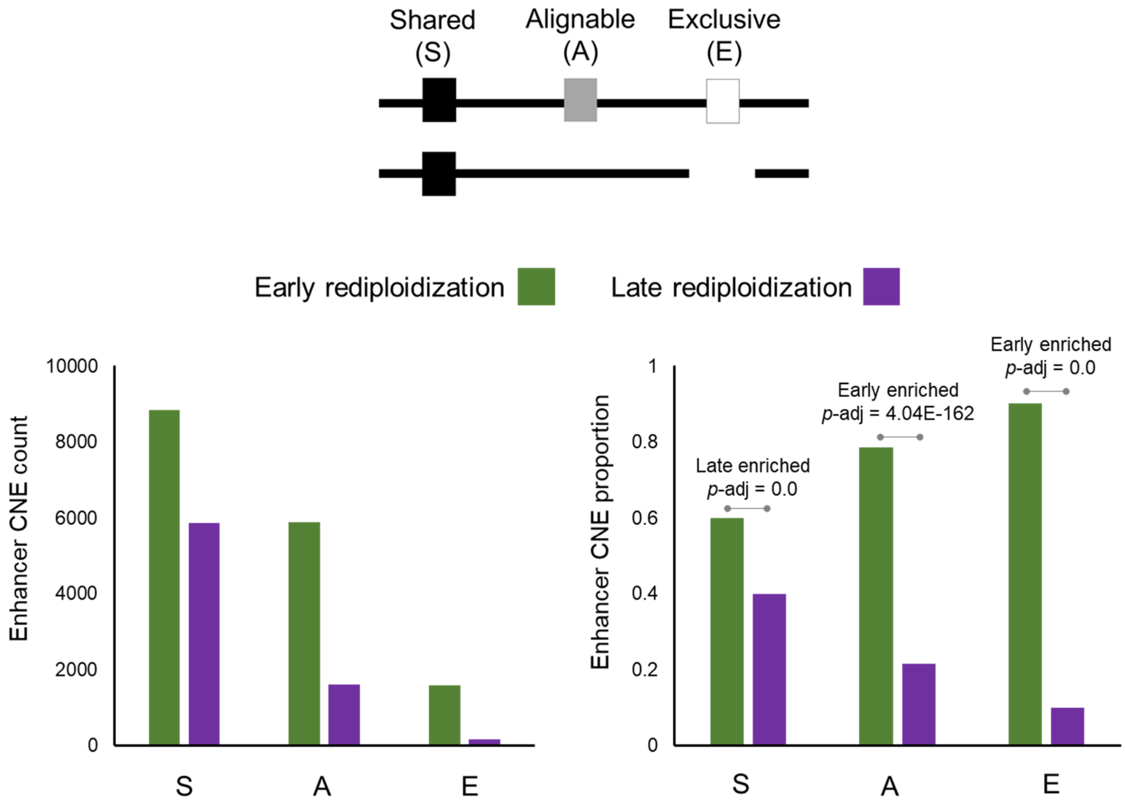

**Fig. S42. Enhancer-CNEs are differentially enriched in the Atlantic salmon genome dependent on rediploidization history**. Enhancer-CNE counts (left graph) and proportions (right graph) are shown for Early and Late rediploidization regions and each WGD-alignment category. The Shared enhancer-CNEs are significantly enriched in Late rediploidization regions, whereas the opposite is the case for Alignable only and Exclusive enhancer-CNEs.

### Supplementary Text

#### Text S1. Sampling Protocols for DevMap and BodyMap

*Executive summary*: This section provides the detailed protocols used for Atlantic salmon and rainbow trout samples taken: i) during embryogenesis (DevMap) and ii) from adult tissues of both fish sexes at two stages of maturation (BodyMap), for the purpose of RNA-seq ATAC-seq and ChIP-seq.

#### S1.1. DevMap sampling

##### S1.1.1. Materials and Reagents

- Petri dishes
- Watchmaker forceps
- 1.5 mL Eppendorf tubes
- Vortex
- Fixed angle centrifuge
- Ice tray
- Crushed ice
- Cold water from egg trays
- PBS

##### S1.1.2. Number of salmonid eggs required per stage

- Blastula: n=40
- Gastrula: n=30
- Early somitogenesis: n=20
- Mid-late somitogenesis: n=10
- Pharyngula stage: n=5

##### S1.1.3. Blastula to early gastrulation

*Note*: During blastulation to early stages of gastrulation, the cell mass is not tightly bound to the yolk and can be easily dissociated mechanically. However, it is also easy to lose or break cells. Because of this, the best approach is to dechorionate eggs under a dissection microscope and use a bore tip pipette to transfer each blastula to an Eppendorf tube.

1. Place the required number of eggs in a petri dish with de-chlorinated, aerated, cold water (from egg trays). Required number of eggs is stage dependent (above).
2. Place another petri dish with cold PBS under a dissection microscope.
3. Transfer one egg to the petri dish under the microscope.
4. Dechorionate using watchmaker forceps. Take care not to damage the blastula.
5. Use light mechanical pressure from the forceps to separate embryo material from yolk. For this step, a very careful and light scraping motion on the sides of the blastula works best.
6. Using a P1000 pipette set to 100 µL, with a bore pipette tip, transfer the embryo material to a 1.5 mL Eppendorf tube on ice.
7. Repeat for the required number of eggs for each replicate.
8. Let the embryo material decant to the bottom of the tube, then remove as much supernatant as possible and add 1000 µL of cold fresh PBS.
9. For samples meant for ATAC-seq library preparation, continue from step S3.1.3, for samples meant for crosslinking and subsequent ChIP-seq library preparation, continue

from step S4.1.

**S1.1.4. Mid-gastrulation onwards**

During gastrulation, normally at the onset of epiboly, cells will become more tightly bound to each other and to the yolk surface. This makes it more challenging to isolate embryos under the dissection microscope, but it also makes it easier to separate the gastrulas from the yolk without accidentally dissociating them. Because of this, eggs at this stage are dechorionated within a well of a six well plate, and vortexed to separate embryonic mass from yolk and oils.

1. Pre-chill swinging bucket centrifuge to 4ºC.
2. Place the required number of eggs for one biological replicate in the well of a six well plate, fill well with de-chlorinated, aerated, cold water (from egg trays). Place six well plate on ice.
3. Using watchmaker forceps, transfer one egg into another well, along with 1 mL of PBS.
4. Dechorionate egg and remove chorion, take care not to leave any embryonic material inside chorion.
5. Repeat for all eggs in one biological replicate.
6. Transfer contents of well into an 1.5 mL Eppendorf tube, on ice.
7. Vortex tube for ~10 secs.
8. Centrifuge on fixed angle centrifuge, 300g for 6 mins at 4ºC
9. Remove supernatant and add 1000 µL of fresh cold PBS.
10. Repeat the last centrifugation.
11. The samples are ready for cell isolation (S3.1.4) or crosslinking (S4.1), depending on downstream applications.

**S1.2. BodyMap sampling**

**S1.2.1. Materials and Reagents**

- One polystyrene box with ice, one with dry ice
- PBS plus (see buffers table)
- Labelled cryotubes or Eppendorf tubes
- Balance and weigh boats
- Scalpel and blades
- Tweezers and scissors
- Labelled Petri dishes
- 1 L of cold PBS
- Ethanol and distilled water to clean the tools
- Aluminum foil, kimwipes and absorbent paper
- Marker, pen and notepad
- Dietrich solution (histology)
- Air-tight containers (histology)

*Note*: Fish should be starved for 24 hr before sampling. Fish should be humanely killed using a Schedule 1 method before proceeding immediately with measurements and tissue sampling.

**S1.2.2. Tissue sampling**

1. Brain: make a transverse cut across the skull and lateral cuts (posterior to anterior) to create a flap which you can remove along with any skin, cartilage, or fat. Identify the

brain (A & B, directly below).

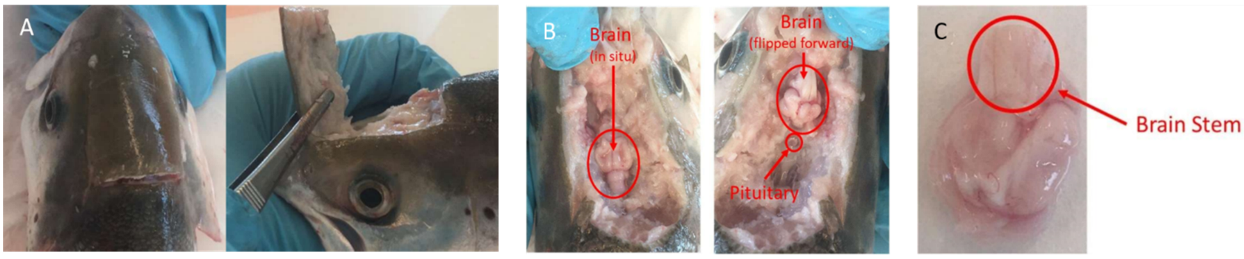

**Brain sampling.** (A) Access to the brain. (B) Sampling of the brain. (C) Removal of the brain stem

2. Carefully cut across the brain stem and transfer the whole brain to a petri dish placed on ice and containing sufficient PBS to rinse the tissue. Remove the brain stem and pituitary (B & C, directly above) and any contaminating tissue, then transfer to a clean petri dish.
3. Open the fish by using a shallow horizontal cut 5mm below the lateral line from the gill to the anal fin, and make two vertical cuts from the lateral incision to the belly. Cut through the skin and muscle but not so deep that you damage the internal organs (A, directly below).
4. Open the fish by flipping the muscle and skin layer down and stretching it to expose the internal organs (B, directly below).
5. Remove fat if necessary and expose the digestive organs (B, directly below). Then sample target organs in the following order: liver, gonad, distal intestine, head kidney, gill and skeletal muscle.

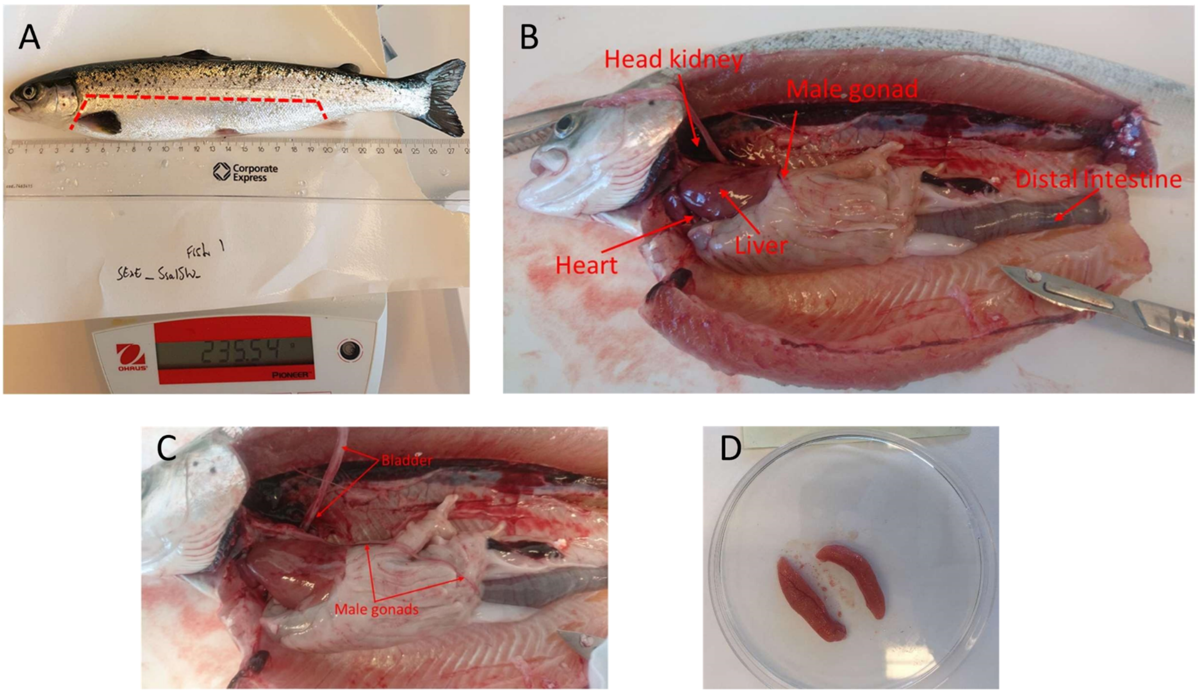

**Tissue sampling in an immature Atlantic salmon.** (A) lateral incision. (B) View of salmonid organs. (C) Immature male gonads. (D) Immature female gonads

6. Liver: Remove the intact liver after cutting the hepatic vein, and transfer to a clean petri dish on ice containing PBS. If necessary, remove a 1cm strip from the middle of the tissue

(in an anterior → posterior direction). Avoid or remove the vascular network and transfer the final trimmed sample to a sample petri dish.

*Note*: Because liver is an easy tissue to access and relatively large, it can be used as a source of DNA for whole genome sequencing of the individual. Collect at least 2 mg of tissue.

7. Gonads:

- In immature male_fish, testes are similar in appearance to the swim bladder (C, directly above), both are fine and delicate and look like sausage skins. Take a photo of the gonads *in situ*, then release the testes by cutting at the anterior end. The bladder lies over the testes and can be lifted to expose the underlying testes.
- In mature male fish, testes appear as large white sacks. Take a photo of the gonads *in situ*, then release the testes by cutting at the anterior end.
- In immature female fish, the ovaries are found under the digestive organs and appear as pinkish sacks (D, immediately above). Take a photo of the gonads *in situ*, then using tweezers, pull these away from the body cavity and cut at the point of attachment to liberate the organ.
- In mature female fish, the ovaries are large pink sacks and possibly contain eggs. Take a photo of the gonads *in situ*, then using tweezers, pull these away from the body cavity and cut at the point of attachment to liberate the organ.
- Gonad histology sampling: Remove sample(s) for histological analysis and place in appropriate histology petri dish ('H', directly below), then add PBS to remaining whole tissue (in collection dish; 'C') and histology sub-samples to prevent the tissue from drying out. If possible, wash away or press out and discard as many eggs/sperm as possible. The protocol for histology is provided in section S1.3 below.

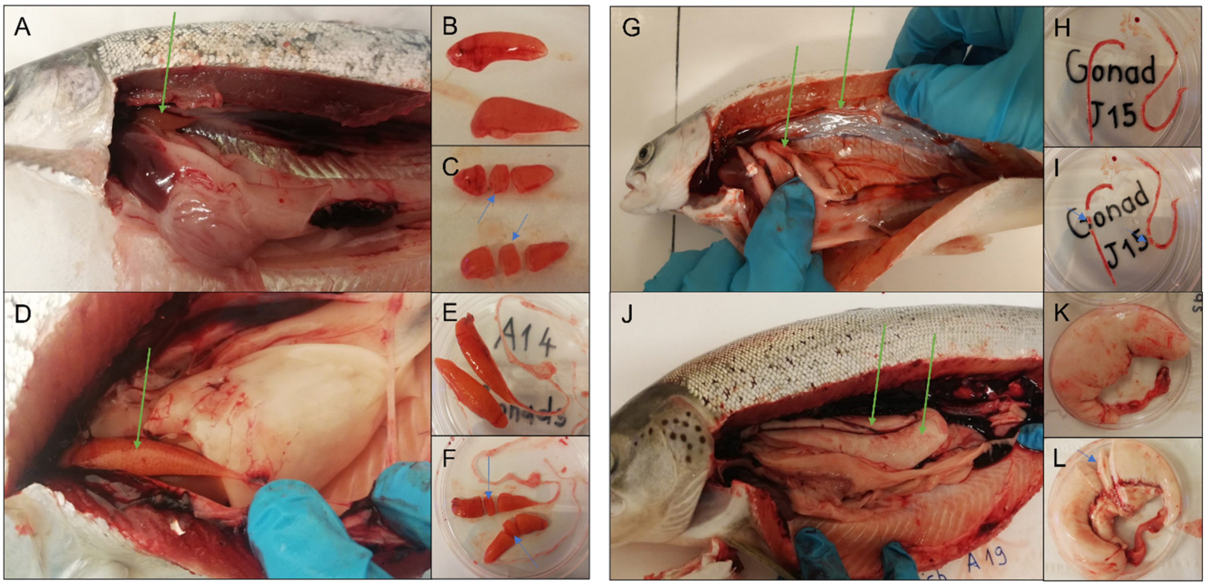

**Gonad sampling and sampling for histology.** (A to C) Immature female gonads. (D to F) Mature female gonads. (G to I) (A, D, G and J) green arrows show the location of the gonads. (B, E, H and K) Gonads after extraction. (C, F, I, L) blue arrows show the portion of the gonad sent for histology investigation in our study.

8. Distal intestine: Make a cut in the anterior part of the main digestive link (at the esophagus/ pharynx level) to release the beginning of the digestive tract. Lift the intestines out and locate the most terminal (distal) region proximal to the anus, cut to

release the tract completely.

9. Locate and isolate the distal intestine (DI). If feces are present, place the DI in a clean petri dish and gently use your finger to press them out. Transfer the DI to a petri dish containing PBS and cut it open along its length to expose its interior, then gently scrape with the back of a scalpel or spatula to remove mucus. Wash thoroughly in PBS, and transfer to the Sample petri dish (A & B, directly below).

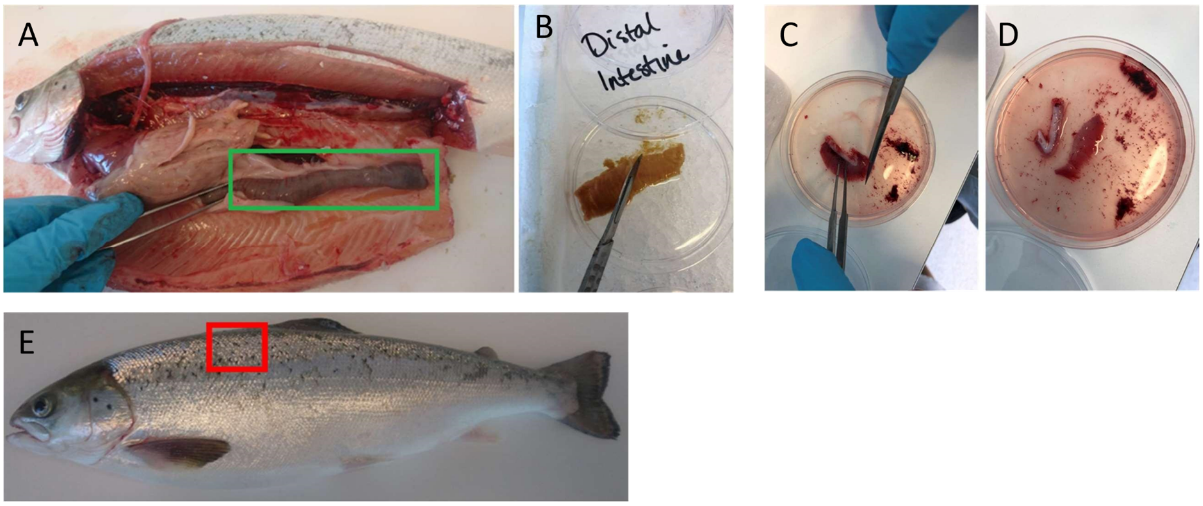

**Sampling images.** (A and B) Distal intestine sampling. (C and D) Gill sampling. (E) Localization of the muscle sampling

10. Head kidney: Remove and discard the heart to give better access to the head kidney, which is the most anterior portion of dark red tissue found under vertebral line and behind the head. The head/anterior kidney can be identified by its location and because it looks wider than the remaining more posterior kidney that follows the vertebral line along the body. The head kidney is fragile so care must be taken when extracting it. A small plastic spoon or spatula may be appropriate. Place directly into a sampling petri dish containing PBS.
11. Gill: Cut the gill at its root and immerse in a petri dish filled with PBS and scrape the surface to remove blood. Cut out and discard cartilaginous tissues (rakers) but retain the gill filaments and transfer these to a fresh petri dish with PBS (C & D, directly above).
12. Muscle: Sample a generous amount of fast-twitch myotomal muscle from the region anterior to the dorsal fin (E, directly above), avoiding the darker red muscle layer found close to the skin. Place into a corresponding petri dish and remove any skin and fat. Transfer cleaned muscle sample to petri dish and immerse in PBS.

*Note*: Frozen skeletal muscle is a relatively poor source of nuclei and the library protocols involve extensive filtering, which will return low yields of nuclei. For this reason, we recommend sampling at least 6g to ensure sufficient material.

#### S1.2.3. Preservation of tissues for RNA preservation

1. Place an unfolded piece of aluminum foil (no less than 10 x 10cm) or disposable weight-boat on a flat layer of dry ice ensuring good contact. At the same time, place two pairs of tweezers in dry ice, such that the ends of the tweezers become very cold. Then place pre-labelled cryotubes into dry ice so that they are cold when receiving samples.
2. Transfer tissue from sampling petri dishes to a new and dry petri dish. When possible, use

Kimwipes or equivalent to remove excess PBS. Then record the sample weight (subtracting the weight of the petri dish). Recommended sample collection weights (in mg) are shown for each tissue in the table immediately below.

| **Tissue** | **Mg to collect** | **Approx. µg total RNA per mg** |
|---|---|---|
| Brain | 15 | 1 |
| Distal Intestine | 15 | 1 |
| Gill | 15 | 1 |
| Gonad | 15 | 1 |
| Muscle | 100 | 0.25 |
| Liver | 15 | 3 |
| Head kidney | 15 | 2 |

3. With a sterile and clean scalpel, cut tissue samples into small pieces (approximately 2-3mm$^3$).
4. Using room temperature tweezers, drop tissue pieces of the recommended weights into 5-10 volumes RNAlater (or an equivalent reagent). See recommendations from the manufacturer, but we recommend storing the sample at 4°C overnight before transferring to -20°C for longer-term storage.
5. Next, using room temperature tweezers, drop all remaining tissue pieces onto the foil (if the tissue is sticky, this may require some aggressive flicking, or the use of a second pair of tweezers).
6. Tissue will freeze rapidly, try to place pieces separately on the foil and avoid clumping. After freezing the last tissue piece, use the chilled tweezers from step 2 to pluck each tissue piece from the foil and place in a pre-labeled chilled cryotube. If everything is kept cold, the small frozen pieces should not stick to each other or to the cryotube.

*Note*: the chilled tweezers will rapidly warm up once removed from dry ice. We recommend swapping between at least two pairs of tweezers to ensure only cold tweezers interact with samples.

7. Next, place the tubes in -80°C. This will allow you to later extract tissue without thawing the whole sample. Always work on dry ice and with cold tweezers for this later step.

#### S1.3. Gonadal sampling for histological analysis

##### S1.3.1. Materials & Reagents

- Dietrich's solution
- Small volume (15-20 mL) containers/tubes for storage of histological samples (e.g. https://www.fishersci.ca/shop/products/simport-scientific-securtainer-ii-tamper-evident-specimen-containers-5/p-4121042). These must be leak-proof.

**S1.3.2. Buffer recipes**

Dietrich's solution (1,020 mL, sufficient for approx. 50 samples):

- 600 mL dd$H_2O$
- 300 mL 95% ethanol
- 100 mL 40% formaldehyde
- 20 mL acetic acid

**S1.3.3. Protocol**

After excision, take a photo and weigh the entire gonads (section S1.2.2). If the gonad is large enough, prepare to sample three portions (anterior, middle and posterior), otherwise sample just the middle portion. Take another picture after sampling.

1. For histological sampling, make a clean cut using a sharp blade to ensure the integrity of the tissues and remove a section(s) not larger than $1cm^3$.
2. Place the samples in 10 volumes (at least) of Dietrich's solution for 10 days in sample containers. During this time, the samples should be kept cool and in the dark.
3. After 10 days, transfer the sample into new sample containers, then fill with 50% ethanol (at least 10 volumes) and let sit for 5 days.
4. After 5 days, samples are dehydrated through a graded series of ethanol and embedded in paraffin. Sections should be cut at a thickness of 4 µm using a microtome, mounted on slides, stained with haematoxylin and eosin and examined under photomicroscope with imaging capabilities (e.g. we used an Olympus Vanox, New York Microscope Company, Hicksville, NY, USA) to evaluate the maturity stage of the individuals (sexually immature or mature) by general morphology observation of the gametes development inside the slides.

Resulting histology images of each individual from our study is provided below for Atlantic salmon and rainbow trout.

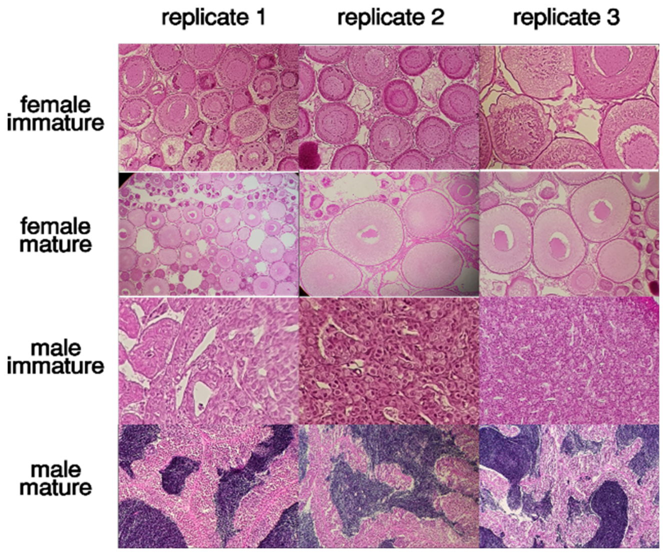

Histology images confirming the sexual maturation status of the 12 Atlantic salmon individuals used in our study.

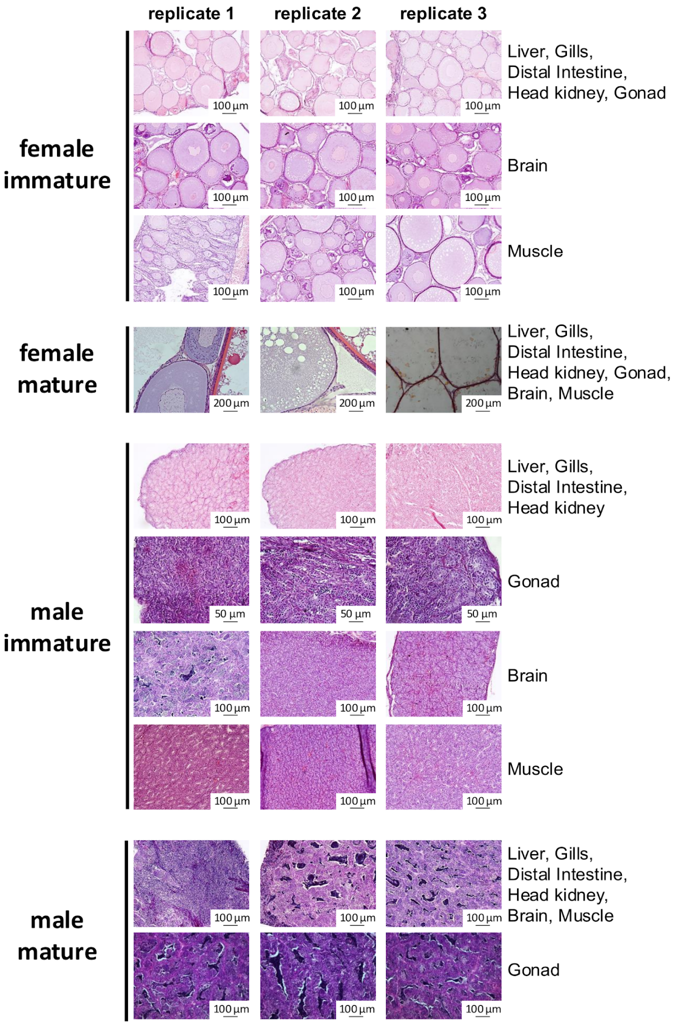

Histology images confirming the sexual maturation status of the 30 rainbow trout individuals used in our study.

### Text S2: DevMap and BodyMap RNA extraction protocols

Executive summary: This protocol describes the total RNA extraction methods used for Atlantic salmon and rainbow trout for DevMap and BodyMap studies.

#### S2.1. RNA extraction from salmonid embryos (DevMap)

##### S2.1.1. Materials and Reagents

- 1.5 mL Eppendorf tubes with safety lock
- Trizol (or equivalent)
- Sterile homogenization beads
- Tissuelyser or equivalent
- Water from egg trays
- Precipitation solution (recipe below)
- Isopropanol, anhydrous
- 1-Bromo-3-Chloropropane
- Nuclease-free water (NF-H2O)
- Ethanol 70%
- Sodium chloride (NaCl)
- Sodium citrate sesquihydrate
- RNAse-AWAY or equivalent
- MilliQ water

*Precipitation solution*: 100 mL of 1.2M NaCl and 0.8M sodium citrate sesquihydrate. To prepare, dissolve 7g NaCl in 50 mL NF-$H_2O$, then add 21.05 g sodium citrate sesquihydrate, then dissolve by heating and stirring. Filter through 0.2 µM syringe filter. Aliquot and store at -20ºC. Use only pre-autoclaved and RNA-AWAY washed bottles.

**S2.1.2. Protocol:**

1. Label required number of 1.5 mL Eppendorf tubes depending on the required number of biological replicates. No more than 3 eggs per tube should be used in salmonids.
2. Fill each tube with the required amount of homogenization beads.

*Note*: The amount of beads depends on the type of beads used. We successfully used 2x 3mm tungsten beads and 500 µL of both 1 mm Zirconium beads or 1 mm glass beads. It may be necessary to monitor the homogenate and use more beads/homogenization time if necessary.

3. In a flow hood, add 1 mL of TRIzol™ or equivalent to each tube. Store tubes in fridge while collecting embryos.

*Note*: The number of tubes used will depend on sampling strategy.

4. Place the required number of eggs for all biological replicates in water from egg trays within a Petri dish on ice.

*Note*: Using water from egg trays to temporarily keep the eggs while sampling allows for easy identification of dead/dying eggs, which will turn white and opaque and should not be sampled. If eggs are kept in PBS they will not turn white when they die, instead the chorion will turn more transparent and the yolk/embryo mass will clump into a ball. This makes it more difficult to tell between dead and alive eggs.

5. Using watchmaker forceps, transfer each egg to an empty petri dish, and puncture it with a pair of closed watchmaker forceps (pair 1), while holding the egg with a second pair of watchmaker forceps (pair 2). Do not remove forceps from the puncture site, instead, continue puncturing until pair 1 comes out through the other side.
6. Place egg over an open tube while still held by both pairs of forceps, and tear open the puncture by opening pair 1, dechorionating and depositing the egg in the tube at the same time.
7. Add no more than three punctured eggs per tube. Make sure tubes close tightly.

*Note*: this is to avoid overfilling of tubes due to the large egg size in salmonids. Egg puncturing is not necessary if using heavy 3mm tungsten beads.

8. Homogenize using a Tissuelyser at a setting of 30Hz for 6 mins.
9. Store tubes at -80ºC until RNA extraction or proceed with following steps.
10. Prepare (or thaw) the precipitation solution and label 3 additional Eppendorf tubes per sample.
11. Transfer the homogenate (without beads) to a new tube.
12. Centrifuge at 12,000g, for 10 mins at 4ºC.
13. Transfer supernatant to a new tube.

*Note*: A pellet will form at the bottom of the tube, while a layer of oil will form at the top. Extract only the middle layer between the pellet and the oil. If the oil is not visible, avoid aspirating trace amounts by keeping the pipette tip well below the surface.

14. Add 100 µL of 1-Bromo-3-Chloropropane.
15. Vortex vigorously for 15 sec. Incubate for 30 mins at room temperature.
16. Add 100 µL of previously prepared precipitation solution and 100 µL of anhydrous Isopropanol to each separate and empty tube.

*Note*: The precipitation solution and the isopropanol do not mix, hence one extra tube per sample is used here to avoid adding solution in incorrect proportions.

17. Centrifuge samples at 20,000g, for 15 mins at 4ºC.
18. Carefully place samples on ice without disturbing newly formed layers, and transfer 2x 200 µL of aqueous phase, slowly and carefully, into new empty 1.5 mL Eppendorf tubes. Use pipette tip with blunt end and keep tip in the middle, as far away from the interphase, the meniscus and the sides of the tube as possible. If there is debris stuck to the sides of the tube, stop pipetting before the meniscus level reaches the debris.
19. Add 200 µL of precipitation solution to each of the tubes where the aqueous layer was placed. Gently invert 6 times.
20. Incubate at room temperature for 10 mins or incubate overnight at -20ºC for increased RNA recovery.
21. Centrifuge samples at 20,000g for 10 mins at 4ºC.
22. Remove supernatant and resuspend in 1000 µL of 70% ethanol.
23. Incubate on rotation for 30 mins.
24. Repeat steps 21-23. Then, repeat step 21 again.
25. Remove supernatant, including remaining ethanol with a p-10 pipette.
26. Air dry pellets for 10 mins at room temperature.
27. Resuspend in 20-100 µL NF-H2O (depending on size of pellet).
28. Incubate at room temperature for 60 mins, flicking the tubes every 10 mins.
29. Assess RNA concentration using a Qubit fluorimeter
30. Dilute samples as necessary with NF-H2O, aiming for ~400 ng/µL.
31. Assess the quality of sample using Tapestation (to obtain RNA integrity numbers) and Nanodrop (to assess for purity).
32. Store your sample at -80ºC.

**S2.2. RNA extraction from Atlantic salmon tissues (BodyMap)**

A Qiagen column-based RNA extraction kit was used (miRNeasy Mini kit, #217004) following the manufacturer's recommendations, with an optional DNAse treatment and column drying as instructed by the manufacturer. Extracted RNA was stored at -80°C.

**S2.2. RNA extraction from rainbow trout tissues (BodyMap)**

*Summary*: In our study, rainbow trout RNA extraction followed the protocol of Chomczynski et al. (*104*). The tissues are ground into the TRIzol™ solution containing guanidium isothiocyanate, phenol and sodium acetate. After addition of chloroform, the samples are centrifuged to separate the total RNA (upper aqueous phase). The aqueous phase is extracted, and the total RNA is reconcentrated by conventional alcoholic precipitation.

**S2.2.1. Materials and Reagents**

- 2 mL Eppendorf tubes
- Precellys (Bertin Instruments, # P000669-PR240-A)
- TRIzol™: TriReagent (Euromedex, #TR118)
- Chloroform (Carlo Erba, #438603)
- Ethanol (Carlo Erba, #4146012)
- Isopropanol (Sigma, #33539)
- CK14 ceramic beads 1.4 mm, tubes 2.0 mL (#P000912-LYSK0-A.0 OZYME)
- 75% EtOH

**S2.2.1. Protocol:**

1. Place the tissue in a Precllyus-specific 2 mL grind tube with ceramic beads.
2. Add 1 mL TRIzol™ for each 200 mg tissue.
3. Grind tissue with Precellys (2 times 15 sec with 30 sec break).
4. Transfer the homogenized tissue to a 2 mL Eppendorf tube.
5. Add 200 µL chloroform (for 1 mL TRIzol™) and quickly mix by manual inversion for 15 sec (without vortex).
6. Incubate for 5-10 min at room temperature.
7. Centrifuge at 12 000g, for 15-30 mins at 4°C.
8. Collect the colourless upper phase and transfer it in a new Eppendorf tube.
9. Add isopropanol: 0.5 mL per 1 mL TRIzol™ and vortex quickly.
10. Incubate for 2 h at -20°C.
11. Centrifuge at 12,000g, for 25 mins at 4°C.
12. Remove the supernatant.
13. Wash the pellet with 1 mL of 75% EtOH (per 1 mL TRIzol™).
    13.1. Add 75% EtOH at the opposite wall of the pellet.
    13.2. Centrifuge at 15 000g, for 15 mins at 4°C.
    13.3. Remove carefully the supernatant.
14. Resuspend the pellet in 75% EtOH (0.5 mL per 1 mL TRIzol™ ).
    14.1 Add 75% EtOH at the opposite wall of the pellet.

14.2. Centrifuge at 12 000g, for 15 mins at 4°C.

14.3. Carefully remove the supernatant.

14.4. Spin down the tubes to remove the remaining supernatant.

14.5. Leave the lid open to let the pellet dry (2-5 mins).

15. Dissolve the pellet in RNAse-free $H_2O$.

16. After quality check by Nanodrop and Bioanalyzer, store at -80°C.

**Text S3: DevMap & BodyMap Omni-ATAC-seq**

*Executive summary*: This protocol describes how we performed the assay for transposase-accessible chromatin with sequencing (ATAC-seq) in embryos (DevMap) and frozen tissues (BodyMap), adapted from the Omni-ATAC method (*59*). It is divided into 3 main steps: i) nuclear preparation, ii) transposition and iii) PCR with size selection. Due to a diversity of input material, the first step (nuclear preparation) is different between DevMap and BodyMap, but also contains several tissue-specific modifications within BodyMap. All protocol alterations have been added below the main reference protocol.

Nuclear preparation for BodyMap consists of tissue disruption, filtration and homogenization, release of nuclei from cells, then purification and concentration of nuclei suspension via density gradient centrifugation. A different method of nuclei extraction was used for embryos, with ST and TST buffers developed previously for nuclei preparations (*104*). An aliquot of isolated nuclei is used to estimate nuclei concentration and to validate nuclei quality before transposition. The transposition reaction is sensitive to the ratio of transposase to nuclei used and requires prior optimizations depending on the sample type. Following transposition, the DNA is purified and frozen.

We performed the abovementioned steps in tissue-specific batches, then once a larger number of eluted DNA was accumulated, the samples were PCR-amplified and size-selected (180-700bp). It is important to assess by qPCR the optimal amount of PCR cycles for each tissue. Therefore, for all the PCR steps, we recommend using PCR tubes and not plates, then re-order them according to the amount of PCR cycles. Size selection is not required but often needed, either to remove high molecular weight fragments or left-over primers or primer dimers. The profile of each library was assessed by High Sensitivity Bioanalyzer or similar, with a recommended loading concentration of 1-2 ng/ µL.

**S3.1. Modifications to Omni-ATAC for DevMap**

*Notes*: The DevMap-specific modification starts with extracted embryos as input material, from which the cells are isolated (section S1.1). After the nuclei have been extracted from isolated cells (as described below), we recommend following the reference Omni-ATAC protocol from the transposition step onwards. There are also developmental-stage specific protocol adjustments, which have been divided into different sections for i) blastulation to early gastrulation stages, and ii) mid-gastrula to pharyngula stage. Additional comments within sections below describe any additional minor adjustments for closely related embryo stages.

**3.1.1. Materials and Reagents**

- 0.25% Trypsin-EDTA
- 15 mL Falcon tubes

- Rotator system
- Dissection microscope (optional)
- FBS (Heat inactivated)
- DMEM
- DMSO
- 40 µm filters (Thermo Fisher Scientific, catalog no. 08-771-2)
- 2 mL cryotubes
- Mr Frosty cooling chamber (Nalgene)
- Isopropyl alcohol

**3.1.2. Buffer recipes**

**<u>2x ST stock solution (Salt-Tris buffer, 10 mL)</u>**

- 146 mM of NaCl (292 µL)
- 10 mM of Tris-HCl pH 7.5 (100 µL)
- 1 mM of CaCl2 (10 µL)
- 21 mM of MgCl2 (210 µL)
- Nuclease-free water (9,388 µL)

*Notes*: Prepare the day before or fresh and keep on ice

**<u>1x ST solution (Salt-Tris buffer, 10 mL)</u>**

- 2x ST (5 mL)
- Nuclease-free water (5 mL)

**<u>TST working solution (10 mL)</u>**

5 mL of 2x ST stock solution

300 µL of 1% Tween-20

50 µL of 2% BSA

4.65 mL Nuclease-free water

1x proteinase inhibitor cocktail (PIC) tablet (Sigma-Aldrich, 11836170001)

*Note*: Prepare fresh and keep on ice

**3.1.3. Cell isolation for Blastulation to early gastrulation (follows end of section S1.1 above).**

1. Pipette the 900 µL suspension containing the extracted embryos up and down 10 times. Use a wide bore pipette tip and a p1000 pipette.
2. Vortex at a slow speed for 15 sec.

3. Pipette the 900 μL suspension containing the extracted embryos up and down 10 times. Use a wide bore pipette tip and a p1000 pipette.
4. Check solution is homogeneous with no clumps. If clumps are observed, repeat step 1.
5. Transfer solution to a 2 mL cryotube.
6. Add 10 μL of Trypan blue to a 1.5 mL Eppendorf tube.
7. Pipette sample up and down 3 times with a bore tip p1000 pipette set to 500 μL. Take a 10 μL aliquot for cell counting and add it to the tube with Trypan blue. Pipette up and down 3 times.

*Note*: This step is meant to homogenize the sample before taking an aliquot for cell counting, and only needs to be done with one replicate. Trypan blue stain needs an incubation period of 5 min; leave this tube for cell counting on ice while continuing to the next step with the sample tube.

8. Add 500ul of DMEM and 400ul of FBS to each sample tube.
9. Add 100 μL of DMSO. Invert tube 4 times. Perform this step as quickly as possible for all replicates at the same time to minimize cytotoxicity of DMSO.
10. Store in a pre-chilled Mr Frosty cooling chamber and transfer to a -80ºC freezer for up to 4h
11. Recover tube containing cells and Trypan blue. Assess cell numbers per mL and percentage of live cells using a hemocytometer.
12. Store cryopreserved cells at -80ºC.

**3.1.4. Cell isolation for mid-gastrula to pharyngula stage (follows endpoint of section S1.1 above).**

1. Remove supernatant from 900 μL PBS suspension containing extracted embryos and add required volume of cold Trypsin-EDTA indicated below. It may be required to use a 15 mL Falcon tube for the trypsin digestion when higher volumes of Trypsin EDTA are required:
    - Mid gastrula: 1,000 μL for 30 gastrulas (33 μL per gastrula)
    - Early somitogenesis: 1,000 μL for 20 embryos (50 μL per embryo)
    - Mid somitogenesis: 2,000 μL for 10 embryos (200 μL per embryo)
    - Late somitogenesis: 2,000 μL for 10 embryos (200 μL per embryo)
    - Pharyngula: 1,000 μL per embryo
2. Rotate for required time indicated below, 50RPM at room temperature:
    - Mid gastrula: 5 min
    - Early somitogenesis: 5 min, monitor digestion and increase time if necessary.
    - Mid somitogenesis: 10 min
    - Late somitogenesis 20 min; observe after 15min and decide if more time required.
    - Pharyngula: 30 min to 1h, with constant monitoring of digestion required. Observe tube against bright background to better observe clumps.
3. Use dissection microscope to estimate digestion progress. Go back to previous step if clumps are observed.

4. Add 500 µL of cold heat inactivated FBS per 1,000 µL of Trypsin EDTA

*Note*: If using a 15 mL Falcon tube for digestion, split homogenate into 1.5 mL Eppendorf tubes for fixed angle centrifugation.

5. Centrifuge 300g for 6 min at 4ºC.
6. Remove supernatant and resuspend in 1000 µL of PBS.
7. Centrifuge 300g for 6 min at 4ºC.
8. Remove supernatant and resuspend in 900 µL of cryopreservation solution without DMSO (45% DMEM, 55% FBS).
9. Pass homogenate through a 40 µm filter into a 2 mL cryotube.
10. Add 10 µL of Trypan blue to a 1.5 mL Eppendorf tube.
11. Pipette sample up and down 3 times with a bore tip p1000 pipette set to 500 µL. Take a 10 µL aliquot for cell counting and add it to the tube with Trypan blue. Pipette up and down 3 times.

*Note*: This step is meant to homogenize sample before taking an aliquot for cell counting and only needs to be done with one replicate. Trypan blue stain needs an incubation period of 5 min; leave this tube for cell counting on ice while continuing to the next step with the sample tube.

12. Add 100 µL of DMSO, close the cryotube and invert several times. Perform this step as quickly as possible for all replicates at the same time to minimize cytotoxicity of DMSO.
13. Place in Mr Frosty Cooling chamber and store at -80ºC for a minimum of 4 h.
14. Recover tube containing cells and Trypan blue. Assess cell numbers per mL and percentage of live cells using a haemocytometer.
15. Store cryopreserved samples at -80ºC.

#### 3.1.5. Nuclei extraction from isolated cells (all stages)

1. To 400 µL of cryopreserved cells, add up to 1 mL, then pipette up and down until defrosted.
2. Centrifuge at 500g for 5 min at 4°C.
3. Discard supernatant and resuspend pellet in 1 mL ice cold PBS + PIC.
4. Centrifuge at 500g for 5 min at 4°C. Discard supernatant and resuspend pellet in 1 mL ice cold TST solution.
5. Leave on ice for 5 min, pipetting gently up and down.
6. Filter through a 40 µm Falcon cell strainer using the rubber end of a syringe to mash the top of the filter and add 1 mL of TST to wash the well and filter.
7. Bring the volume up to 5 mL with 3 mL of 1X ST buffer.
8. Transfer the sample to a 15 mL conical tube and centrifuge in a swinging bucket centrifuge at 500g for 5 min at 4°C.
9. Resuspend the pellet in 1X ST buffer. Resuspension volume is dependent on the size of the pellet, usually within the range of 100-200 µL resuspended to 1 mL.

10. If nuclei clumping is observed, refilter through 40µm filter.
11. Count nuclei with hemocytometer.
12. Put forward 50,000 to 75,000 nuclei for the next step. Make up the solution to 1 mL with 1X ST buffer.
13. Centrifuge the nuclei aliquot at 800g for 10 min at 4ºC, discard the supernatant and add transposase reaction mix to the nuclei. Follow the BodyMap reference protocol (section 3.2) from the transposition step onwards but use 1.25 µL of transposase instead of 2.5 µL.

#### S3.2. Omni-ATAC-seq on frozen tissue (BodyMap)

##### S3.2.1. Materials and Reagents

- 40-60 mg of snap-frozen tissue (section S1.2)
- Milli-Q water
- Homogenization Buffer Unstable Solution, 1x (see recipe)
- 50%, 29%, 40% Iodixanol solution (see recipe)
- ATAC-RSB-Tween-20 (see recipe)
- Phosphate-buffered Saline Solution (PBS) (Sigma-Aldrich, P5368)
- Hoescht Dye Solution (ThermoFisher, 62249) or Trypan Blue (Sigma-Aldrich, T8154)
- Illumina Tagment DNA Enzyme (transposase) and Buffer (Illumina, 20034197)
- 1% (v/v) Digitonin, diluted in purified water (Promega G944A)
- 10% Tween-20 (Sigma-Aldrich, 11332465001)
- Nuclease-Free water (ThermoFisher, AM9937)
- MinElute® PCR Purification Kit (Qiagen, 28004)
- Qubit™ ds DNA HS Kit (ThermoFisher, Q32854)
- NEBNext® Ultra™ II Q5® Master Mix (NEB, M0544S)
- IDT for Illumina DNA/RNA UD Indexes Set (Illumina, 20027213)
- SYBR™ Green I Nucleic Acid Gel Stain, 25x, diluted in water (Invitrogen, S7563)
- Agencourt AMPure XP beads (Beckman Coulter, A63880)
- 80% (v/v) Ethanol (VWR, 20821)
- TET Buffer (see recipe)
- Bioanalyzer High Sensitivity Kit (Agilent Technologies, 5067-4626)
- 2 mL Tissue douncer (VWR, KT885303-0002)
- Pestle A for 2 mL tissue douncer (VWR, KT885301-0002)
- Pestle B for 2 mL tissue douncer (VWR, KT885302-0002)
- 70 µm cell strainer (Corning, 431751)
- 40 µm cell strainer (Corning, 431750)

- 50 mL Falcon tube (Sarstedt, 62.547.254)
- 5 mL Eppendorf Tube snap cap (VWR, 89429-306)
- Centrifuge with swinging bucket for 5 mL tubes and cooling option (Eppendorf, 5810R)
- 1.5 mL DNA LoBind Tube (VWR, 525-0130)
- Microcentrifuge (Beckman Coulter, 20R)
- Counting chamber BLAUBRAND® Thoma pattern (Sigma-Aldrich, BR718020)
- Inverted Microscope (Zeiss, AXIO vert.A1)
- Thermomixer (Eppendorf, 5382000015)
- Qubit 4 Fluorometer (Invitrogen, Q33226)
- Qubit™ assays tubes (ThermoFisher, Q32856)
- 0.2 mL PCR Tubes with Flat Caps (ThermoScientific, AB-0620)
- GeneAmp® PCR Machine (Applied Biosystems, 9700)
- Hard-Shell PCR 96-well Plates (BioRad, HSP9655)
- Plate Microseals® 'B' (BioRad, MSB1001)
- qPCR machine (BioRad, CFX96)
- Vortex Mixer (Gilson, 36110750)
- Magnetic rack (Life Technologies, 12321D)
- BioAnalyzer (Agilent Technologies, 2100)
- Mortar and pestle (Sigma-Aldrich, Z247464-1EA) (muscle-specific)

**S3.2.2. Buffer recipes**

**<u>Homogenization Buffer Unstable Solution, 1X (2 mL/sample)</u>**

- 6X Homogenization Buffer Unstable Solution (see recipe) (333.33 µL)
- 320 mM Sucrose solution (see recipe) (640 µL)
- 0.1 mM EDTA (Sigma-Aldrich, 03690-100 mL) (0.4 µL)
- 0.30% (v/v) of Nonidet P40 Substitute 10% (Roche, 11332473001) (60 µL)
- 0.5 mM Spermidine (Sigma-Aldrich, S2626) (1 µL)
- 0.15 mM Spermine (Sigma-Aldrich, S3256) (2 µL)
- Water (963.27 µL)

*Note*: Prepare fresh and keep on ice.

**<u>Homogenization Buffer Unstable Solution, 6X (910 µL/sample)</u>**

- 12X Protease Inhibitor Cocktail Solution (see recipe) (454.19 µL)
- 6X Homogenization Buffer Stable (454.19 µL)
- 0.1 mM PMSF (Sigma-Aldrich, 93482-50 ML-F) (1.51 µL)

- 1mM beta-mercaptoethanol (Sigma-Aldrich,M6250) (0.11 µL)

*Note*: Prepare fresh under fume hood and keep on ice.

**Sucrose solution, 1M**

- Sucrose dissolved in water and sterilized by filtration using a 0.2 µm filter (Sarstedt)

*Note*: Store at 4°C for up to several months.

**Protease Inhibitor Cocktail Solution, 12X (833 µL/sample)**

- EDTA-Free Protease Inhibitor Cocktail (PIC) tablets (Sigma-Aldrich, 11836170001), dissolved in 6X Homogenization Buffer Stable (833 µL)

*Note*: Prepare fresh and keep on ice.

**Homogenization Buffer Stable, 6X (100 mL)**

- 30 mM $CaCl_2$ (Sigma-Aldrich, C5670) (3 mL)
- 18 mM $Mg(Ac)_2$ (Sigma-Aldrich, 63052) (1.8 mL)
- 60 mM Tris-HCl pH 7.8 (Sigma-Aldrich, T2569) (6 mL)
- Water (89.2 mL)

*Note*: Store at room temperature for up to several months

**ATAC-RSB (50 mL)**

- 10 mM Tris-HCl pH 7.4 (Sigma-Aldrich, T2194) (500 µL)
- 10 mM NaCl (Sigma-Aldrich, S3014-500G) (100 µL)
- 3 mM $MgCl_2$ (Sigma-Aldrich, M1028) (150 µL)
- Water (49.25 mL)

*Note*: Store at 4°C for up to several months.

**TET Buffer (16.5 µL/sample, 100 samples)**

- 10 mM Tris-HCL pH 8.0 (Sigma-Aldrich, T2694) (16.5 µL)
- 1 mM EDTA pH 8.0 (Sigma-Aldrich, 03690) (3.3 µL)
- 0.05% (v/v) Tween-20 (Sigma-Aldrich,11332465001) (8.25 µL)
- Water (1621.95 µL)

*Note*: Store at 4°C for up to several months.

**50% Iodixanol solution (800 µL/sample)**

- 6X Homogenization Buffer Unstable Solution 1X (133.3 µL)
- 50% (v/v) Iodixanol OptiPrep (Sigma-Aldrich, D1556) (666.6 µL)

*Note*: Prepare fresh and keep on ice.

**29% Iodixanol solution (1200 µL/sample)**

- 6X Homogenization Buffer Unstable Solution 1X (200 µL)

- 160 mM Sucrose (see recipe) (192 µL)
- 29% (v/v) Iodixanol OptiPrep (Sigma-Aldrich, D1556) (580 µL)
- Water (228 µL)

*Note*: Prepare fresh and keep on ice.

**<u>40% Iodixanol solution (1200 µL/sample)</u>**

- 6X Homogenization Buffer Unstable Solution 1X (200 µL)
- 160mM Sucrose (see recipe) (192 µL)
- 40% (v/v) Iodixanol OptiPrep (Sigma-Aldrich, D1556) (800 µL)
- Water (8 µL)

*Note*: Prepare fresh and keep on ice.

**<u>ATAC-RSB-TWEEN (1 mL/sample)</u>**

- ATAC-RSB (see recipe), 1X (990 µL)
- 0.1% (v/v) Tween-20 (1 0 µL of 10% Tween-20)

*Note*: Prepare fresh and keep on ice.

**S3.2.3. Tissue disruption and nuclei isolation**

*Note*: To start, pre-chill the swinging bucket centrifuge and micro-centrifuge to 4°C, pre-heat the thermomixer to 68°C and pre-chill the mortar on dry ice (for muscle and gill only). Keep samples on ice while working.

1. Place frozen tissue into ice-cold 2 mL douncer containing 1.2 mL cold 1X homogenization buffer (1x HB).

*Note*: Allow frozen tissue to thaw completely in buffer (max. 5 min). The volume of 1x HB can be adjusted according to the amount of tissue (0.8-2.0 mL).

2. Dounce the tissue with pestle A and B (up to 5 strokes each depending on tissue type) until obtaining a homogenized solution.

*Note*: It is important to visually control the level of homogenization. Different people may apply different strengths during the process, so try to minimise associated impacts.

3. Pre-clear the solution by filtering it through a 70µm (or 40µm) cell strainer suspended in a 50 mL Falcon tube.
4. Transfer 400 µL of filtrated suspension to each of two new 5 mL Eppendorf tubes.

*Note*: When tissue is limited, adjust the initial volume of 1x HB so there are no leftovers of homogenate after splitting it in two 400 µL volumes at this stage

5. Add an equal volume (400 µL) of 50% iodixanol solution and mix by pipetting (to create a 25% mixture).
6. Take 600 µL of 29% iodixanol solution and slowly dispense it under the 25% mixture.

*Note*: Kim-tech or other fine paper can absorb excess drops of solution from the pipette tip before dispensing. A reverse pipetting technique can help avoid bubbles and mixing of layers. Slow motion of the pipette in and out of the tube can avoid abrupt volume displacement and mixing of layers.

7. Take 600 µL of 40% iodixanol solution and slowly dispense it under the 29% mixture.
8. Transfer 5 mL tubes to a pre-cooled (4°C) swinging bucket centrifuge and centrifuge at 3,220g for 25 min with brakes set to 'off'.

*Note*: Density gradient centrifugation is performed in a swinging bucket centrifuge with brakes off to avoid collapse of the gradient layers. Without brakes, the centrifuge needs an additional 8-9 min to stop. If possible, centrifugation at higher speed is better, as it may improve the purity of nuclei.

9. Collect the nuclei band which should be visible at the interface between 29%-40% iodixanol layers and transfer it to a common 1.5 mL LoBind Eppendorf tube.

*Note*: This can be done by careful removal of volume from the top layer (approximately 1250 µL until the nuclei band) or targeting directly the nuclei band.

10. Add 1000 µL of ATAC-RSB-Tween buffer to the tube containing nuclei and mix.
11. Centrifuge in a pre-cooled centrifuge 500g for 10min at 4°C.

*Note*: Note the orientation of the tube so that you can predict the location of the pellet.

12. Remove and discard the supernatant.
13. Resuspend the nuclei pellet in 50 µL of cold PBS.
14. Take an aliquot of 2 µL from nuclei suspension, add 8 µL PBS and mix with 10 µL of dye (Hoescht or Trypan Blue). Count nuclei using hemocytometer and assess the integrity of sample (a majority of the nuclei should be rounded to slightly oval, with intact membrane, showing no shrinkage or dilation, and without blebbing).

*Note*: If there is too much material to accurately count, prepare a new dilution but maintain the ratio of 10 µL diluted nuclei suspension (in PBS): 10 µL dye. If you observe a lot of tissue debris, add 100 µL PBS to your nuclei suspension, filtrate through 40 µm strainer and recount.

15. Dilute the appropriate volume of your original nuclei suspension in a new 1.5 mL tube with cold PBS to obtain 16.5 µL at 3 000 nuclei/µL (approx. 50 000 nuclei).

**S3.2.4. Transposase reaction and clean-up**

1. Prepare the following transpose reaction mix in a new tube:

| | |
|---|---|
| 25.0 µL | 2X TD buffer |
| 2.5 µL | Illumina Transposase |
| 0.5 µL | 1% Digitonin |

| | |
|---|---|
| 0.5 µL | 10% Tween-20 |
| 5.0 µL | Nuclease-free water |
| 33.5 µL | Total volume |

Immediately add 33.5 µL transposase reaction mix to 16.5 µL of nuclei suspension and mix by pipetting up and down 6 times.

2. Incubate the reaction at 37ºC for 30 min in a Thermomixer with 1000 RPM mixing. Then, proceed immediately with the next step.
3. Purify the resulting DNA fragments with MinElute PCR purification kit (follow manufacturer instructions).

*Note*: Eluted DNA can be stored at -20ºC until ready to amplify.

4. Measure DNA concentration with HS Qubit or equivalent.

**S3.2.5. Library amplification**

1. Thaw transposed DNA, indexes, NEBNext® Ultra™ II Q5® Master Mix and SYBRGreen on ice.
2. Prepare the following PCR reaction in PCR tubes.

| | |
|---|---|
| 5.0 µL | IDT for Illumina DNA/RNA UD Indexes |
| 25.0 µL | NEBNext® Ultra™ II Q5® Master Mix |
| 20.0 µL | Sample (transposed DNA) |
| 50.0 µL | Total volume |

3. Short-spin the PCR tubes to collect all the liquid to the bottom.
4. Run the 'Initial PCR amplification' program (choose the pre-heat option for lid and set to 100ºC).

| | |
|---|---|
| 72ºC for 5 min | x 1 |
| 98ºC for 30 sec | x 1 |
| 98ºC for 10 sec | x 5 |

| | |
|---|---|
| 63ºC for 30 sec | |
| 72ºC for 30 sec | |
| 4ºC hold | x 1 |

5. Set up 15 µL qPCR reactions in a qPCR plate to determine the appropriate number of additional cycles needed. Prepare a pre-mix of nuclease-free water, NEBNext® UltraTM II Q5® Master Mix and 25X SYBRGreen for n+2 samples (n samples + 1 NTC + 1 additional), distribute 9 µL in each well and then add IDT for Illumina DNA/RNA UD Indexes and the sample as per below:

| | |
|---|---|
| 3.76 µL | Nuclease-free water |
| 5.00 µL | NEBNext® Ultra™ II Q5® Master Mix |
| 0.24 µL | 25x SYBRGreen (diluted in water) |
| 5.00 µL | Sample (Initially amplified DNA or water for NTC) |
| 1.00 µL | IDT for Illumina DNA/RNA UD Indexes |
| 15.00 µL | Total volume |

6. Seal, vortex and centrifuge the qPCR plate 280g for 1 min at room temperature.
7. Run the qPCR as follows:

| | |
|---|---|
| 98ºC for 30 sec | x 1 |
| 98ºC for 10 sec | x 20 |
| 63ºC for 30 sec | |
| 72ºC for 1 minute | |
| 4ºC hold | x 1 |

8. Assess the amplification profiles and determine the required number of additional cycles to amplify. The number of cycles should equal 1⁄4 of the maximum achieved fluorescence. For each curve, subtract the baseline value from the end-point value. Then, divide by 4 and check the number of cycles corresponding to that value. If the 1⁄4 value is between two cycles, choose the lower one.

9. Short-spin the PCR tubes containing your Initially amplified DNA (45 µL) and run the 'Final PCR amplification' program (see below Table) without adding anything else.

*Note*: After final PCR amplification (done as per below table), keep samples at 4ºC until the size selection step. It is possible to assess the profile of the libraries before that. To do so, use High Sensitivity Qubit kit to measure the concentration then dilute a 1 µL aliquot of your PCR product for Bioanalyzer profiling.

| | |
|---|---|
| 98ºC for 30 sec | x 1 |
| 98ºC for 10 sec | # of cycles determined individually for each sample by qPCR |
| 63ºC for 30 sec | |
| 72ºC for 1 minute | |
| 4ºC hold | x 1 |

**3.2.6. Library size selection**

1. Thoroughly resuspend Ampure XP beads by vortexing for at least 1 min.
2. Add 0.53x volume of beads (i.e. 23.9 µL assuming 45 µL reaction) to the sample and mix by pipetting (avoiding bubbles).
3. Incubate at room temperature for 10 min and mix gently every 5 min.
4. Touch-spin the tubes and place them on the magnetic rack for 5 min (until the supernatant is clear of beads).
5. Transfer the supernatant to a new tube and add another 1.3x original volume of Ampure XP beads (58.5 µL assuming 45 µL reaction).
6. Mix well by pipetting, avoiding bubbles.
7. Incubate at room temperature for 15 min while mixing gently every 5min.
8. Place on the magnetic rack for 5 min (until the supernatant is clear of beads).
9. Remove and discard supernatant.
10. While the tubes are still on the magnet, and without disturbing the pellet, wash the beads with 100 µL of freshly prepared 80% ethanol. Remove ethanol and discard. Repeat this step again.
11. Remove the tubes from the magnet, touch spin the samples, place back on the magnet and carefully remove the last traces of ethanol.
12. Remove the samples from the magnet and allow the tubes to air dry with caps open (30 sec to 2 min, do not wait until you see small cracks in the beads).
13. Add 16.5 µL TET Buffer. Resuspend beads by pipetting.
14. Rehydrate at room temperature for a minimum of 2 min.
15. Place on the magnetic rack for 5 min (until the supernatant is clear of beads).
16. Transfer the supernatant (eluted DNA) to a LoBind Eppendorf tube or 96 well plate.

17. Measure DNA library concentration with Qubit High Sensitivity Kit.

18. Validate DNA fragment size with Bioanalyzer or Tapestation High Sensitivity DNA kit

*Note*: The DNA fragment size distribution should follow a nucleosome pattern (ca. 200bp-350bp-500bp peaks for libraries), like in the example provided immediately to the right. If you have an excess of primer dimers (narrow peak around 120 bp), do an additional clean up with MinElute PCR purification kit and elute in 10 µL. If you have an excess of long fragments in your samples (>1000 bp), do an additional round of bead purification.

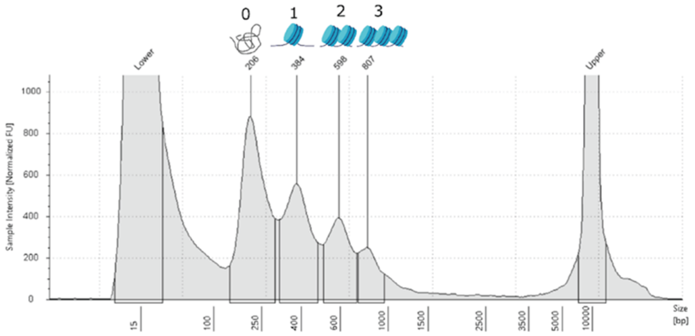

#### S3.2.6. Tissue-specific modifications

- *Brain, liver and immature gonads*: Follow reference protocol. Recommended to use 40-50mg total of pieces for liver or immature gonad and 20-30 mg for brain. No filtering is needed after homogenization.
- *Muscle*: Follow the reference protocol. The initial amount of tissue should be larger than for other tissues. Start with minimum 100 mg total of pieces. Larger amount of tissue also requires a larger volume of homogenization buffer (1xHB). Start with minimum 1.6 mL of 1xHB. Grind the tissue on a mortar previously chilled on dry ice until it is powdery or 'corn-flake' looking. Filter through 70 µm cell strainer and/or through 40µm if debris are still visible.
- *Mature male gonad*: Buffer preparation needs larger volumes of 1x HB (recommendation to prepare 9 mL more per sample) and 50% iodixanol solution (500µL more per sample) than reference protocol, due to additional washing steps and need for a second iodixanol centrifugation. 150-200 mg is a suitable weight of tissue. Recommended modifications as follows:

    1. Place the tissue in a 5 mL tube and add 1 mL of 1x HB. Gently squeeze the tissue with a pestle against the tube side to release the sperm remains, but do not break the tissue into smaller pieces.
    2. Vortex for at least 30 sec and pass through a 40 µm cell strainer suspended in a 50 mL Falcon tube. Add 200 µL of 1x HB over the strainer to rinse the tissue and place the tissue back to the 5 mL tube. Discard the filtrate containing mostly sperm (you can control the content of the filtrate by microscopy with Trypan blue coloration).
    3. Repeat steps 2 and 3 until the solution containing the tissue is clear.
    4. Cut the tissue into pieces and place it into an ice-cold 2 mL douncer containing 1 mL cold 1x HB. Homogenize as per reference protocol (step 2, section S3.2.3).
    5. Filter through a 70 µm cell strainer suspended in a 50 mL Falcon tube.
    6. Perform gradient centrifugation and recover nuclei band as per reference protocol (step 3-9, section S3.2.3).

7. Prepare a second gradient centrifugation adding an equal volume of 1x HB to the collected nuclei band (400 µL) resulting in a 15-20% iodixanol suspension and add carefully (avoid mixing layers) 500 µL of 50% iodixanol solution to the base of the 2 mL tube.
8. Centrifuge samples at 4ºC at 12,000g for 40 min (using soft stop).
9. Collect the nuclei band (around 100-150 µL) found between the top (15-20% iodixanol) and bottom (50% iodixanol) layers. Do not collect the pellet found at the bottom of the tube, which mostly contains sperm.
10. Continue as per reference protocol

**Text S4. DevMap & BodyMap ChIP-seq**

*Executive summary*: This protocol describes how we performed Chromatin Immunoprecipitation with Sequencing (ChIP-seq) in embryos and frozen tissues in Atlantic salmon and rainbow trout.

Due to a diversity of input material, two methods were required. For all samples with low input material (DevMap samples and brain samples from BodyMap of rainbow trout), a µChIPmentation Kit for Histones (Diagenode #C01011011) was used together with Chromatin EasyShear Kit-High SDS (Diagenode # C01020012) with a few modifications to the manufacturer's recommended steps (as described below). For all the remaining BodyMap samples, a classic custom ChIP protocol was developed along with a V3 MicroPlex Library Preparation Kit (Diagenode; #C05010002). This protocol was optimized for each tissue separately. Hence, a reference BodyMap protocol is described below (as applied for liver tissue), followed by a list of the tissue-specific modifications applied. The same antibodies were used across all assays for the immunoprecipitation step. For both ChIP methods used, the input controls (i.e. lacking immunoprecipitation steps) from embryos from the same developmental stage (DevMap) or from the same tissue/sex/maturity (BodyMap) were equimolarly pooled before library preparation for sequencing.

**S4.1. Crosslinking of embryos (DevMap)**

This protocol describes a method used to crosslink proteins to DNA of whole salmonid embryos, with the purpose of subsequent ChIP-seq library preparation.

**S4.1.1. Materials and Reagents**

- 4% PFA solution in PBS (Thermo-Fisher #30525-89-4)
- Rotator wheel with 1.5ml conical tube inserts (such as IKA™ 0004015000)
- 1M glycine solution (Sigma Aldrich #67419)
- PBS (Gibco™ 10010023)
- Fixed angle centrifuge

**Steps:**

1. Pre-chill fixed angle centrifuge

2. Prepare chilled 1% PFA solution in PBS by diluting 4% stock PFA solution 1:3 in PBS. 875µL per tube will be required.

2. Starting from S1.1.3 (step 9) or S1.1.4 (step 11). Remove supernatant and resuspend in 875 µL of 1% PFA.

3. Put tubes on a rotator at 50 RPM for X min, at room temperature.

*Note: X min = 8 mins for blastula and gastrula, 10 mins during segmentation stages, 15 mins post-segmentation stages.*

5. Add 125 µL of 1 M glycine. Continue rotating for an additional 10 mins at 50 RPM, at room temperature.
6. Centrifuge at 300 RCF, for 6 min at 4ºC.
7. Discard supernatant and resuspend in PBS, then store at -80ºC

**S4.2. ChIP-seq on crosslinked embryos using µChIPmentation (DevMap)**

**S4.2.1. Materials and Reagents**

- µChIPmentation Kit for Histones (Diagenode #C01011011)
- Chromatin EasyShear Kit - High SDS (Diagenode # C01020012
- 24 unique dual indexes (UDI) for tagmented libraries Set I and Set II from Diagenode (#C0101134 and #C0101135)

Steps:

1. Thaw crosslinked embryo material on ice.
2. Wash once with 1 mL of ice-cold PBS.
3. Spin down. Remove PBS. Homogenize using 2 mL dounce with 1 mL homogenization buffer (PBS + 1x PIC, 20-30 pestle strokes).
4. Perform the cell lysis with Chromatin EasyShear Kit-High SDS, using 500,000 homogenized cells for 315 µL of tL1 lysis buffer with 1x PIC on ice for 20-30min.
5. Follow the manufacturer's recommendations of µChIPmentation Kit for Histones kit with following parameters of sonication: 500 µL, 3-7 cycles (1 cycle = 30sec ON & 30sec OFF), 0.65 mL Bioruptor® Microtubes (#C30010011) and with following modifications:
    - immunoprecipitation step performed with 100 ng of chromatin (37,000 cells) and 0.8-1 µg of antibody per IP
    - Tagmentation of immunoprecipitated samples: 37°C for 12 min, PCR amplification: 14-15 cycles

**S4.2.2. Modifications of the manufacturer's protocol that apply only to the input sample:**

6. Use 6 µL for input sample + 21 µL of ChIPmentation mix (1µL Tn5 + 19 µL tagmentation buffer + 1µL $MgCl_2$)
7. Mix with Protein A beads pellet (equivalent to 10 µL of washed Protein A beads)
8. Tagment for 15-17 min
9. Add 25 µL of 2x High Fidelity mix, continue with repair and reverse crosslinking as in the protocol

10. Magnetise beads and transfer supernatant in a new tube
11. Add 2 µL of diluted UDI primer pair
12. PCR using 14-15 cycles and purify
13. Purify with 1x volume AMPure beads (to remove the excess of adapters)

**S4.3. ChIP-seq on frozen tissue (BodyMap)**

**S4.3.1. Materials and Reagents**

- Snap frozen tissue (see tissue-specific protocol modifications)
- PBS Plus (see recipe below)
- Hoescht Dye Solution (ThermoFisher, 62249) or Trypan Blue (Sigma-Aldrich, T8154)
- 37% (v/v) Formaldehyde solution (Sigma-Aldrich, 252549)
- 1M Glycine (w.v) (Sigma-Aldrich, 50048)
- Complete Sonication Buffer (see recipe below)
- Complete IP Buffer (see recipe below)
- Elution Buffer (see recipe below)
- RNase A (Qiagen, 19101)
- Proteinase K (Qiagen, 19133)
- 5 M NaCl (Sigma-Aldrich, S3014)
- MinElute® PCR Purification Kit (Qiagen, 28004)
- Qubit™ ds DNA BR kit (ThermoFisher, Q32853)
- Bioanalyzer DNA 1000 kit (Agilent Technologies, 5067-1504)
- Protein A Dynabeads (Life Technologies, 10004D)
- Protein G Dynabeads (Life Technologies, 100002D)
- Antibodies H3K4me1(Diagenode, C15410194)
- Antibodies H3K4me3 (Diagenode, C15410003)
- Antibodies H2K27ac (Diagenode, C15410196)
- Antibodies H3K27me3 (Diagenode, C15410195)
- Recovery Buffer (see recipe below)
- Complete Low Salt Buffer (see recipe below)
- Complete High Salt Buffer (see recipe below)
- 20% SDS (Fluka Analytical, 05030-1L-F)
- Low TE buffer (see recipe below)
- Qubit™ ds DNA HS Kit (ThermoFisher, Q32854)
- Microplex Library Preparation Kit v3 (Diagenode, C05010001)

- 60% Iodixanol OptiPrep (Sigma-Aldrich, D1556)
- 50%, 29%, 40% Iodixanol solution (see recipe below)
- Dounce Tissue Grinder Tube of 15 mL and Pestles (A & B) (VWR, 102095-542)
- 5 mL Eppendorf Tube snap cap (VWR, 89429-306)
- Counting chamber BLAUBRAND® Thoma pattern (Sigma-Aldrich, BR718020)
- Inverted Microscope (Zeiss, AXIO vert.A1)
- Centrifuge with swinging bucket for 5 mL tubes and cooling option (Eppendorf, 5810R)
- HulaMixer™ (Sigma-Aldrich, 15920D)
- Vortex Mixer (Gilson, 36110750)
- Micro-centrifuge (Beckman Coulter, 20R)
- 1.5 mL DNA LoBind Tube (VWR, 525-0130)
- Corning™ CoolRack™ (Fisher Scientific, C432034)
- EpiShear Probe Sonicator (Active Motif, 53052)
- Thermomixer (Eppendorf, 5382000015)
- Qubit 4 Fluorometer (Invitrogen, Q33226)
- Qubit™ assays tubes (ThermoFisher, Q32856)
- BioAnalyzer (Agilent Technologies, 2100)
- Magnetic rack (Life Technologies, 12321D)
- 2.0 mL DNA LoBind Tube (VWR, 525-0131)
- Mortar and pestle (Sigma-Aldrich, Z247464-1EA)

**S4.3.2. Buffer recipes**

**PBS Plus**

- PBS (Sigma-Aldrich, P5368), 1x
- PIC (Sigma-Aldrich, 11836170001), 1x
- 0.1 mM PMSF
- 5 mM Sodium Butyrate

*Note*: Prepare fresh under fume hood and keep on ice

**Sonication Buffer**

- 50 mM Tris-HCl pH 8.0
- 10 mM EDTA
- 1% (v/v) SDS
- Water

*Note*: Store at room temperature for up to 1 month. Complete the buffer for use with 7x PIC, 1 mM PMSF and 12.5 mM Sodium Butyrate (under a fume hood, while keeping on ice). If precipitation of SDS occurs, warm and agitate a little.

**IP Buffer**

- 20 mM Tris-HCl pH 8.0 (Sigma-Aldrich
- 2 mM EDTA pH 8.0 (Sigma-Aldrich, 03690)
- 0.1% (v/v) Triton X-100
- 150 mM NaCl
- Water

*Note*: Store at room temperature for up to 3 months. Complete the buffer for use with 1x PIC, 1 mM PMSF and 1 mM Sodium Butyrate (under a fume hood, while keeping on ice).

**Elution Buffer**

- 10 mM Tris-HCl pH 8.0 (Sigma-Aldrich
- Water

*Note*: Store at room temperature

**Recovery Buffer**

- 50 mM Tris-HCl pH 7.5 (Sigma-Aldrich
- 5 mM EDTA pH 8.0 (Sigma-Aldrich, 03690)
- 1% (v/v) SDS
- 50 mM NaCl
- Water

*Note*: Store at room temperature for up to 3 months. Complete the buffer for use with 0.1 mM PMSF (under a fume hood, while keeping on ice).

**Low Salt Wash Solution**

- 20 mM Tris-HCl pH 8.0 (Sigma-Aldrich
- 2 mM EDTA pH 8.0 (Sigma-Aldrich, 03690)
- 150 mM NaCl
- 1% (v/v) Triton X-100
- 0.1% (v/v) SDS
- Water

*Note*: Store at room temperature for up to 3 months. Complete the buffer for use with 1 mM Sodium Butyrate (under a fume hood, while keeping on ice).

**High Salt Wash Solution**

- 20 mM Tris-HCl pH 8.0 (Sigma-Aldrich

- 2 mM EDTA pH 8.0 (Sigma-Aldrich, 03690)
- 500 mM NaCl
- 1% (v/v) Triton X-100
- 0.1% (v/v) SDS
- Water

*Note*: Store at 4°C temperature for up to 3 months Complete the buffer for use with 1 mM Sodium Butyrate (under a fume hood, while keeping on ice).

**Low TE Buffer**

- 10 mM Tris-HCl pH 8.0 (Sigma-Aldrich
- 1 mM EDTA pH 8.0 (Sigma-Aldrich, 03690)
- Water

*Note*: Store at room temperature.

**S4.3.3. Tissue disruption, crosslinking and sonication**

*Note*: Pre-chill swinging bucket centrifuge and microcentrifuge to 4°C. Keep samples on ice

1. Place the frozen tissue (here assumes liver, but see tissue-specific modifications, section S4.3) into an ice-cold 15 mL douncer containing 5 mL cold PBS Plus.

*Note*: Allow the frozen tissue to thaw completely in the buffer.

2. Dounce the tissue with pestle A and B (approx. 5 strokes each) until obtaining a homogenized solution.

*Note*: It is important to carefully control the level of homogenization. Note that different people may apply different strength during the process.

3. Transfer the homogenate into a 5 mL Eppendorf tube.
4. Rinse the dounce and pestles with 0.5 mL of PBS Plus and add to the previous solution.

*Note*: A pipette tip cut at the bottom can be used to facilitate the aspiration.

5. Take an aliquot of 10 µL from the suspension and mix with 10 µL of dye (Hoescht or Trypan Blue). Count nuclei manually using a hemocytometer.
6. Spin in a pre-cooled centrifuge at 2,500g for 3 min at 4°C.
7. Remove and discard the supernatant.
8. Resuspend the pellet in 2.43 mL of PBS Plus buffer and add 67.5 µL of 37% formaldehyde solution at room temperature. Mix well.

*Note*: If more than 24 million nuclei, resuspend in 4.86 mL of PBS Plus buffer then add 135 µL of 37% formaldehyde solution and split into two tubes.

9. Rotate (360° rotation, 40 rpm) at room temperature for 5 min.

10. Quench the reaction with 360 µL of glycine 1M (0.125M final) and place back in the mixer for 10 min.
11. Centrifuge at 800g for 5 min at 4°C.
12. Remove the supernatant containing formaldehyde.
13. Resuspend with 4 mL of PBS Plus buffer by pipetting (applying brief vortex if necessary).

*Note*: If more than 24 million nuclei, resuspend each pellet in 2 mL of PBS Plus and then merge.

14. Centrifuge 800g for 5 min at 4°C.
15. Remove and eliminate the supernatant.
16. Resuspend with 5 mL of PBS Plus buffer by pipetting (applying brief vortex if necessary).
17. Centrifuge 800g for 5 min at 4°C.
18. Remove and eliminate the supernatant. Resuspend with 1 mL of PBS Plus buffer and transfer to a clean 1.5 mL Eppendorf tube.
19. Centrifuge on a pre-cooled micro-centrifuge 3000g for 10 min at 4°C.
20. Remove the supernatant.

*Note*: The nuclei pellet can be snap frozen and stored in -80ºC for up to 2 months.

21. Resuspend the pellet by pipetting in a ratio of 100 µL of complete sonication buffer per 1 million nuclei. Incubate on ice for 5 min.
22. Sonicate the chromatin while keeping the samples cold with the following settings:

| | |
|---|---|
| Number of cycles | 16 |
| Amplitude | 55 % |
| Cycle interval | 1 second 'ON' / 1 second 'OFF' |
| Cycle time | 30 sec |

*Note*: Each cycle = 30 sec total, composed by a repetition of 1 sec 'ON' / 1 sec 'OFF'. The total 'ON' period is 15 sec. These parameters are specific to the sonication device EpiShear Probe Sonicator (Active Motif, 53052), used for Atlantic salmon.

23. Centrifuge the sonicated chromatin 8,000g for 10 min at 4°C.
24. Transfer the supernatant into a new 1.5 mL Eppendorf tube. Record the volume.

*Note*: If sonication done in multiple tubes, keep separate until quality control for each is performed.

25. Use an aliquot of 20 µL per tube for quality control.
26. Add 3x volumes of complete IP buffer to the remaining chromatin.

27. Store the diluted chromatin at -80°C (safe stop point).

#### S4.3.4. Quality control of sonication

28. Add 67 µL of elution buffer, 2 µL of RNase A, 5 µL of proteinase K and 6 µL of NaCl to the 20 µL aliquot.
29. Incubate in thermomixer at 500 rpm for 1h 30 min at 68℃.
30. Purify using MinElute purification kit and elute in 20 µL of elution buffer.
31. Record chromatin concentration using Qubit Broad Range kit using 1µL of eluate.
32. Validate the fragments size distribution with Bioanalyzer DNA 1000 kit or equivalent (e.g. Tapestation)

*Note*: A good quality sonication profile has a minimum of 70% of the chromatin in a size distribution of 100-600 bp centered around 350 bp. Later, you can pool together sample tubes with this expected fragment distribution to run the immunoprecipitation.

#### S4.3.5. Beads preparation

*Note*: The workflow below is intended for 4 immunoprecipitations with 5 µg of material each and 50 ng for input control, from sonicated chromatin sample with the following parameters: 400 µL volume before 3x dilution (step 25), Qubit-based concentration of aliquot (step 30) = 60 ng/µL.

33. Clean the beads by mixing 528µL of protein A beads, 528µL of protein G beads and 4576µL of complete IP buffer and split by 1276 µL in four LoBind 2 mL tubes.

*Note*: ratio 1:1 of protein A beads and protein B beads, and ratio 60 µL of beads for 260 µL of IP buffer. 10 % is added to avoid repetition of the beads cleaning in case of loss during washing and pipetting.

34. Mix by pipetting and/or vortexing and touch spin the tubes.
35. Place the tubes in the magnetic rack for a couple of min (or until the supernatant is clear of beads) and discard the supernatant.
36. Remove the tubes from the magnetic rack and add 1144 µL of complete IP buffer.
37. Repeat 2 more times the steps 34-36.

#### S4.3.6. Coupling of beads and antibodies

38. As per below table, mix the cleaned beads with each antibody in a separate 1.5 mL tube and rotate (360° rotation, 40 rpm for 30 min at room temperature).

| | |
|---|---|
| 130 µl of beads with | 0.7µl (1.05µg) of H3K4me1 |
| 130 µl of beads with | 0.8µl (1.12µg) of H3K4me3 |
| 130 µl of beads with | 0.8µl (2.24µg) of H3K27ac |
| 130 µl of beads with | 1.9µl (2.09µg) of H3K27me3 |

#### S4.3.7. Chromatin pre-clearing

39. Place 520 µL of remaining cleaned beads in a 2 mL tube in the magnetic rack for 2 min and discard supernatant.

*Note:* If the sonicated chromatin sample is more diluted, adjust the size of the tubes used for pre-clearing from 2 mL capacity to 5 mL. As 5 mL tubes do not fit the magnetic rack, holding the tube and the rack close together by hand is recommended

40. Add 1364 µL of the diluted chromatin and rotate (360° rotation, 40 rpm) for 30 min at room temperature.

*Note: 2%* of the actual volume of diluted chromatin is added to avoid repetition of the pre-clearing in case of loss during washing and pipetting.

#### S4.3.8. Immunoprecipitation

41. Touch spin the tubes containing beads coupled with antibodies from step 38, then place them in the magnetic rack for 2 min and discard the supernatant.
42. Touch spin the tube of cleaned beads with chromatin from step 40, and place it in the magnetic rack for 2 min and transfer the supernatant to a new 2 mL tube.

*Note*: The new 5 mL tube contains pre-cleared chromatin.

43. Separate 3.34 µL (50 ng) of pre-cleared chromatin as 'input control', add recovery buffer to a total volume of 82 µL and store at 4°C.
44. Place 333.34 µL of pre-cleared chromatin into each of the tubes with beads coupled with antibodies from step 38, then add 166.67 µL of complete IP buffer to each.
45. Rotate (360° rotation, 40 rpm) overnight (12-16 hr) at 4°C.

#### S4.3.9. Bead washing, decrosslinking and DNA recovery

46. Touch spin the tubes, place them on the magnetic rack for 2 min and discard supernatant.
47. Resuspend the beads in 200 µL of complete IP buffer.
48. Touch spin the tubes, place them on the magnetic rack for 5 min and discard supernatant.
49. Resuspend the beads in 200 µL of complete low salt wash buffer.
50. Touch spin the tubes, place them on the magnetic rack for 5 min and discard supernatant.
51. Repeat the washing with low salt wash buffer.
52. Resuspend the beads in 200 µL of complete high salt wash buffer.
53. Touch spin the tubes, place them on the magnetic rack for 5 min and discard supernatant.
54. Repeat the washing with high salt wash buffer.
55. Resuspend the beads in 57 µL of recovery buffer.
56. Elute the DNA from the beads in a Thermomixer at 1,000 rpm for 1h 30 min at 65ºC.
57. Touch spin the tubes and place them on the magnetic rack for 5 min.
58. Transfer the eluted chromatin to new clean LoBind 1.5 mL Eppendorf tubes.
59. Wash the beads with 30 µL of recovery buffer. Mix and touch spin the tubes.

60. Place the tubes on the magnetic rack. Transfer into the full volume of eluted chromatin from step 55 (i.e. 87 µL final).
61. Take out the ‘input control’ tube stored in the fridge.
62. Add 5 µL 20% SDS, 2 µL of RNase A, 5µL of proteinase K and 6 µL of NaCl to the ‘input control’ tube.
63. Add 2 µL of RNase A, 5 µL of proteinase K and 6 µL of NaCl to the tubes containing immunoprecipitated chromatin.
64. Incubate in a Thermomixer at 500 rpm for 1h 30 min at 68°C.
65. Purify the samples using MinElute PCR purification kit and elute in 32 µL of low TE buffer.

*Note*: DNA from the input and immunoprecipitated chromatin can be stored at - 20°C until library preparation.

66. Quantify samples with Qubit High Sensitivity kit as follows:

| Input | H3K4me1 | H3K4me3 | H3K27ac | H3K27me3 |
|---|---|---|---|---|
| 1 µl of 1/10 dilution | 6µl | 6µl | 2µl | 2µl |

*Note*: Different volumes of immunoprecipitated DNA are recommended for Qubit quantifications to ensure that the detected concentration is not below Qubit sensitivity limits.

67. Prepare the library using the Microplex Library Preparation Kit v3 following the manufacturer’s instructions.

*Note*: Recommended minimum input DNA 0.2 ng. It should be started from a volume of 10 µL.

| DNA input (ng) | Number of cycles |
|---|---|
| 50 | 5 |
| 20 | 6 |
| 10 | 7 |
| 5 | 8 |
| 3 | 9 |
| 2 | 10 |
| 1 | 11 |
| 0.5 | 12 |
| 0.3-0.2 | 13 |

#### S4.4. Modifications to BodyMap reference protocol

##### S4.4.1. Atlantic salmon brain

Follow the reference protocol, but use 50-75mg of initial tissue. Crosslink for 5 min. Centrifugate at 2,500g (step 11) and 1,500g (steps 14 and 17). Sonicate for 15 cycles (step 22).

##### S4.4.2. Immature gonads

Follow the reference protocol, but use 30-100mg of initial tissue. Sonicate for 18 cycles (step 22).

##### S4.4.3. Mature female gonads

Follow the reference protocol, but use 200mg of initial tissue. Sonicate for 18 cycles (step 22).

##### S4.4.4. Mature male gonads

More modifications were needed to perform ChIP from mature male gonads material due to sperm content. This protocol may need to be performed 2-4 times to ensure sufficient amount of generated chromatin from the gonad excluding contamination from sperm. This is important for successful immunoprecipitation. Please follow these instructions, in a similar way as for the ATAC-seq procedure;

*S4.3.4.1. Buffer recipes*

**Iodixanol, Gradient 1, Bottom layer solution**

- 40% (v/v) Iodixanol OptiPrep (Sigma-Aldrich, D1556)
- PBS Plus, 1x

*Note*: Prepare fresh and keep on ice

**Iodixanol, Gradient 1, Middle layer solution**

- 29% (v/v) Iodixanol OptiPrep (Sigma-Aldrich, D1556)
- PBS Plus, 1x
- 0.1% (v/v) Nonidet P40 Substitute (Roche, 11332473001)
- 0.15 mM Spermine (Sigma-Aldrich, S3256)
- 0.5 mM Spermidine (Sigma-Aldrich, S2626)

Note: *Prepare fresh and keep on ice*

**Iodixanol, Gradient 2 solution**

- 50% (v/v) Iodixanol OptiPrep (Sigma-Aldrich, D1556)
- PBS Plus, 1x

*Note*: Prepare fresh and keep on ice

*S4.3.4.2. Modifications for mature male gonad*

68. Transfer the tissue (400-600 mg) in a 5 mL tube containing 1 mL of PBS Plus buffer. Gently squeeze the tissue with a pestle against the tube to release the sperm, but do not disrupt the tissue into smaller pieces.
69. Vortex for at least 30 sec and filter through a 40 µm cell strainer suspended in a 50 mL Falcon tube. Add 500 µL of PBS Plus buffer over the cell strainer to rinse the tissue and place the tissue back into the 5 mL tube. Discard the filtrate containing mostly sperm.
70. Repeat step 1 and 2 to release sperm from the tissue as much as possible.
71. Transfer the tissue into an ice-cold dounce containing 5 mL cold PBS Plus and homogenize with pestle A and B.
72. Transfer the homogenate into a 5 mL Eppendorf tube. Avoid transferring pieces that did not effectively homogenize.
73. Centrifuge on a pre-cooled micro-centrifuge at 3,000g for 10 min at 4°C.
74. Discard the supernatant and add 800 µL of fresh PBS Plus buffer.
75. Add 560 µL 60% iodixanol and mix to obtain 1,400µL to achieve a solution containing 25% iodixanol.
76. Prepare a 5 mL tube for the first gradient centrifugation: place 1200 µL of bottom layer buffer (40% iodixanol) and gently add 1200 µL of middle layer buffer (29% iodixanol). Avoid mixing the layers.
77. Add solution from step 8 to the 5 mL tube over the middle layer solution.
78. Centrifuge in a pre-cooled swinging bucket centrifuge with brakes set to 'off' at 3,200g for 30 min at 4°C. This will result in a visible band between the 29% and 40% iodixanol layers.
79. Recover the nuclei band (200-400 µL) into a clean 1.5 mL Eppendorf tube. This can be done by careful removal of the first two layers (25% and 29% iodixanol) or by targeting the nuclei band.
80. Add and mix 0.7x volume of PBS Plus buffer to obtain a 15-20% iodixanol solution with nuclei.
81. Let the solution sediment for 5-10 min on ice.
82. Carefully transfer supernatant into another clean 1.5 mL Eppendorf tube. Avoid transferring pieces of tissue or big aggregates.
83. Prepare two 2 mL tubes for the second gradient centrifugation: place 300 µL of bottom layer buffer (50% iodixanol) and gently add to it half of the recovered nuclei solution from the previous step.
84. Centrifuge in a pre-cooled micro-centrifuge with brakes set to 'off' at 12,000g for 40 min at 4°C. A band should be visible between the 15-20% and 50% iodixanol layers.
85. Remove and discard the first gradient layer containing 15-20% iodixanol solution in both tubes.
86. Recover the nuclei layers and transfer them into a common 1.5 mL Eppendorf tube (100-200µL).
87. Dilute the nuclei layer 10 times with PBS plus to obtain a clean solution with nuclei.

88. Centrifuge at 5,000g for 10 min at 4°C.
89. Discard the supernatant and resuspend the nuclei pellet with 2.5 mL of PBS Plus buffer.
90. Take an aliquot of 10 µL from the suspension and mix with 10 µLof dye (Hoescht or Trypan Blue). Count the nuclei and sperm using hemocytometer. Gonad cell nuclei to sperm nuclei ratio between 1 to 2 and 1 to 5 is expected.
91. Add 67.5 µL of 37% formaldehyde solution at room temperature and mix by pipetting.

*Note*: When there are more than 24 million nuclei, add 2.43 mL more of PBS Plus buffer, then add 135 µL of 37% formaldehyde solution and split into two tubes.

92. Continue the protocol as stated in reference protocol (steps 9-20).
93. Resuspend the pellet and sonicate using the settings of the reference protocol as follows:

| Gonad cell nuclei to sperm nuclei ratio | Nuclei per 300µl sonication buffer | Sonication cycles |
|---|---|---|
| 1 to 5 | 1.5 million | 18 |
| 1 to 4 | 2 million | 18 |
| 1 to 3 | 2.5 million | 20 |
| 1 to 2 | 3 million | 20 |

94. Continue protocol as stated in reference protocol (step 23 onwards).

#### S4.4.5. Muscle

*S4.3.5.1 Buffer recipes*

**Hypotonic Buffer**

- 10 mM HEPES pH 7.4
- 10 mM KCl
- 5 mM $MgCl_2$
- 0.1% (v/v) Nonidet P40 Substitute (Roche, 11332473001)
- Water

*Note*: Store at 4°C temperature for up to 1 month. Complete the buffer for use with 1x PIC, 0.1 mM PMSF and 1.25 mM Sodium Butyrate (done under fume hood and keeping on ice)

*S4.3.5.2. Modifications for muscle*

95. Grind 1g of tissue using a mortar chilled on dry ice until it is powdery
96. Transfer the grinded material into a 5 mL tube containing 5 mL of cold complete hypotonic buffer.
97. Rotate (360° rotation, 40 rpm) for 10 min at 4°C.

98. Transfer to a douncer and homogenate using pestle A (approx. 5 strokes).

99. Transfer homogenate back to a 5 mL tube and add hypotonic buffer to make 5 mL volume if needed.

100. Take an aliquot of 10 µL from the suspension and mix with 10 µL of dye (Hoescht or Trypan Blue). Count nuclei using hemocytometer.

101. Add 135 µL of 37% formaldehyde solution at room temperature.

102. Rotate (360° rotation, 40 rpm) for 10min at 4°C

103. Quench the reaction with 720 µL of glycine 1M (0.125M final) and place it back in rotation for 10 min.

104. Transfer crosslinked tissue back to douncer and homogenize with pestle A (5 strokes).

105. Transfer the homogenate to a 5 mL tube.

106. Centrifuge at 2,500g for 10 min at 4°C.

107. Remove and eliminate the supernatant.

108. Resuspend with 5 mL of complete hypotonic buffer.

109. Transfer solution back to the douncer and homogenize with pestle A (5 strokes).

110. Filter using a 40 µm strainer placed over a 50 mL Falcon tube.

111. Transfer the filtrate into a new 5 mL tube.

112. Centrifuge at 1,000g for 10 min at 4°C.

113. Remove the supernatant.

114. Resuspend pellet in 4 mL of hypotonic buffer. Split the solution in order to obtain 3 million nuclei per 1.5mL Eppendorf tube.

115. Centrifuge at 3,000g for 10 min at 4°C.

116. Remove the supernatant.

*Note*: The nuclei pellet can be snap frozen and stored in -80ºC for up to 2 months.

117. Resuspend the pellet in a ratio of 300 µL of complete sonication buffer per 3 million of nuclei. Incubate on ice for 5min.

118. Sonicate the chromatin for 18 cycles, keeping the samples ice cold and using settings of the reference protocol (step 22).

119. Continue the protocol as stated in reference protocol from step 23 onwards.

**Text S5. Transcription factor binding site (TFBS) enrichment results**

Transcription factors (TF) are nuclear proteins that regulate transcription to modulate cell identity or tissue homeostasis by binding specific DNA sequence motifs, i.e., TF binding sites (TFBS). As explained in the main text, we used the robust open chromatin regions annotated as active promoters or enhancers by ChromHMM in TFBS enrichment analyses to seek evidence for enrichment of vertebrate TFs according to the known biology of embryogenesis (DevMaps) or adult tissue differentiation (BodyMaps). This was done separately for both salmonid species (Figs S12-S13), with the results compared to establish evidence in support of shared TFBS usage across species (Fig. S14). Though not exhaustive, examples of enriched TFBS are provided below, consistent with expected functions of vertebrate TFs, providing evidence for the high quality of our ChromHMM predictions.

#### S5.1. DevMap enriched TFBS

The Sox2::Pou5f1 complex is a critical regulator of pluripotency in vertebrates including zebrafish (*37*, *105*). In our study, the Sox2::Pou5f1 TFBS was most enriched at the late blastula and mid-gastrulation stages in active promoters and enhancers of both species (Fig. S12), with activity sharply declining from somitogenesis onwards. In zebrafish, Sox2 family and Pouf51 genes, alongside Nanog, are essential to initiate zygotic genome activation (ZGA) and the termination of the maternal programme that dominates before the maternal-to-zygotic transition (MZT) (*37*). The enrichment of this motif at these stages of salmonid embryogenesis suggests a conserved role in the MZT. Another TFBS showing the strongest enrichment in the late blastula, followed by mid-gastrulation, was MGA::EVX1, enriched in active promoters and enhancers in both salmonid species (Fig. S12). MGA is a bHLH TF that in zebrafish is highly expressed as a maternal transcript, before showing zygotic expression during gastrulation, and ongoing expression in developing brain, heart and gut during organogenesis, which is essential to the development of these organs (*106*). The enrichment of this TFBS in active regulatory elements at the earliest profiled stages of salmonid development is consistent with this TF being a maternal factor, with a role in early organogenesis.

In Atlantic salmon, we identified enrichment of the TFBS for Sox10 at mid-gastrulation and early somitogenesis in both active enhancers and promoters (Fig. S12). The same TFBS was most enriched at early somitogenesis in active enhancers of rainbow trout (Fig. S12). Sox10 plays a key role in the formation and differentiation of neural crest cell lineages including neural cells and melanocytes (*107*, *108*). The enrichment of Sox10 TFBS in our data is consistent with neural crest development being captured during salmonid embryogenesis. The TFBS for IRF6 similarly showed enrichment at mid-gastrulation of active promoters for both species. IRF6 is a maternal TF that causes severe gastrulation defects when ablated in *Xenopus* or zebrafish (*109*). TP63 binding sites were enriched in active regulatory elements of Atlantic salmon at early somitogenesis, which in zebrafish is a master regulator of ectoderm development, with ectoderm tissues specified during early somitogenesis (*110*). We also identified a strong enrichment of the TFBS for Hoxd10 specifically for active enhancers of both species at early somitogenesis, consistent with the established importance of Hox10 genes in the presomitic mesoderm (*111*) a structure that appears during the earliest stage of segmentation, leading into somitogenesis.

At later stages of embryogenesis, we identified enriched TFBSs consistent with the functions of conserved TFs. For example, the TFBS for MEF2B was enriched in both mid- to late-somitogenesis stage for both active enhancers and promoters of both species (Fig. S12). MEF2 family members are key regulators of cardiac and skeletal myogenic differentiation in mammals and teleosts (*112*, *113*), which are both highly active processes during the segmentation period. NRF1 (Nuclear respiratory factor 1) was an interesting TFBS enriched specifically in active promoters of both species at late somitogenesis (*114*), recognized as a master regulator of mitochondrial biogenesis, which may reflect the growing energy demands of late stage embryos, e.g. for coordinated muscle movement and brain activity.

#### S5.2. BodyMap enriched TFBS

Hepatocyte nuclear factors (HNFs) are a group of transcription factors with master functions in transcriptional regulation of liver-specific genes, promoting hepatocyte development (*115*). In our analysis, TFBS for two distinct HNFs, HNF1 and HNF4, were enriched in liver for both active enhancers and promoters of both species (across sexes and maturation stages) (Fig. S13). Both HNF1 and HNF4 play critical roles in maintaining hepatocyte function (*116*, *117*). HNF4 is a conserved nuclear receptor superfamily member indispensable for liver development in mammals (*117*). TFBS for IRF6 and TP63 were also enriched within active enhancer and promoters from liver samples of both species (Fig. S13). In mammals, IRF6 protects hepatocytes against liver steatosis by binding regulatory elements in peroxisome proliferator-activated receptor $\gamma$ (*118*). TP63 is a transcription factor that binds the promoter of *SIRT1* in mammals to regulate hepatic glucose production and insulin sensitivity (*119*).

In brain-active regulatory elements of both species, the most enriched TFBS, particularly in active enhancers, was Neurod1/2, representing master TFs for central nervous system and brain development (*120*). Specific to promoters of both species, the binding site for NRF1 was highly enriched in brain samples (Fig. S13). In mammals, NRF1 is expressed broadly across tissues and plays a key role in regulating mitochondrial biogenesis, respiration and cell redox homeostasis by binding the promoter of target genes (*121*, *122*). Hence, the enrichment of NRF1 binding sites in salmonid brains is logical, given the extensive metabolic demands of this tissue. Consistent with this, NRF1 binding sites were also highly enriched in regulatory elements activated during late stages of embryogenesis in both species (Fig. S12), where the brain is developing, and energy demands of the embryo are greatly increasing (e.g., according to our GO transcriptome analysis; Text S6). RFX2 was interesting an TFBS enriched in active brain enhancers and promoters in both salmonid species; this TF is considered essential for vertebrate ciliogenesis and is highly expressed in brain, where it regulates neuronal cilia differentiation (*123*). The enrichment of RFX2 binding sites in salmonid brain regulatory elements likely represents the proportionate greater importance of cilia differentiation in abundant brains neurons, compared to other tested tissues in the analysis (i.e., muscle, liver). Similar to NRF1, binding sites for RFX2 were also enriched in active enhancers of both salmonids at the latest stage of embryogenesis (Fig. S12), where brain and neural differentiation is highly active.

In muscle samples, we observed enrichment of binding sites for MEF2B in active regulatory elements, particularly in Atlantic salmon. As noted above, MEF2B binding sites were also enriched in active regulatory elements of both salmonid species at late stages of embryogenesis (Fig. S12), when muscle development is highly active. Six2 binding sites were enriched in active muscle regulatory elements of both species (Fig. S13). This TFBS was predicted to be an important regulator of skeletal muscle development in cattle, including for both myoblasts and differentiated muscle fibres (*124*).

#### S5.3 Overview of enriched TFBS shared between salmonid species

Overall, we identified 672 enriched TFBS across the four enrichment analyses performed, of which 58% were common between the two species (Fig. S14). We found more enriched TFBS shared across the two species in DevMap samples, 71% for active promoters and 56% for active promoters, which compared to 51% and 48% for the equivalent BodyMap regulatory elements, respectively. It was observed that rainbow trout had a larger proportion of unique rather than shared enriched TFBS compared to Atlantic salmon specifically for BodyMap samples (Fig. S14). Specifically, for rainbow trout BodyMap samples, 45% and 42% of active promoters and enhancers were not shared with Atlantic salmon, whereas the reciprocal values for Atlantic salmon were 12% and 27%, respectively (Fig. S14). Conversely, for DevMap samples, the number of species-specific enriched TFBS in both active enhancers and promoters was comparable in both species. These results may reflect the fact that the DevMap ATAC-seq and ChIP-seq experiments were performed at the same site, whereas for BodyMap, different collaborators performed these analyses independently, albeit with standardized consortium protocols.

### Text S6. Transcriptome dynamics during salmonid embryogenesis

#### S6.1. DevMap transcriptome dataset overview

For both Atlantic salmon and rainbow trout, our DevMap transcriptome datasets comprise 14 matched stages of embryogenesis, determined by relative age (*125*) and anatomical features observed during sampling. Below we report developmental processes captured across each major sampled stage based on the genes expressed in SOM clusters (Fig. 2A; Fig. S18-19), including GO enrichment analysis (Fig. S22-23; Tables S7-S8).

#### S6.2 Maternal to zygotic transition clusters

The earliest embryonic stage sampled was late cleavage, closely followed by early, mid and late blastulation stages. In Atlantic salmon (SOM clusters: 'Maternal 1', 'Maternal 2', 'Maternal 3' and 'Maternal 4') and rainbow trout (SOM clusters: 'Maternal 1', 'Maternal 2', 'Maternal 3'), SOM captured genes showing highest expression before or during blastulation, with a subsequent decrease in expression. This pattern is consistent with maternal transcripts, or a combination of maternal transcripts and genes expressed during early zygotic genome activation (ZGA). Atlantic salmon cluster 'Maternal 2' and rainbow trout cluster 'Maternal 1' were enriched for GO terms linked to cell division, i.e. '*cell cycle*', '*chromosome segregation*', '*centrosome cycle*', '*cytokinesis*' and '*DNA replication*', consistent with cell cleavages (Figs S22-23). These terms are also associated with housekeeping roles, which may explain why Atlantic salmon 'Maternal 2' cluster genes maintain some expression throughout embryogenesis (Fig. 2A; Fig. S18; Fig. S22). Atlantic salmon cluster 'Maternal 1' strongly resembled a maternal transcript degradation pattern, and was enriched for GO term GO:0048511 '*rhythmic process*', explained by genes including *nocta*, which contributes to 3'-5' exonuclease activity and RNA deadenylation (*126*) and *dbpa*, encoding an RNA helicase involved in RNA structural rearrangements (*127*), suggesting that maternal or early zygotic transcripts are performing roles related to RNA modification and translational regulation. Other terms present in Atlantic salmon cluster 'Maternal 1' , and rainbow trout clusters 'Maternal 1' and 'Maternal 2' included GO:0045892 '*negative regulation of transcription, DNA-templated*' and GO:0046578 '*regulation of Ras protein signal transduction*' (Figs S22-23); these terms are involved in cellular differentiation, suggesting the pre-blastulation stage represents a transition between cleavage and early blastulation, with maternal transcripts still present and the zygotic genome not yet fully activated. Other terms found in maternal SOM clusters, including GO:0051090 '*regulation of DNA-binding transcription factor activity*', GO:0006406 '*mRNA export from nucleus*' (rainbow trout 'Maternal 2') are however consistent with some degree of ZGA in late cleavage stages (Figs S22-23).

Atlantic salmon SOM clusters showing up-expression during blastulation include 'Maternal 4', capturing genes also highly expressed in pre-blastulation, and 'Blastula 1', a highly specific mid-blastulation cluster (Fig. 2A; Fig. S18). Atlantic salmon SOM cluster 'Blastula 2' spans the transition from late blastulation to early gastrulation. The most closely related rainbow trout SOM clusters showed less overlap between pre-blastulation and blastulation, with no obvious equivalent to Atlantic salmon ‘Maternal 4’ (Fig. 2A; Fig. S19). Rainbow trout SOM cluster ‘Blastula 1’ with mid-blastulation specific expression, showed no enriched GO terms, despite being comprised of 1,261 genes, potentially highlighting issues in the annotation of early development genes. Enriched GO terms for Atlantic salmon ‘Blastula 1’ included GO:0007399 ‘*nervous system* development’. However, the genes explaining this term were associated with endoderm development, such as *her5* (*128*) as well as genes associated with microtubule stabilization, such as *campsap2a* (*129*) (Fig. S22). Other terms enriched in both blastulation-specific clusters were GO:0043408 ‘regulation of *MAPK* cascade’, GO:0007049 ‘cell cycle’, GO:0030178 ‘negative regulation of *Wnt* signalling pathway’ and several terms associated with RNA modification and metabolism (Fig. S22).

#### 6.3. Gastrulation clusters

Atlantic salmon had two SOM clusters capturing genes showing up-regulation during gastrulation (Fig. 2A; Fig. S18). As noted above, Atlantic salmon cluster ‘Blastula 2’ span the transition from blastulation to gastrulation, whereas ‘Gastrula 1’ showed gastrulation-specific expression, with highest expression in the late gastrula stage. In rainbow trout (Fig. 2A; Fig. S19), cluster ‘Blastula’ spanned the transition from blastulation to gastrulation, while cluster ‘Gastrula 1’ had gastrulation-specific expression, with the highest expression found towards late gastrulation. Atlantic salmon SOM cluster ‘Gastrula 1’ was enriched for two main processes: GO:0072330 ‘monocarboxylic acid biosynthesis’ (explained by eight genes, including *acaca*, *elovl6* and *ptgs2v*), and GO:0042981 ‘regulation of apoptosis’ (explained by eighteen genes, including *dedd*, *mdm2*, *socs3* and *diabloa*) (Fig. S22). Several remaining terms below the employed significance threshold were associated with gastrulation, including GO:0007369 ‘Gastrulation’, GO:0060027 ‘Convergent extension during gastrulation’, GO:0098609 ‘cell-cell adhesion’, as well as terms associated with morphogenesis, cell differentiation and Wnt signalling. Gastrulation clusters in both species were highly enriched in terms associated with Wnt signalling (Fig. S22-23), explained by genes including *wnt8a*, *wnt3a*, *wnt10b* among others. Other enriched cell-cell signalling pathways were Jak/Stat (Atlantic salmon), as well as Hippo, Ras and Notch (both species). Additionally, both SOM clusters were enriched for terms associated with nervous system development, explained by genes of the Brinp, Gata and Fox families. Both SOM clusters were also enriched in terms associated with cell-cell adhesion, explained by genes of the cadherin superfamily among others (Figs S22-23).

#### S6.4. Somitogenesis clusters

For Atlantic salmon, SOM clusters ‘Somitogenesis 1’ and ‘Somitogenesis 2’ correspond to genes showing up-regulation during early-to-mid segmentation and mid-to-late segmentation, respectively (Fig. 2A; Fig. S18). Cluster ‘Pharyngula 1’ included genes that were relatively up-regulated during late segmentation, but showed highest expression in pharyngula stages. Cluster ‘Somitogenesis 1’ was enriched for GO:0016055 ‘Wnt signalling pathway’ (twenty-three genes), as well as GO:0048513 ‘animal organ development’ (thirty-seven genes) (Fig. S22). Enriched GO terms for this cluster are broadly linked with tissue development, organogenesis, cell signalling and, importantly, GO:0007389 ‘pattern specification process’. Cluster ‘Somitogenesis 2’ showed similar GO enrichment, including ‘Wnt signalling pathway’, but also terms associated with muscle development (Fig. S22). Additionally, GO:0001568 ‘blood vessel development’ was enriched in Atlantic salmon ‘Somitogenesis 1’. For rainbow trout, SOM clusters ‘Somitogenesis 1’ and ‘Somitogenesis 2’ captured genes showing highest expression during early-to-mid segmentation and mid-to-late segmentation respectively (Fig. 2A; Fig. S19), mirroring results in

Atlantic salmon. While the enriched GO terms in 'Somitogenesis 1' clusters were almost identical, rainbow trout cluster 'Somitogenesis 2' also showed enrichment for GO:0033014 'tetrapyrrole biosynthetic process', and terms associated with the circulatory system, including GO:0015671 'gas transport' and GO:0008015 'blood circulation' (Fig. S23). Despite this difference, other terms associated with morphogenesis and tissue development were shared between Atlantic salmon and rainbow trout.

#### S6.5. Pharyngula clusters

Genes comprising the Atlantic salmon SOM clusters 'Pharyngula 1' and 'Pharyngula 2' showed highest expression during the pharyngula period, with 'Pharyngula 1' having the highest normalized expression. A third cluster ('Early/late') had high expression in both late cleavage and pharyngula (Fig. 2A; Fig. S18). A similar pattern was observed for rainbow trout (Fig. 2A; Fig. S19), with cluster two pharyngula specific clusters and one Early/late cluster. Enriched GO terms for the pharyngula clusters reflect upregulation of processes associated with metabolism, such as ATP production, glucose, carbohydrate and fatty acid metabolism, consistent with increased energy production demands of the developing fish (Figs S22-23). Pharyngula-specific clusters were particularly enriched in metabolic processes in both species (Figs S22-23). Terms associated with brain function (e.g., synaptic signalling or regulation of neurotransmitter levels) and nervous system development were also prevalent in all pharyngula clusters. Moreover, all clusters were highly enriched for GO terms associated with development, especially organ and tissue development, including urogenital system development, renal development, eye development and epithelium development, among others (Figs S22-23).

#### S6.6. Constitutive embryonic gene expression

The largest SOM clusters in the Atlantic salmon and rainbow trout embryonic datasets were represented by genes with low normalized expression variability across developmental stages, capturing ~40% and 50% of all genes in Atlantic salmon and rainbow trout, respectively (Fig. 2A; Figs S18-19). Enriched GO terms for these clusters were composed almost exclusively of housekeeping functions linked to DNA repair, the cell cycle, nucleic acid metabolism, ribosome production and maintenance, among others. Although these constitutive SOM clusters did not display stage-specific expression, there were small variations in terms of expression pattern. For example, cluster 'Constitutive 1' in Atlantic salmon and cluster 'Constitutive 3' in rainbow trout both display an increase in expression towards the pharyngula period. Interestingly, the most highly enriched terms in these clusters were associated with cellular respiration (Atlantic salmon) and ATP metabolism (rainbow trout) (Fig. S22-23), mirroring the prevalence of metabolic genes in pharyngula clusters, consistent with the increased energy requirements in late embryonic stages, likely due to the demands of brain and muscle tissue, which together comprise a large fraction of the embryo mass.

### Text S7. Transcriptome dynamics across salmonid tissues at different maturation stages

#### S7.1. BodyMap transcriptome dataset overview

BodyMap datasets are composed of eight tissues in adult male and female Atlantic salmon and rainbow trout at two stages of ontogeny (sexually immature; sexually mature), with samples matched across species. SOM analysis captured a highly heterogeneous landscape of tissue expression, with most clusters displaying tissue-specific expression (23/36 in Atlantic salmon and 21/36 in rainbow trout) (Figs S25-S27). Moreover, 63% (29,787 out of 47,109) and 60% (23,037 out of 38,230) of Ensembl annotated coding genes were captured by tissue-specific

SOM clusters in Atlantic salmon and rainbow trout, respectively. An additional type of SOM cluster (hereafter 'multi-expression') was observed capturing genes showing up-regulation in more than one tissue. Below we explore the different SOM clusters based on tissue expression, along with enriched GO terms (Tables S7-8).

#### S7.2. Ovary clusters

In Atlantic salmon, three out of thirty-six SOM clusters showed highly ovary-specific expression (Figs S25-26). Combined, these clusters were composed of 5,024 genes, representing 10.6% of the tissue transcriptome. Rainbow trout similarly had three SOM clusters in this category (Figs S25-26), comprising 3,022 genes, or 7.9% of the tissue transcriptome. The most strongly enriched GO terms for both species were associated with egg production, including GO:0048477 'oogenesis' (Fig. S30-31), explained by genes including *bmp15*, which acts upstream of female sex differentiation and regulates oocyte maturation (*130*) and *buc*, which plays a role in the specification of the oocyte animal-vegetal axis, and is present within the blastomeres (*131*), suggesting maternally deposited genes are captured in these clusters. Other GO terms enriched in ovary clusters included GO:0051321 'meiotic cell cycle', explained by genes such as *rnf212*, a dosage sensitive ring finger protein with a role in crossing over during meiosis (*132*). Additional enriched GO terms included those involved in fatty acid metabolism, sperm-egg recognition and cytoskeleton organisation. While similar GO terms were enriched in ovary SOM clusters in both species (Figs S25-S27), rainbow trout showed more terms associated with non-coding RNA processing and RNA methylation, as well as embryonic development, including GO:0007369 'gastrulation' (Fig. S31), which included genes such as *bbs12*, *mkks* and *desi2*, which are each involved in convergent extension during gastrulation, as well as *msgn1*, involved in mesoderm differentiation in the transition from gastrulation to somitogenesis (*133*).

#### S7.3. Brain clusters

In Atlantic salmon and rainbow trout four and three respective SOM clusters captured brain-specific genes, which together represent 14% and 15% of all expressed genes in the tissue transcriptome (Figs S25-S27), respectively. Most enriched GO terms in these clusters (Fig. S29-30) were associated with brain and nervous system functions in both species, including GO:0007399 'nervous system development', GO:0099536 'synaptic signalling' and GO:0006813 'potassium ion transport'. Additionally, GO terms such as GO:0008015 'blood circulation' and GO:0006939 'smooth muscle contraction', among others, highlight the role of vascularisation and oxygenation for brain function.

#### S7.4. Testes clusters

In Atlantic salmon, four SOM clusters composed of 3,910 genes (6.5% of the tissue transcriptome) capture testes-specific gene expression (Figs S25-S26). These clusters could be demarcated into immature and mature testes specific gene expression, the strongest observed effect of maturity across all tissues sampled. The rainbow trout tissue panel dataset was missing mature testes samples due to sampling, so the same effect cannot be compared across species. The missing mature testis samples also explain the lower number (two) of testes-specific SOM clusters in rainbow trout, which captured 1,593 genes (4% of the tissue transcriptome). Enriched GO terms for rainbow trout SOM clusters showing testes-specific expression were associated with meiosis and cell division (Fig. S31), highlighting the high rate of cell division in testes. This included GO terms such as GO:0051321 'meiotic cell cycle', with most other enriched terms associated with DNA repair and rRNA production. Other enriched terms were associated with sperm production and function, such as GO:0007283 'spermatogenesis' and GO:0001539 ‘cilium or flagellum-dependent cell motility’. Finally, terms such as GO:0034470 ‘ncRNA processing’ suggest a role of ncRNA in sperm production. Enriched GO terms in Atlantic salmon testes-specific SOM clusters were very similar to rainbow trout (Figs S30-31), with notable differences linked to terms associated with development, including embryo, head, brain, neural tube, kidney and eye development, as well as pathways commonly associated with early development, such as Notch and Wnt.

#### S7.5. Muscle clusters

In Atlantic salmon, three SOM clusters captured muscle-specific genes (Figs S25-S26), composed of 3,183 genes (7% of the tissue transcriptome). This includes a small (292 genes) cluster expressed almost exclusively in immature females. Enriched GO terms for these clusters (Fig. S30) are exclusively associated with muscle structural proteins, including GO:0061061 ‘muscle contraction’, GO:0051146 ‘striated muscle cell differentiation’, GO:0048741 ‘skeletal muscle fibre development’ and GO:0006936 ‘muscle contraction’. The other enriched GO terms were largely associated with metabolism and energy production/storage, including GO:0006757 ‘ATP generation’, GO:0005977 ‘glycogen metabolic process’, GO:0006603 ‘phosphocreatine metabolic process’, GO:0005975 ‘carbohydrate metabolic process’ and GO:0006112 ‘energy reserve metabolic process’. Enriched GO terms in a female muscle specific cluster for Atlantic salmon were mainly linked to metabolism, including triglyceride, carbohydrate, fructose and proline and folic-acid metabolism, but also the cellular response to insulin (Fig. S30).

The four rainbow trout muscle-specific SOM clusters (Fig. S25, Fig. S27) captured 3,486 genes (9% of the tissue transcriptome) were strongly conserved with Atlantic salmon, including a small immature female-specific cluster. Enriched GO terms extensively overlapped for muscle-specific SOM clusters across species (Figs S30-31), with the exception that rainbow trout clusters were specifically enriched for embryonic development associated terms. The rainbow trout immature female-specific muscle SOM cluster comprised 494 genes and enriched for GO terms associated with metabolism. Other enriched rainbow trout GO terms not present in Atlantic salmon included T-cell activation, endocytosis and adaptive immune response.

#### S7.6. Liver clusters

In Atlantic salmon, three liver-specific SOM clusters captured 4,057 genes (8.5% of the tissue transcriptome) (Figs S25-S26). One of these clusters captured 314 genes specifically up-regulated in mature males. Rainbow trout had four liver-specific SOM clusters (Fig. S25, Fig. S27), which displayed higher heterogeneity, including clusters showing up-regulation in mature males, mature females, immature fish of both sexes, and one showing more homogeneous up-regulation. Enriched GO terms in the liver clusters for both species included diverse terms associated with metabolism, highlighting the important metabolic role played by the liver (Fig. S30-31; Tables S7-S8). Enriched GO terms associated with hemostasis in both species highlight the liver’s role in coagulation (Figs S30-31). Liver immune functions are represented by enriched GO terms in both species (Figs S30-31), including GO:0006956 ‘complement activation’ and GO:0002449 ‘lymphocyte mediated immunity’

#### S7.7. Head kidney clusters

In Atlantic salmon, two SOM clusters were head kidney specific (Figs S25-S26), capturing 2,261 genes (4.7% of the tissue transcriptome). Rainbow trout also had two head kidney specific SOM clusters comprised of 1,783 genes (4.6% of the tissue transcriptome) (Fig. S25, Fig. S27). In both species, enriched GO terms (Fig. S29-S30; Table S6-S7) were consistent with the head kidney’s primary role in immunity, for example, GO:0050776 ‘regulation of immune response’, GO:0006935 ‘chemotaxis’, GO:0019221 ‘cytokine mediated signalling pathway’, GO:0050863 ‘regulation of T-cell activation’, GO:0002250 ‘adaptive immune response’ and GO:0002250 ‘phagocytosis’, reflecting both primary and secondary immune functions. The primary hematopoietic function of the head kidney is reflected in the enriched term GO:0060218 ‘hematopoietic stem cell differentiation’.

**S7.8. Gill clusters**

In Atlantic salmon, two SOM clusters showed gill-specific expression (Figs S25-S26), capturing 3,169 genes (6.7% of the tissue transcriptome). Rainbow trout similarly had two gill-specific SOM clusters (Fig. 2; Fig. S26), capturing 1,662 genes (4.3% the tissue transcriptome). In both species, enriched GO terms for these gill clusters (Fig. S30-S31; Table S7-S8) captured many immune terms, including GO:0006955 ‘immune response’, GO:0060326 ‘cell chemotaxis’, GO:0019882 ‘antigen processing and presentation’ and GO:0031640 ‘killing of cells of other organism’, highlighting the gill’s key immune functions. Enriched GO terms associated with waste excretion and homeostatic function were also shared across both species (Fig. S30-S31; Table S7-S8), including GO:0072488 ‘ammonium transmembrane transport’ and GO:0098655 ‘cation transmembrane transport’. Other enriched GO terms associated with development and regulation of the circulatory system reflect the highly vascularized nature of the gill as the key respiration site.

**S7.9. Distal intestine clusters**

In Atlantic salmon and rainbow trout, two 2 SOM clusters showed distal intestine-specific expression, comprising 2,249 and 2,079 genes, respectively (Figs S25-S27). One notable species difference was evidence for higher gill expression heterogeneity in rainbow trout, with one SOM cluster presenting lower expression in mature females, and the other presenting lower expression in mature males, while remaining at constant levels in immature individuals of both sexes (Figs S25-S27). This observation is consistent with sexual dimorphism in intestine physiology linked to sexual maturation in rainbow trout. Enriched GO terms for salmonid gill SOM clusters captured the expected physiological roles of the distal intestine, including metabolism, ion transport and extracellular matrix organisation, as well as terms associated with muscle and circulatory system function.

**S7.10. Constitutive clusters**

Several SOM clusters captured genes showing homogenous expression across tissues, sexes and maturation stages in both species. In Atlantic salmon, four such ‘constitutive’ SOM clusters captured 9,458 genes (20% of the tissue transcriptome) (Fig. S26). In rainbow trout, three constitutive SOM clusters captured 6,179 genes (16% of the tissue transcriptome) were captured (Fig. S27). Enriched GO terms in the constitutive SOM clusters for both species largely represent housekeeping functions shared across cells types, including translation, histone modification, protein modification and protein catabolism, ncRNA metabolism, DNA repair and biosynthesis, ribosome biogenesis, ATP biosynthesis, and GO terms associated with mRNA processing, modification and transport, along with other functions associated with cell viability (Tables S7-8).

**S7.11. Multi-expression clusters**

For both the Atlantic salmon and rainbow trout datasets, an interesting group of SOM clusters showed upregulation in more than one tissue, potentially highlighting shared biological processes among different groups of tissues. In Atlantic salmon, nine multi-expression SOM clusters were captured (Fig. S26). Four were up-regulated in ovary and either brain, muscle, liver or head kidney. Three further Atlantic salmon SOM clusters showed up-regulation in brain and one each of testes, muscle or distal intestine. The remaining two multi-expression SOM clusters were up-regulated specifically in head-kidney and liver, or specifically in head-kidney and gill. Eleven rainbow trout multi-expression SOM clusters were observed (Fig. S27). Four showed upregulation in ovary and either brain, testes, muscle or head kidney, with an ovary plus head-kidney cluster also showing up-regulation in testes. Three multi-expression clusters were expressed in brain and either testes, muscle or head kidney. The remaining multi-expression clusters were gill plus intestine, muscle plus liver, muscle plus liver plus head kidney, and head kidney plus gill/intestine.

**Text S8. Supplementary text on regulatory element evolution (supporting Fig. 4)**

72.8% of active promoters were found in Early regions, with 48.5%, 22.8% and 28.8% belonging to the Shared, Alignable only, and Exclusive categories, respectively (Table S11). A similar proportion of active enhancers (74.8%) were located in Early regions, with 25.9%, 46.2% and 27.8% belonging to the same respective categories. A larger proportion of active promoters (67.0%, $p < 0.0001$, Fisher's Exact Test) and enhancers (55.1%, $p < 0.00001$, Fisher's Exact Test) belonged to the Shared category in Late compared to Early regions (Table S11). Reciprocally, fewer active promoters (14.4%, $p < 0.0001$, Fisher's Exact Test) and enhancers (8.34%, $p < 0.0001$, Fisher's Exact Test) belonged to the Exclusive category in Late regions. For active promoters and enhancers, 22.8% vs. 18.6% ($p < 0.0001$, Fisher's Exact Test) and 46.2 % vs. 36.6% ($p = <0.0001$, Fisher's Exact Test) belonged to the Alignable only category in Early vs. Late regions, respectively (Table S11). While a smaller proportion of active promoters and enhancers were located in Late regions, they show increased density compared to Early regions (one promoter every 74.5 kb vs. 116 kb, and one enhancer every 57.9 kb vs. 80.1 kb, respectively) when normalised to the size of each region.
