## Supplementary Data for "Salmonids reveal principles of regulatory evolution following autotetraploidization"

**Supplementary Datasets:**

*Available through embedded hyperlinks*

[**Supplementary Dataset 1**.](https://drive.google.com/file/d/1XB0s-YK-IFhQZpyzP6sE7td2g-4EaMLe/view) Genomic coordinates of 225,801 reference robust open chromatin regions in the Atlantic salmon genome, including coordinates of their syntenic/orthologous regions within the Atlantic salmon, rainbow trout and northern pike genomes, based on whole genome alignment.

[**Supplementary Dataset 2**](https://docs.google.com/spreadsheets/d/19SfbN0PXo3YTmqBGFns1YTsUZq7CSXdg/edit?usp=drive_link&ouid=117876630598606092083&rtpof=true&sd=true). Final dataset of active promoter and enhancer elements belonging to the three Alignment categories (‘Shared’, ‘Alignable only’ and ‘Exclusive only’)

[**Supplementary Dataset 3**](https://drive.google.com/file/d/1feh8sxGzrrjpJvoD5NRiPKxKXMVLwAhe/view?usp=drive_link). All conserved CNEs defined in the northern pike genome, including their estimated phylogenetic age.

[**Supplementary Dataset 4**](https://drive.google.com/file/d/1bR2N8sNaZNVBdEzMhVDvOyURYT11r9WG/view). Atlantic salmon enhancer-CNEs from the three alignment categories: ‘Shared’, ‘Alignable only’ and ‘Exclusive’.

[**Supplementary Dataset 5**](https://docs.google.com/spreadsheets/d/1MG-X3TjbryDCPCRGK15RQKjXdkHYx9z5/edit?gid=1531917611#gid=1531917611). Atlantic salmon enhancer-CNEs counts and length information split per: sample-types), ii) alignment category (Shared, Alignable only, Exclusive) and iii) rediploidization region (Early and Late).
